# Phosphorylation tunes electrostatically driven protein-RNA interactions

**DOI:** 10.64898/2026.04.27.720971

**Authors:** Marie Synakewicz, Adhvitha Premanand, Héloïse Bürgisser, Sarah Habeler, Lucia R. Franchini, Noémie Kociolek, Mark Nüesch, Antoine Cléry, Daniel Nettels, Frédéric Allain, Nina Hartrampf, Benjamin Schuler

**Affiliations:** Department of Biochemistry, University of Zurich, Switzerland; Department of Chemistry, University of Zurich, Switzerland; Department of Biology, Institute of Biochemistry, ETH Zurich, Switzerland; Department of Physics, University of Zurich, Switzerland

**Keywords:** RNA binding, SR protein, phosphorylation, phosphomimetic, automated fast-flow peptide synthesis (AFPS), single-molecule spectroscopy, Förster resonance energy transfer (FRET)

## Abstract

Phosphorylation of intrinsically disordered proteins (IDPs) is essential for regulating biomolecular interactions in many cellular processes. However, a quantitative understanding of how phosphorylation tunes the affinity between highly charged IDPs and nucleic acids is lacking. Here, we show that multi-site phosphorylation of the disordered arginine/serine-rich (RS) domain of the splicing factor SRSF1 acts as an electrostatic rheostat that governs RNA binding. By combining enzymatic phosphorylation, phosphomimetic variants, and chemically synthesised phosphopeptides with single-molecule Förster resonance energy transfer measurements, we reveal the RS domain to be a potent driver of protein–RNA association. Increasing phosphorylation progressively reduces this interaction, and extensive phosphorylation eliminates detectable RNA binding. Remarkably, the binding free energy depends linearly on RS-domain net charge, regardless of whether the charge arises from phosphorylation or acidic residues introduced as phosphomimetics. Together, our findings uncover a quantitative framework for how phosphorylation tunes the interactions of charged IDPs and rationalize why two acidic residues are required to mimic a single phosphorylation event.

## INTRODUCTION

Cellular communication and homeostasis are often regulated by post-translational modifications [1]. An important example is phosphorylation, the effects of which range from the modulation of enzymatic activity through conformational changes triggered by highly specific single-site phosphorylation to an overall influence on the electrostatic interactions between charged biomolecules. Examples include the binding interactions between proteins, especially for intrinsically disordered proteins (IDPs)[2, 3], which are particularly prone to post-translational modification[4]. Many IDPs are highly charged, especially those involved in binding nucleic acids[5], and we thus expect an important role of phosphorylation on such interactions[6]. However, a quantitative understanding of the influence of phosphorylation on the affinity between IDPs and nucleic acids is currently lacking.

Here, we address this question for an important example of an RNA-binding protein, the highly positively charged serine/arginine-rich splicing factor 1 (SRSF1). As a member of the Ser-Arg (SR) protein family, it regulates exon inclusion in the constitutive and alternative splicing of pre-messenger RNAs (pre-mRNAs), as well as other crucial biological functions, such as nuclear export of RNAs, translation, mRNA homeostasis, and mRNA decay [7–9]. Notably, the function and sub-cellular localization of SRSF1 are regulated by massive phosphorylation of its RS domain, a disordered Arg-Ser-rich region. Previous work has qualitatively shown an influence of phosphorylation on RNA binding [10]. To be able to quantitatively investigate the role of phosphorylation and charge on the interactions between SRSF1 — and especially the RS domain — and RNA, we use single-molecule Förster resonance energy transfer (FRET) spectroscopy combined with enzymatic phosphorylation, site-specific phosphoserine incorporation by chemical synthesis, and glutamate-based phosphomimetics. By varying the phosphorylation state and the net charge over a broad range, the affinity of the RS domain for RNA can be tuned across several orders of magnitude, ranging from nanomolar affinity to complete loss of binding. Phosphorylation of the RS domain is thus a major determinant of the interactions between SRSF1 and RNA, and its net charge is related roughly linearly with the binding free energy.

## RESULTS

SRSF1 comprises two structured RNA recognition motifs (RRMs) and a C-terminal intrinsically disordered RS domain (Figure 1a) [11–16]. The RRMs interact specifically with the bases of the target sequences in single-stranded RNA (ssRNA), and the positively charged RS domain is thought to interact largely non-specifically with RNA [15–18]. The RS domain contains 19 Arg residues and 20 Ser residues, and all of the latter can be phosphorylated [19–22]. To examine the role of phosphorylation in the SRSF1-RNA interactions, we first dissected the contributions of the RRMs and the RS domain to RNA binding by producing different SRSF1 constructs and measuring their affinity to RNA. In a second step, we used enzymatic phosphorylation of recombinantly expressed RS domain as well as the incorporation of site-specific phosphoserine residues by automated fast-flow peptide synthesis (AFPS) to assess the RNA affinity as a function of the number and position of phosphate groups. We then compared their RNA affinity with those of recombinantly expressed phosphomimetic variants. Throughout, single-molecule FRET experiments enabled us to probe biomolecular interactions at very low protein and RNA concentrations, which minimizes many of the notorious complications arising from protein aggregation and (micro-)phase separation in systems such as SRSF1 [23, 24].

**FIGURE 1.**
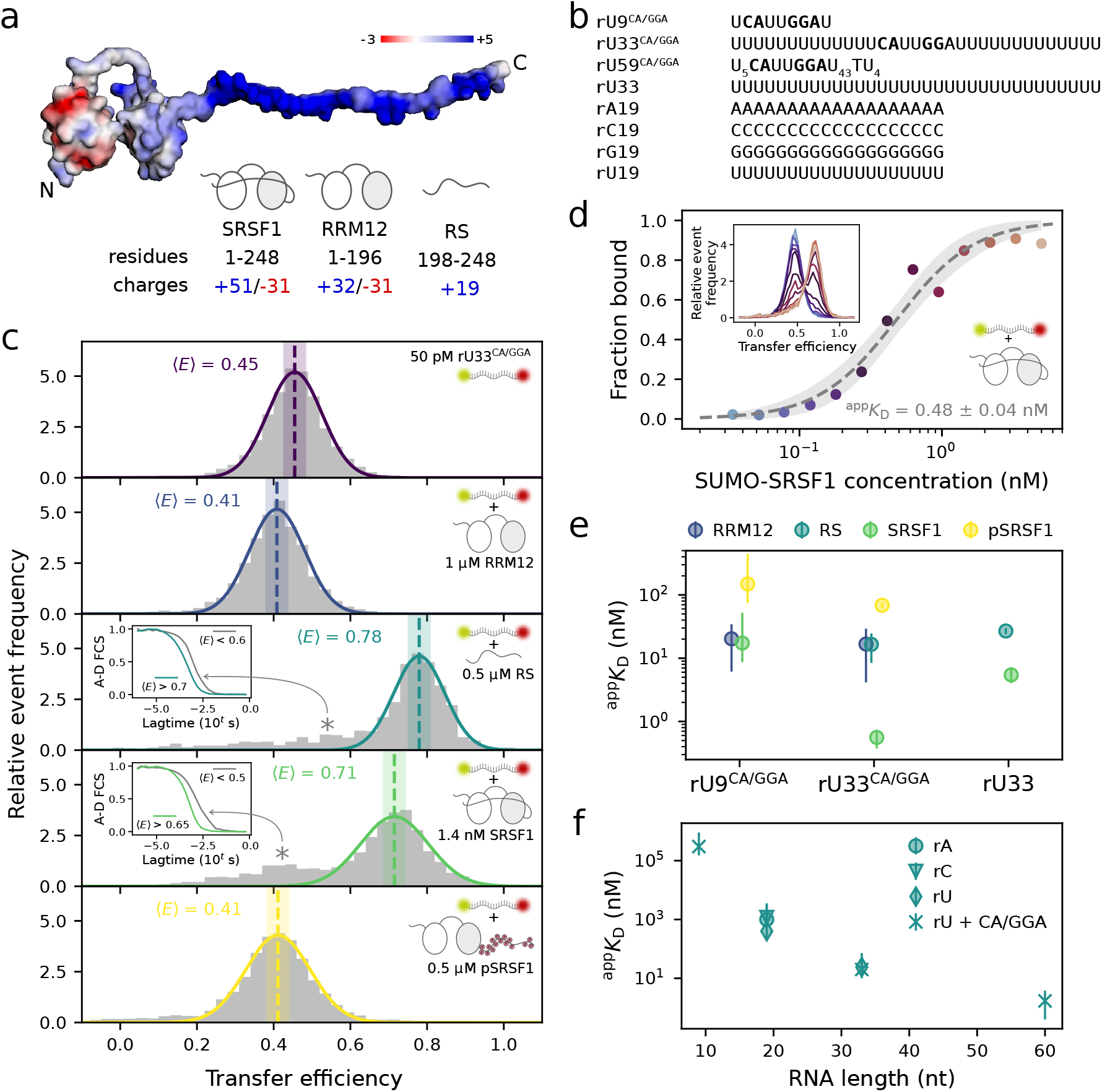
Quantifying SRSF1-RNA interactions with single-molecule spectroscopy. (a) Model of SRSF1 based on a structure generated by AlphaFold[25] (Q07955-F1), with the electrostatic surface potential colored in units of *k*_B_*T* per unit charge[26]. For each protein construct indicated by the cartoons, the residue boundaries and the total number of positive and negatively charged amino acids are given. (b) Nucleotide sequences (5^*′*^ → 3^*′*^) of single-stranded RNAs used to probe protein-RNA interactions. The binding motifs for RRM1 (CA) and RRM2 (GGA) are highlighted in bold. Terminal and internal modifications used for labeling are omitted for clarity but are shown in Table S1. (c) Representative normalised transfer efficiency histograms showing how protein binding affects the compaction of RNA. The mean of the Gaussian peak fit is indicated by a dashed line, the shaded band represents an uncertainty of 0.03 [27]. Asterisks indicate populations arising from large oligomeric species or aggregates, as highlighted by subpopulation-specific FCS in the corresponding insets: the diffusion time of the 1:1 bound population (solid lines in respective color) is of the same timescale as that of the free RNA (see Figure S1), whereas the diffusion times of the large oligomers or aggregates are much longer (solid lines in gray). (d) Representative titration of donor/acceptor-labeled rU33^CA/GGA^ with SUMO-SRSF1. The inset shows an overlay of the normalised transfer efficiency histograms of individual measurements, color-coded according to protein concentration. The fraction bound is fitted with Equation (2) (gray dashed line with shaded area corresponding to 95% confidence intervals) to obtain the apparent equilibrium dissociation constant, ^app^*K*_D_ (fit value and error of the fit shown). For more details on data analysis, see Figure S2 and Methods. (e) Comparison of ^app^*K*_D_ for the different protein variants and RNAs. Binding of RRM12 to rU33 could not be detected at RRM12 concentrations of up to 5 µM. The onset of the interaction of the RS domain with rU9^CA/GGA^ could be observed, but only a rough estimate of the affinity could be obtained (Figure S4). Means represent values obtained for both SUMO-tagged and untagged proteins, error bars represent the standard deviation (*N* > 1 measurements) or an uncertainty of a factor of 2 (*N* = 1 measurements)[28]. See Table S2 for details of individual protein constructs. (f) ^app^*K*_D_ of the untagged RS domain binding to ssRNA as a function of RNA length. The value for 9 nt is an estimate based on the binding response observed up to 30 µM RS domain (Figure S4). Error bars represent standard deviations (*N* > 1 measurements), an uncertainty of a factor of 2 (*N* = 1 measurements)[28], or a value derived from bootstrapping (rU59^CA/GGA^). See Table S2 and Methods for details. All affinities were measured at an ionic strength of 0.2 M.

We thus recombinantly expressed full-length SRSF1 (containing the Y37S and Y72S substitutions to increase solubility without affecting RNA binding [16]); a construct containing only the N-terminal RRMs (here termed RRM12); and the RS domain (Figure 1a). Since neither of the RRMs interacts specifically with uracil, we designed ssRNA sequences of different lengths that contain only one target sequence for each RRM, CA for RRM1, and GGA for RRM2 [15, 16], flanked by stretches of uracil (rU9^CA/GGA^, rU33^CA/GGA^ and rU59^CA/GGA^, see Figure 1b and Table S1). To distinguish sequence- and nucleobase-specific from purely electrostatically driven interactions, we furthermore included homopolymeric ssRNAs without these binding motifs (rA19, rC19, rG19, rU19 and rU33, see Table S1) in our measurements. We site-specifically labeled the 5’ and 3’ ends of the RNAs with acceptor and donor fluorophores, respectively, to study the protein-RNA interactions using single-molecule FRET. This experimental design allows us to obtain binding affinities at the same time as information on intramolecular distances and distance changes in the ssRNA conformational ensemble.

### RRMs and RS domain display different binding characteristics but similar affinities

To quantify the affinity of the different SRSF1 constructs to RNA and to elucidate their mode of interaction, we employ the changes in FRET efficiency that occur upon binding of the labeled RNA. Remarkably, RRM12 and the RS domain have very different effects on the RNA. Binding of the RRM12 construct to the rU33^CA/GGA^ RNA causes a shift in mean FRET efficiency from ⟨*E*⟩ ≈ 0.46 to ⟨*E*⟩ ≈ 0.41, corresponding to a slight extension of the RNA upon binding (Figure 1c). This observation suggests that binding of the two folded domains reduces the flexibility of the RNA segment containing the target sequence, resulting in an increase of the average end-to-end distance of the oligonucleotide chain. In contrast, binding of the RS domain causes a shift from ⟨*E*⟩ ≈ 0.46 to ⟨*E*⟩ ≈ 0.78, indicating a pronounced compaction of the RNA (Figure 1c). Similar to previous observations on other systems [29, 30], it is the flexibility of both the disordered RS domain and the ssRNA that enables this compaction upon mutual charge screening between the two oppositely charged polyelectrolytes. In the binding of full-length SRSF1, the combined effects of the RRMs and the RS domain cause a shift from ⟨*E*⟩ ≈ 0.46 to ⟨*E*⟩ ≈ 0.71 (Figure 1c). Presumably, binding of the RRMs constrains the conformational ensemble of RS domain-RNA interactions within the context of the full-length protein, leading to a slightly less pronounced overall compaction than for the isolated RS domain. Altogether, these results illustrate the different effects of SRSF1 and its component domains on its target RNA.

In a next step, the changes in FRET efficiency we observe can be employed to measure binding affinities. It is worth emphasizing that for a system such as SRSF1, which is prone to phase separation and aggregation even at relatively low concentrations [23, 24, 31], single-molecule spectroscopy provides a powerful way of identifying the presence of larger species in solution that interfere with the measurements. Moreover, subpopulation-specific FRET and fluorescence correlation spectroscopy (FCS) enable us to restrict the analysis to the species of interest (Figure S1), thus minimizing contributions from high-molecular-weight oligomers and aggregates that would interfere with the signal in ensemble measurements[32]. Based on these findings, we established an analysis that allowed us to to quantify RNA binding for all the constructs under study (Figure S2). Nevertheless, the low solubility of full-length SRSF1 required us to use a SUMO-tagged variant to obtain full titration curves (Figure 1d). However, a systematic comparison shows that the tag has little effect on RNA affinity (Figure S3).

To obtain information on the affinity and specificity of the binding of SRSF1 and its domains to RNA, we titrated different lengths and sequences of donor/acceptor-labeled ssRNAs with RRM12, the RS domain, and full-length SRSF1 (Figure 1e, Table S2). The apparent dissociation constant, ^app^*K*_D_, of RRM12 to RNA containing the binding motifs was 20 ± 10 nM for both rU9^CA/GGA^ and rU33^CA/GGA^ (Figure 1e). We could not detect binding of RRM12 to rU33, which lacks the binding motif, in the accessible protein concentration range of up to 5 µM, suggesting that any non-sequence-specific binding is at least three orders of magnitude weaker than specific binding. In contrast, the RS domain lacks sequence specificity and binds RNAs with and without the RRM binding motifs with similarly high affinity of 17 ± 8 nM (rU33^CA/GGA^) and 27 ± 3 nM (rU33), respectively (Figure 1d). For the interaction between the RS domain and rU9^CA/GGA^, we can only estimate ^app^*K*_D_ to be ∼300 µM, since the formation of large oligomers or aggregates at higher concentrations prevented measurements of a complete titration curve (Figure S4). Furthermore, the RS domain-RNA affinity increases by several orders of magnitude as the length of the RNA is increased from 9 to 60 nucleotides (Figure 1f), consistent with previous studies on protein-nucleic acid interactions that show higher affinity with greater length and net charge of the nucleic acid due to the larger number of possible binding configurations [33, 34]. Using a series of homopolymeric single-stranded RNAs, rA_19_, rC_19_, and rU_19_ [35], we find that the affinity of the RS domain is similar for all nucleobases (see Figure S5 for more detail) – especially, when compared with the strong effect of RNA length (Figure 1e). It was not feasible to quantify the affinity to rG_19_ owing to pronounced aggregation in the presence of RS domain, which occurred over the same range of protein concentrations where the other homopolymeric RNAs showed binding (Figure S5). Altogether, we have no indications that the RS domain-RNA interactions are specific for a particular nucleobase; rather, they are dominated by charge interactions.

In full-length SRSF1, the combined sequence-specific interactions of the folded RRMs for the binding motif and the electrostatic interactions of the disordered RS domain with RNA increase the total affinity to rU33^CA/GGA^ by almost two orders of magnitude compared to the RRMs alone, to ^app^*K*_D_ =0.6 ± 0.2 nM (Figure 1e). On the much shorter rU9^CA/GGA^ RNA, full-length SRSF1 exhibits a similar affinity as the RRM12 construct, suggesting that interactions of the RS domain are blocked by the footprint of the RRMs. Likewise, if the RRM binding motifs are removed, as in the rU33 RNA, SRSF1 exhibits an affinity closer to that of the isolated RS domain (Figure 1e). Together, these observations suggest that the binding of SRSF1 to RNA results both from sequence-specific interaction of the RRMs with the target sequences and from non-sequence-specific interaction of the disordered RS domain driven by its charges opposite to those of the phosphate groups on the RNA.

### Enzymatic phosphorylation by SRPK1 abolishes RS domain-RNA binding

Depending on the cellular localization and function of SRSF1, the RS domain can be heavily phosphorylated [36], raising the question of how these modifications affect RNA affinity. If the interaction between the RS domain and RNA is predominantly charge-driven, introducing negative charges by phosphorylation should weaken binding. In the cell, SRSF1 is phosphorylated by two kinases: in a first step, the N-terminal serines of the RS domain are phosphorylated processively by SRPK1, starting from the centre of the RS domain [37, 38]. After transport into the nucleus, SRSF1 is fully phosphorylated by Clk1[20–22]. Here, we used a three-pronged approach for assessing how different degrees of phosphorylation affect RNA binding, by combining enzymatic phosphorylation, chemical synthesis, and phosphomimetics. Enzymatically, we can introduce a large number of phosphates; by chemical synthesis, we can introduce a few phosphates at specific locations; and phosphomimetic amino acid substitutions allow us to introduce acidic amino acids at arbitrary positions and assess their effect compared to phosphorylated residues.

Using SRPK1, we were able to obtain phosphorylated SRSF1 and RS domain with an average of 16 added phosphates (Figure S6), which we refer to as pSRSF1 and RS-16xpS, respectively. With a charge of -2 per phosphate at pH 8 based on the relevant phosphoserine side chain *pK*_*a*_ of ∼ 5.6[39], phosphorylation thus inverts the net charge of SRSF1 and the RS domain, resulting in values of -12 and -13 for pSRSF1 and RS-16xpS, respectively. Owing to the repetitive nature of the RS domain, we were not able to identify the exact positions of the modifications, but given the processivity of SRPK1 [37, 38], the first 13 and final three serine residues are most likely to be phosphorylated. Transfer efficiency histograms of rU33^CA/GGA^ RNA bound to pSRSF1 no longer indicate compaction but instead a slight expansion, similar to the behavior observed with RRM12 (Figure 1c). We did not observe binding of RS-16xpS to rU33^CA/GGA^ RNA up to protein concentrations of 30 µM (Figure S7), indicating that SRPK1-mediated phosphorylation of the RS domain abolishes its affinity to RNA. However, we found a slightly but reproducibly lower affinity of pSRSF1 for RNAs containing the binding motif compared to RRM12 (Figure 1e). Since the affinities of pSRSF1 and RRM12 are similar for rU9^CA/GGA^ and rU33^CA/GGA^, electrostatic repulsion between RNA and the phosphorylated RS domain is unlikely to be the source of this decrease in affinity. Recently, Zhang and co-workers reported a similar trend and suggested that the phosphorylated RS domain interacts with arginine residues close to the RNA binding interfaces of the RRMs and in the inter-RRM linker, thereby reducing the overall affinity to RNA [40].

### Titrating the net charge of the RS domain

Enzymatic phosphorylation shows that charge inversion of the RS domain prevents it from binding to RNA. Addressing the effect of phosphorylation on the RS domain affinity for RNA more quantitatively thus requires us to introduce a smaller and well-defined number of phosphate groups. Since enzymatic modification does not provide the necessary control, we leveraged automated fast-flow peptide synthesis (AFPS) to introduce site-specific phosphoserine residues [41–43] . A recently optimized method for the reliable incorporation of phosphorylated amino acids using AFPS [44] allowed us to synthesise RS domain variants containing one (RS-pS_199_, RS-pS_223_, RS-pS_225_), two (RS-pS_223,225_ termed RS-2×pS) or four phosphoserines (RS-pS_199,209,217,225_ termed RS-4×pS, see Figure 2a and Table S3). Due to synthetic challenges of incorporating two phosphoserines in close proximity [45], we did not follow the SRPK1 phosphorylation pattern from the centre to the N-terminus of the RS domain but spaced them more widely in the RS-4×pS construct. Furthermore, as commonly observed with incorporating multiple phosphorylated residues, increasing side reactions led to an overall reduction in yield [45], preventing us from synthesising RS domain peptides with more than 4 phosphoserines.

**FIGURE 2.**
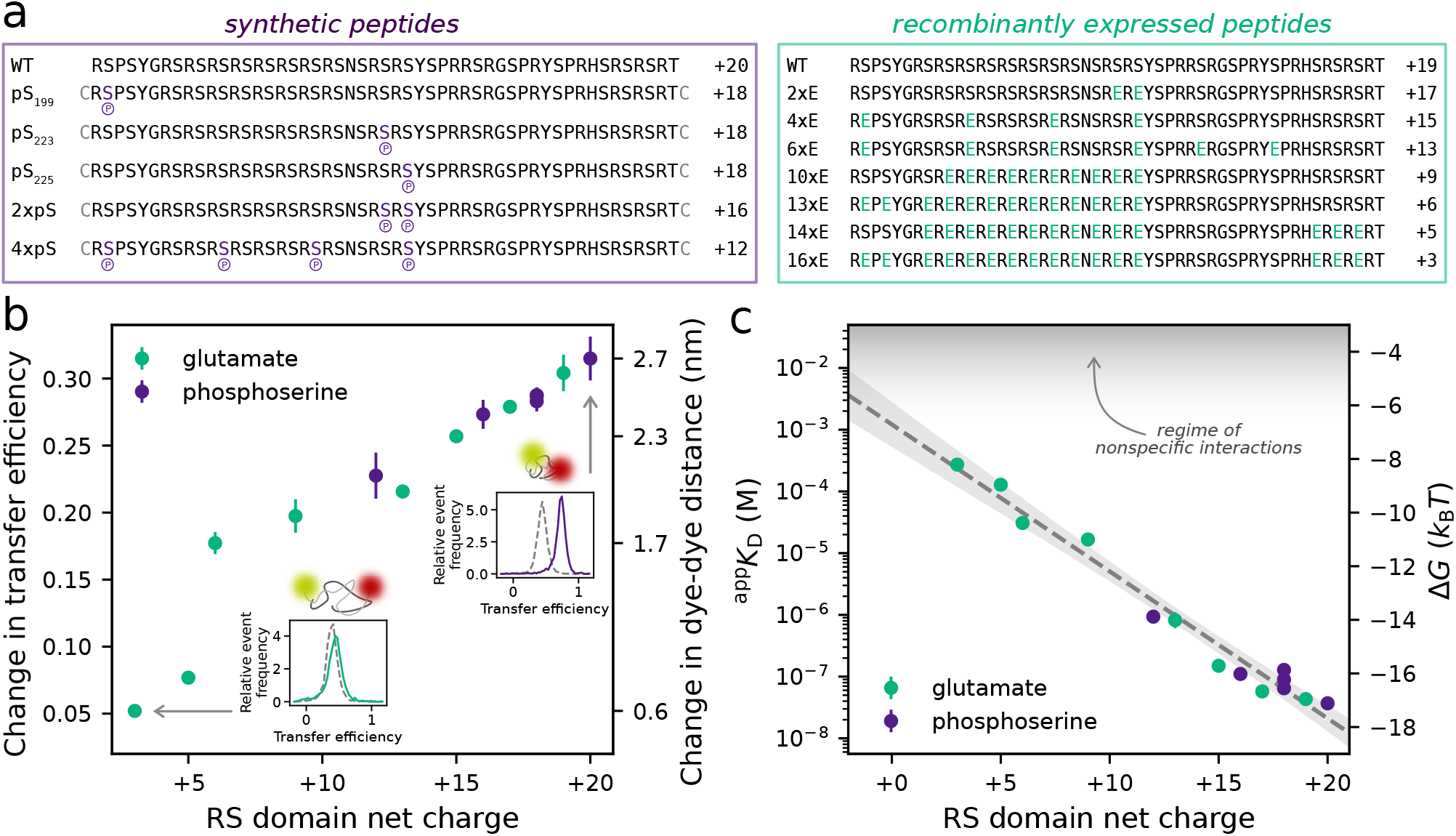
Changes in net charge fine-tune RS domain-RNA interactions. (a) Overview of peptide sequences and net charges. Altered amino acids are highlighted in purple (phosphoserine) and green (glutamate). Phospho-peptides were synthesized with terminal cysteines, which were found not to affect RNA-RS domain interactions (Figure S8). Note that synthetic peptides contain a C-terminal amide group (uncharged at pH 8) instead of a carboxyl group (negatively charged at pH 8). (b) Compaction of rU33^CA/GGA^ RNA upon RS domain binding as a function of RS domain net charge (in units of the elementary charge *e*), plotted both in terms of the measured change in transfer efficiency (left axis) and as a change in donor-acceptor dye distance of the RNA (right axis). The data points show the average and standard deviation of at least triplicates; error bars are those of the change in transfer efficiency. Insets show normalised transfer efficiency histograms of unbound (gray dashed) and saturated RNA (purple/green) to illustrate the degree of compaction at either extreme of RS domain net charge. For representative histograms of RNA bound to each RS domain construct, see Figure S10. (c) Apparent equilibrium dissociation constant of the RS domain-rU33^CA/GGA^ complex and corresponding free energy of binding as a function of RS domain net charge. The dashed line shows a fit of Equation (1) to the data, using *ε*_*r*_ = 80, *T* = 295 K, *q*_RNA_ = −34 *e* (taking into account the charges of both RNA and dyes) and an ionic strength of 0.2 M, yielding ⟨*r*⟩ = 2.08 ± 0.02 nm and Δ*G*_0_ = − 6.7 ± 0.4 *k*_B_*T*. The shaded band around the fit line corresponds to 95% confidence intervals. The area shaded in gray at the top highlights the affinity regime where interactions can occur due to nonspecific stickiness [54]. All data shown represent the average and standard deviation of at least triplicates. In most cases, the error bars are smaller than the symbols.

We thus complemented our set of synthetic phosphopeptides with a series of bacterially expressed RS domain variants in which serines were replaced with glutamate residues. This ‘phosphomimetic’ approach is commonly used to imitate posttranslational phosphorylation in cell biology and biochemistry [see 46, and references therein] and gives us the opportunity to obtain peptides across a wide range of net charge (Figure 2a). Phosphomimetic variants can obviously not be expected to be fully equivalent to phosphorylated protein [47–53], but the availability of RS domain phosphopeptides from AFPS and phosphomimetic variants across the same range of net charge (+12 to +20) allows us to quantitatively compare their affinities to RNA and address the question of whether glutamate-based phosphomimetics are suitable for imitating the effect of phosphorylation on IDP-RNA binding.

To quantify how the affinity to RNA is affected by a change in the net charge of the RS domain, we titrated donor/acceptor-labeled rU33^CA/GGA^ RNA with our thirteen different synthetic phospho-peptides and glutamate phosphomimetic RS domain variants. Since peptides bearing two and four phosphoserines were prone to aggregation, we performed all titrations in the presence of 1 M urea, an uncharged denaturant that is expected to have only minor effects on charge-dominated interactions. Indeed, compared to 0 M urea, the change in transfer efficiency between unbound and bound populations at 1 M urea differed by <10%, and values of ^app^*K*_D_ changed only by up to a factor of ∼2 (i.e. <1 *k*_B_*T* in binding free energy, Figure S9 and Table S2). From the resulting full titration curves for all peptides, we obtained the change in transfer efficiency upon binding and ^app^*K*_D_ as a function of RS domain net charge (Figure 2b,c). The change in transfer efficiency shows that the end-to-end distance of the RNA decreases less and less upon binding as the positive net charge of the RS domain is reduced by the introduction of negatively charged residues (Figure 2b, Figure S10). A deviation from the trend is observed for RS-10×E and RS-13×E, possibly due to their block-copolymeric character. Concomitant with the decrease in compaction of the complex, the affinity between RNA and RS-domain decreases from ^app^*K*_D_ ≈ 40 nM for the WT sequence to 270 ± 30 µM for RS-16 × E. Notably, the free energy of binding inferred from ^app^*K*_D_ scales roughly linearly with net charge, and the charge distribution across the sequence appears to only play a secondary role for the affinity. This contribution of charge location is best illustrated by the data for the three different single-phosphoserine peptides (Table S2), which show that affinity can vary slightly depending on the position of the phosphorylation site, making charge patterning one likely source of the residual deviations from the linear trend.

If we follow a very simple approach and assume the binding free energy, Δ*G*, to consist of a charge-independent contribution, Δ*G*_0_, and an electrostatic term described by a screened Coulomb interaction, essentially a mean-field description of the dynamically distributed charges in the two disordered binding partners, we can approximate the charge dependence of Δ*G* as [2]

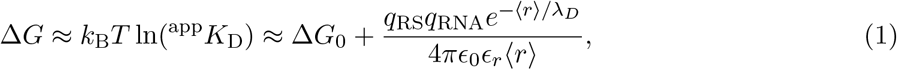

where *q*_RS_ is the net charge of the RS domain variant; *q*_RNA_ is the net charge of the RNA, separated from the RS domain by an effective average distance ⟨*r*⟩; *λ*_*D*_ is the Debye screening length at our ionic strength of 0.2 M; *ϵ*_0_ is the permittivity of vacuum; and *ε*_*r*_ is the dielectric constant of water. Fitting this linear dependence on *q*_*RS*_ to our data (Figure 2c), we obtain Δ*G*_0_ = −6.7 ± 0.4 *k*_B_*T* for the charge-independent contribution, and an average change in Δ*G* of −0.55 ± 0.03 *k*_B_*T* per single charge at near-physiological ionic strength. Extrapolated to a net charge of -13, the fit result suggests that an RS domain bearing 16 phosphorylations would exhibit an affinity for the rU33^CA/GGA^ RNA in the molar range, corresponding to the loss of binding we observed experimentally. An RS domain with a net charge of zero would correspond to millimolar affinity, indicating that the non-electrostatic contribution to binding is small. Last but not least, the linear dependence of Δ*G* on *q*_RS_ and the slope we observe is independent of whether the charge is altered by phosphorylation or by introducing acidic residues, implying that the charge of two glutamate residues is required to mimic one phosphoserine. On the one hand, it is thus obvious that a single acidic amino acid is not appropriate for imitating a phosphate group, but on the other hand, introducing two glutamates, each bearing a charge of -1 at pH 8, is remarkably accurate in accounting for a phosphate group in the charge-dominated interaction between the RS domain and RNA.

## DISCUSSION

SR proteins such as SRSF1 exemplify extreme cases of post-translational modification in which extensive phosphorylation inverts the net charge of regions highly enriched in positively charged residues. We find the electrostatic interactions of the disordered RS repeat domain to be the main driver of non-sequence-specific protein-RNA interactions, which can result in similar or even higher affinity than from the sequence-specific binding of folded RRMs. The binding affinity to RNA strongly depends on the phosphorylation state of the RS domain: it decreases as the positive net charge is reduced, and the variant heavily phosphorylated by SRPK1 exhibits no RNA affinity. We observe a remarkable linearity of the binding free energy on the net charge of the RS domain, and the change in free energy of ∼ 0.5 *k*_B_*T* per charge unit is surprisingly independent of whether the net charge is altered by phosphorylation or by introducing acidic amino acid residues. This finding implies that the use of glutamate as phosphomimetic residues, which is common practice in molecular biology [46], can be a valid approach for imitating the change in net charge for this type of protein-RNA interactions if two acidic residues are used to mimic the effect of one phosphate group.

Our findings highlight several practical considerations that can be applied to a variety of systems. First, mimicking one phosphorylation by two glutamates is a good rule of thumb for electrostatically driven interactions at a pH sufficiently far from the respective *pK*_*a*_ values of the phosphorylated and phosphomimetic amino acids. Although phosphomimetics fail to capture the charge density and hydration properties of phosphorylated amino acids [55], they remain an attractive choice when site-specific phosphorylation cannot be achieved. However, whether or not such double-phosphomimics accurately imitate the phosphorylated protein will depend on the specific system and its interaction partners [47, 56]. Second, in cases where only a single-residue substitution is feasible for each phosphorylation site, the interpretation of the data has to account for effects from the difference in net charge. For example, depending on the context, phosphomimetic variants of SRSF1 only partially reproduce the effects of binding to interaction partners and localization of the phosphorylated WT[10, 57].

The roughly linear dependence of the binding free energy on the RS domain net charge is in line with previous results on electrostatic interactions involving IDPs [2, 3, 58], even though the systems under study differ in several respects: protein-protein *vs*. protein-nucleic acid interactions; interactions between one structured and one disordered binding partner *vs*. two disordered binding partners; differences in net charge, charge clustering and length of the IDPs; and the number of negative charges added or positive charges removed. Additionally, the measurement conditions in these studies differed in pH (7 < pH < 8) and ionic strength (0.06M < IS < 0.2M). Hence, it is remarkable that, despite these variations, the change in binding free energy per unit charge is consistently approximated by a linear dependence on the net charge of the ligand. Collectively, these findings indicate that, for electrostatically driven systems, phosphorylation, and charge content in general, play a central role in tuning binding affinity, with the overall net charge exerting a stronger influence than the specific positioning or local environment of individual charges. As such, similar principles of charge interactions are expected to be important for the majority of protein-RNA and protein-protein interactions that are regulated by SRSF1 phosphorylation, e.g. the inter- and intramolecular interactions between RS domain and RRMs [59–62], binding to other nuclear proteins involved in splicing regulation [36], and SRSF1 phase separation [23, 24, 40].

Splicing is a tightly regulated and dynamic process involving the controlled binding and dissociation of many proteins and ribonucleoprotein complexes, many of which compete for overlapping or neighboring RNA sequences [63]. SRSF1 is fully phosphorylated when localizing to active transcription sites, where it recognizes splicing enhancer sequences within nascent pre-mRNA transcripts [64, 65]. If SRSF1 were not fully phosphorylated, RS domains lacking or containing only a small number of phosphorylations could lead to non-sequence-specific binding of SRSF1 outside of enhancer motifs and non-productive compaction of RNA. Both effects could physically limit access of other splicing factors to their cognate binding motifs. That is, in the context of splice-site recognition and initial spliceosome recruitment, phosphorylation appears to function as a regulatory switch that reduces SRSF1-RNA interactions and promotes protein-protein interactions [14]. A quantitative understanding of the role of phosphorylation in RS domains and highly charged IDPs in general is thus essential for elucidating the resulting regulatory processes.

## METHODS

### Preparation and labeling of RNA

RNAs modified with a 5’ di-thiol group and a 3’ or internal amino group (Table S1) were purchased from Integrated DNA Technologies (IDT). RNAs were dissolved to ∼100 µM in 100 mM NaP pH 7 or RNAse-free water. TCEP was added to a final concentration of 5 to 10 mM to reduce the dithiol. After incubation at room temperature for ∼30 min, all RNAs but G19 were buffer exchanged into 100 mM NaP pH 7 by centrifugal filtration (Amicon Ultra-0.5 MWCO 3 kDa). G19 was buffer-exchanged into 20 mM NaP pH 7, 6.4 M GdnHCl. An approximately 10-fold excess of CF660R-maleimide (Sigma-Aldrich), dissolved in a total volume of 5 µL DMSO, was added to the RNA solution and incubated on ice overnight or at room temperature for 1 h. Subsequently, the pH was raised to 8 by addition of 0.5 M NaP pH 8 (all RNAs but G19) or 20 mM NaP pH 8.6, 6.4 M GdnHCl (G19), and a 10-fold excess of Cy3B-NHS (Cytiva or Lumiprobe) dissolved in 5 µL DMSO was added. The reaction was incubated further at 37 °C for 2 h. Unreacted dye was removed by buffer exchange using ZebaSpin desalting columns (ThermoFischer Scientific, MWCO 7 kDa), or by ethanol precipitation. The labeled RNAs were applied to a Reprosil Pur 200 C18-AQ column (4×250 mm, 5 µm, Dr. Maisch) equilibrated in 0.1 M acetic acid, 0.1 M triethylamine, 3.75% acetonitrile, and eluted with gradients of 0.25 to 1.5 %/min to achieve separation of labeled species for each RNA. Fractions containing donor/acceptor-labeled RNA were lyophilised and resuspended in RNase-free water, and stored at −80 °C.

### Plasmids and molecular cloning

Sequences of full-length SRSF1 containing the solubilising mutations Y37S and Y72S in RRM1 [16], a construct containing only the RRMs including the same mutations (RRM12, amino acids 1-196 of SRSF1), and the RS domain (amino acids 198-248) were subcloned from a pET24b plasmid into a pSUMO plasmid (kind gift of B. B. Kragelund, University of Copenhagen, Denmark) using NEB Gibson Assembly Master Mix (NEB,E2611) according to manufacturers specifications (for primers, see Table S4). Glutamate variants were ordered as synthetic genes from Integrated DNA Technologies (IDT) and cloned into pSUMO, linearised using the pSUMO Fwd and Rev primers listed in Table S4 using the NEBuilder HiFi DNA Assembly Master Mix in a 5 µL volume, but otherwise according to the manufacturer’s protocol. The plasmid bearing Ulp1 protease with a C-terminal His6-tag was a kind gift from the Kragelund Lab. GB1-His_6_-SRPK1-His_6_ in a pET24b plasmid was a kind gift of Guillaume Hautbergue (University of Sheffield). DNA sequences were confirmed by Sanger sequencing (Microsynth).

### Preparation of recombinantly expressed proteins and peptides

For a detailed description of the expression and purification of each protein or peptide, please refer to the Supplementary Experimental Methods. Amino acid sequences are listed in Table S5. In brief, all proteins were expressed in *E. coli* BL21(DE3) derivatives, cell pellets were lysed mechanically and purified by immobilised metal ion affinity chromatography (IMAC). Ulp1 protease was dialysed into storage buffer containing 50% glycerol. SRSF1, RRM12 and RS domain variants were further purified by ion exchange (IEX) chromatography and, if necessary, size exclusion chromatography. The His_6_-SUMO-tag was removed by cleaving with Ulp1 followed by further purification (reverse-IMAC or IEX chromatography). SRPK1 was further purified by IEX chromatography. All protein samples were flash-frozen in liquid N_2_ and stored at −80 °C. Protein concentrations were determined by absorbance at 280 nm and converted to molar concentrations using the respective theoretical extinction coefficients. Protein masses were confirmed by electrospray ionisation mass spectrometry (ESI-MS) at the Functional Genomics Center Zurich.

### *In vitro* phosphorylation of SRSF1 and the RS domain

To obtain phosphorylated SRSF1 (pSRSF1), phosphorylation reactions were carried out with 2.5 to 5 µM His_6_-SUMO-SRSF1 or SRSF1 and 1.7 to 5 µM GB1-His_6_-SRPK1-His_6_ in 50 mM Tris-HCl pH 8, 10 mM MgCl_2_, 0.5 mM TCEP, 1 mM ATP. Reactions were incubated at 37 °C for ∼3 h. The phosphorylated protein was then purified by IEX chromatography using a MonoS 5/50 GL (GE Healthcare). Prior to each purification run, the column was incubated in 1 mg/mL pepsin dissolved in 0.5 M NaCl, 0.1 M acetic acid for at least 1 h at room temperature before cleaning with 2 M NaCl and 1 M NaOH according to the manufacturers protocol. The phosphorylation reaction was diluted to a final NaCl concentration of 100 to 150 mM prior to loading onto the column equilibrated in 20 mM HEPES pH 8, 150 mM NaCl, 5% glycerol, 0.01% Tween, 0.5 mM TCEP. The column was then washed with 10 CV of the equilibration buffer. pSRSF1 was collected from the flow through and the wash. SRPK1 and partially phosphorylated or unphosphorylated SRSF1 were recovered from the column using a gradient of 0.15 to 1.5 M NaCl. Flow-through and wash fractions containing pSRSF1 were pooled and concentrated using Amicon Ultra 0.5ml centrifugal filters (10 kDa MWCO, cellulose, Merck Millipore). The concentrated protein was then buffer-exchanged into 20 mM HEPES pH 8, 200 mM NaCl, 0.5 mM TCEP using ZebaSpin Desalting columns (7 MWCO, Thermo Scientific), frozen in liquid nitrogen and stored at −80 °C. The total number of phosphorylations was determined by ESI or MALDI mass spectrometry at the Functional Genomics Center Zurich.

To obtain phosphorylated RS domain, phosphorylation reactions were carried out with equimolar RS domain and SRPK1 (usually 5 to 6 µM of each) in 50 mM TrisHCl pH 8, 10 mM MgCl_2_, 2 mM DTT. Reactions were started by the addition of two-fold excess of ATP per serine residue (effectively 40-fold excess over protein concentration) and incubated at 37 °C for 30 min before adding GdnHCl to a final concentration of 4 M to stop the reactions. The mixture was applied to a 250×4.6 mm, 5 µm ReproSil Gold 200 C18 column (Dr. Maisch) equilibrated with H_2_O, 0.1% TFA, 5% methanol. Proteins were eluted using a gradient of 23 to 30 % methanol over 30 min. Fractions containing the phosphorylated RS domain were combined and lyophilized. The protein was resuspended in 20 mM HEPES pH 8, 150 mM NaCl and stored at −80 °C. To remove Fe^3+^ adducts, samples were dialysed against 20 mM HEPES pH 8, 150 mM NaCl containing 50 µg/ml Chelex (Bio-Rad Laboratories) for 1 hour at room temperature. The total number of phosphorylations was determined by ESI mass spectrometry at the Functional Genomics Center Zurich.

### Peptide synthesis and purification

For a detailed description of peptide synthesis and purification, please refer to the Supplementary Experimental Methods. Amino acid sequences of the synthetic peptides are listed in Table S6. In brief, peptides were synthesised on NovaPEG Rink Amide resin using an automated-flow system built in the Hartrampf lab [66]. Crude peptides were purified by semi-preparative reverse-phase HPLC (RP-HPLC). Both crude and pure peptides were analyzed by analytical ultra-high performance liquid chromatography (UHPLC) and liquid chromatography with high-resolution electrospray ionization mass spectrometry (LC-MS). All phosphopeptides were synthesised with terminal cysteines for future studies. To ensure that their addition did not affect the interaction with RNA, we synthesised the unphosphorylated wild-type peptide with and without terminal cysteines and compared them, especially regarding their affinity to RNA, using single-molecule FRET measurements (see Figure S8). An overview of purities and yields is given in Table S3. Figures containing UHPLC and LCMS data are listed in Appendix A.

### Single-molecule FRET of freely diffusing molecules

Measurements of freely diffusing molecules were performed on a MicroTime 200 (PicoQuant) or a custom-built confocal single-molecule fluorescence instrument. On both instruments, lasers were operated in pulsed mode with interleaved excitation [67] at a pulse repetition rate of 20 MHz. The MicroTime 200 was equipped with an Olympus UplanApo 60x/1.20 W objective. The donor was excited by light from a supercontinuum laser (SuperK EVO HP SC-04, NKT Phototnics) filtered to 532/3 nm (BrightLine HC, Semrock) at 50 µW, as measured at the back aperture of the objective. The acceptor was excited by light from a 640 nm diode laser (LDH-D-C-640, PicoQuant) at 55 µW, as measured at the back aperture. Fluorescence emitted by the sample was collected by the same objective lens and separated from backscattered light by a triple-band mirror (ZT405/530/630RPC, Chroma Technology). Scattered light was further removed by a long-pass filter (LP532, Chroma Technology), before the emitted photons passed a 100 µm pinhole. The emitted photons were separated into four detection channels with a polarizing beam splitter and two dichroic mirrors (635DCXR, Chroma Technology). Donor and acceptor emission was further filtered with ET585/65M band-pass and LP647RU long-pass filters (both Chroma Technology), respectively. Photons were detected by four single-photon avalanche diode detectors (SPCM-AQR-15, PerkinElmer Optoelectronics). The arrival times of single photons were recorded with a HydraHarp 400 counting module (PicoQuant) with a resolution of 16 ps. The custom-built instrument was operated with a different donor excitation laser (EXW-12 SuperK Extreme, NKT Photonics), as described previously [68], with the wavelength selected by a 520/15 nm band-pass filter (BrightLine HC, Semrock) at 45 µW, as measured at the back aperture of the objective. Acceptor excitation and all other filters were as described for the MicroTime 200.

Apparent dissociation constants were determined using single-molecule titrations of freely diffusing molecules. Unless otherwise stated, for all RNAs but rU59^CA/GGA^, titrations were performed using 50 pM labeled RNA in 20 mM HEPES pH 8, 200 mM mM NaCl, 0.01% Tween-20, 0.2 mg/mL BSA or recombinant HSA (both NEB), 10 mM DTT, 0.1 U/µL RNAse inhibitor (Applied Biosystems). Titrations with the respective peptides/proteins were performed in untreated µ-slides (Ibidi) by starting at saturating conditions and then sequentially diluting the protein concentration with buffer containing only donor/acceptor-labeled RNA until the data obtained from the last titration point(s) were indistinguishable from a sample containing only RNA. To keep the ionic strength constant over the whole titration, the amount of NaCl added to the measurement buffer was adjusted to compensate for the amount of NaCl introduced by the protein storage buffer.

We found that the number of photon bursts detected per second decreased with increasing protein concentration (Figure S2a): while barely noticeable for RRM12, it was pronounced for the RS domain and full-length SRSF1 at high protein concentrations. Concurrently, we observed a new population at lower transfer efficiencies (asterisk in Figure 1b) originating from slower-diffusing species, as evident in the overall acceptor-donor fluorescence correlations (AD-FCS) (Figure S2b) and by subpopulation-specific AD-FCS (insets in Figure 1b and Figure S1), in line with the propensity of SRSF1 to phase-separate and aggregate, even at low concentrations [23, 24]. Although the presence of higher-molecular weight species can complicate the interpretation of the titrations, the advantage of our single-molecule approach is that we can identify and distinguish these species spectroscopically and adapt our analysis accordingly (see Figure S2 and data analysis methods below).

For SRSF1 without solubilizing SUMO tag, the burst loss was so large that full titration curves could not be obtained. Since we found the solubility of SRSF1, RRM12, and the RS domain to increase with a solubilisation tag, SUMO-tagged and cleaved proteins were compared in standard measurement conditions using the rU33^CA/GGA^ RNA, and affinities were found to be within error (Figure S3). Importantly, we could obtain titrations to saturation with SUMO-SRSF1 (Figure 1c).

We found that the addition of urea also improved data quality by reducing burst loss at high protein concentrations. Binding of SUMO-RRM12, and the synthetic and expressed RS domains to the rU33^CA/GGA^ RNA was tested at different urea concentrations up to 1 M (Figure S9). For the RS domain peptides, we found that the effect of urea on the binding affinity to RNA is negligible, and hence all measurements used to determine the relationship between of RS domain net charge and its affinity to RNA were performed in the presence of 1 M urea. In contrast, urea clearly affects specific interactions, as observed with the RRM12 construct (either due to partial unfolding of the domains, or due to interfering with base recognition).

### Data analysis of binding titrations

Data analysis was carried out using the Mathematica (Wolfram Research) package Fretica (https://github.com/SchulerLab) and custom-written Python scripts [69–73]. Recorded photon counts were corrected for background photons, acceptor direct excitation, channel cross-talk, differences in detector efficiencies, and quantum yields of the dyes, as previously described [74]. Signal from large aggregates or oligomers was removed by binning photons in 1 s bins and identifying bursts with intensities higher than three standard deviations above the mean. Fluorescence bursts from individual molecules were identified by dual-channel burst search [75]. Successive photons separated by inter-photon times of less than 150 µs were combined and defined as a burst if the corrected total number of photons detected was more than 60 [76]. To select the molecules with active donor and acceptor, only bursts with stochiometry ratio, *S*, within the interval 0.2 < S < 0.7 were retained.

Ratiometric transfer efficiencies were obtained for each burst from *E* = *n*_A_/(*n*_A_+*n*_D_), where *n*_D_ and *n*_A_ are the counts of donor and acceptor photons, respectively, corrected for direct excitation, channel crosstalk, and differences in fluorescence quantum yields and detection efficiencies. The transfer efficiencies of all bursts were binned into histograms. Fluorescence correlations of acceptor and donor photons were calculated as described previously [77]. Subpopulation-specific FCS was performed on selected bursts within a given range of transfer efficiency of the FRET population of interest.

For all data acquired in native buffer conditions to compare interactions between different RNAs and protein constructs, transfer efficiency histograms were globally fitted with Gaussian peak functions for the free and bound populations, respectively, keeping the mean transfer efficiency and the width of the respective Gaussian functions as shared parameters. The peaks were then integrated to extract the fractions of the bound and unbound populations[29, 30]. To obtain the midpoint of the binding titrations, data of the fraction bound, *θ*, vs. protein concentration, *c*_prot_, were fitted with a binding isotherm [29, 78]

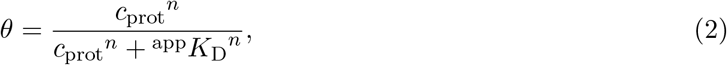

where ^app^*K*_D_ is the apparent dissociation constant corresponding to the midpoint, and *n* is treated as an adjustable parameter. For consistency, affinity data as a function of urea concentration (Figure S9) were all based on fitting the fraction bound.

In the experiment series with RS domain peptides of different net charge, the RS-RNA binding equilibrium varies from being in slow exchange for peptides of high positive net charge (e.g. WT peptides and single phosphorylation variants) to being in fast exchange for peptides approaching neutral net charge (e.g. 14xE and 16xE). Hence, we computed the mean transfer efficiency over all selected bursts with active donor and acceptor dye for each protein concentration, instead of fitting independent peaks, to be consistent across all data sets. Note that the presence of urea ensured that oligomerisation was usually below the detection limit. Data of mean transfer efficiency, *Ē*, vs. protein concentration were fitted with

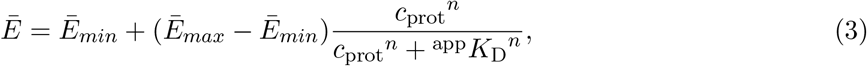

where *Ē*_*max*_ and *Ē*_*min*_ are the mean transfer efficiencies of the fully bound and unbound states, respectively. Where possible, uncertainties in 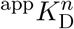 are given as standard deviations estimated from multiple measurements. Otherwise, we assume an uncertainty of a factor of two as an estimate[28], based on the typical variability we have observed among binding measurements.

To calculate the net charge of RNAs and proteins, we use the following values (all in units of elementary charge): *q* = −1 per RNA nucleotide (due to backbone phosphate), *q* = −1 for CF660R, *q* = 0 for Cy3B, *q* = +1 for arginine, *q* = −1 for glutamate, *q* = −2 for phosphoserine, *q* = −1 for the C-terminal carboxyl group, *q* = 0 for the C-terminal amide group, and *q* = +1 for the N-terminal amino group. We note that the N-terminal amine might be partially deprotonated at pH 8. However, this would only introduce an offset for all peptides and would thus not change the outcome of the data analysis or the conclusions.

### Quantifying the change in RNA compaction upon binding

We use the change in transfer efficiency upon binding to estimate the corresponding change in average end-to-end distance of the RNA. To minimize variability due to burst loss and poor photon statistics at high protein concentrations, we averaged several saturated transfer efficiency histograms (usually the two highest protein concentrations) for the fully bound state. Every titration data set contains a measurement of free, unbound donor/acceptor-labeled RNA (without protein). We fitted both the saturated and the RNA-only transfer efficiency histograms with Gaussian peak functions to obtain the mean transfer efficiency of the bound (*E*_*b*_) and unbound population (*E*_*u*_), respectively. Multiple measurements of the difference in transfer efficiency, Δ*E*_*b*→*u*_ = *E*_*b*_ − *E*_*u*_, were then averaged for each RS domain peptide (see Figure 2b).

For converting the transfer efficiency change into a change in average end-to-end distance, we approximate the distribution of the inter-dye distance, *r*, by a Gaussian chain with the probability density function

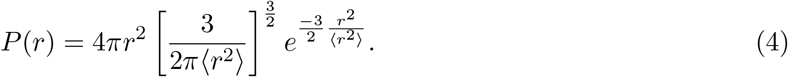

From the measured mean values of *E*_*b*_ and *E*_*u*_, we obtain the respective root-mean-square distances, 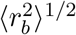 and 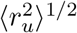, by numerically solving

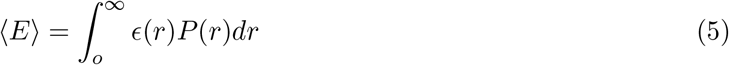

with respect to ⟨*r*^2^⟩ [79], where *ϵ*(*r*) is the transfer efficiency

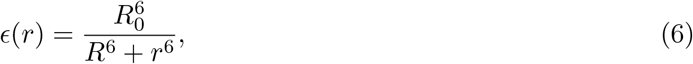

with *R*_0_ being the Förster radius (*R*_0_ = 6.0 nm for Cy3B-CF660R [80]). Values obtained from several titrations were averaged for each RS domain peptide.

As an estimate of the systematic uncertainty dominated by instrument calibration, we report an uncertainty of ∼ 0.03 on single mean transfer efficiency values [27, 80, 81]. Note, however, that the statistical uncertainty of the measurements taken on the same instrument on the same day are much lower (typically ≤0.01 [80]). The measured differences in transfer efficiency are thus more accurate than individual transfer efficiencies and less dependent on systematic errors, and hence we quote one standard deviation based on multiple measurements as an uncertainty for the difference in transfer efficiency between bound and unbound RNA.

### Single-molecule FRET of surface-immobilised molecules

Due to the much larger net charge of the 60-nucleotide RNA than of the RS domain, we could observe two RS domain molecules binding to one rU59^CA/GGA^ molecule. However, as both binding events were within one order of magnitude in terms of affinity, we chose to perform surface experiments to best resolve single- and double-bound states and to obtain an accurate affinity for the 1:1 interaction. The MicroTime 200 described above was used with a 532 nm continuous wave laser (LaserBoxx LBX-532-50-COL-PP, Oxxius) instead of the white light source. The laser power for acceptor excitation that was used to scan the surface and locate immobilized molecules was set to 5 µW (Sepia PLD 828 Multichannel Picosecond Diode Laser Driver, PicoQuant). Slides for surface measurements were prepared using PEGylated, biotinylated fused silica cover slides (MicroSurfaces, Inc) and a hybridization chamber (Secure Seal Hybridization Chambers SA8R-2.5, Grace Bio-Labs). Before assembling the two parts, the cover slide was sonicated for 15 min in MilliQ water containing 0.1% Tween [82]. The slides were then washed with MilliQ water and dried with compressed air, before binding them to the adhesive hybridization chambers.

First, 0.2 mg/mL neutravidin in 50 mM HEPES pH 8 and 128 mM NaCl was added to the measurement chamber and incubated for 10 - 15 minutes. Afterwards, the chamber was washed three times with buffer (50 mM HEPES pH 8, 128 mM NaCl) before 10 pM donor/acceptor-labeled rU59^CA/GGA^, diluted in the same buffer, was added. After approximately one minute, the chamber was washed again with buffer to remove unbound RNA. Measurements were performed with different protein concentrations in 50 mM HEPES pH 8, 128 mM NaCl, 0.01 % Tween, 0.2 mg/ml BSA, 5 mM ascorbic acid, 5 mM methyl viologen, 5 mM protocatechuic acid (PCA) and 10 nM protocatechuate-3,4-dioxygenase (PCD). Due to the additional ions introduced by the redox and oxygen scavenger systems, the NaCl concentration was reduced to match the overall ionic strength to that used in the free-diffusion experiments. The experiment was performed under argon atmosphere to reduce dye bleaching. The donor dye was excited with a laser power of 1.4 µW, and trajectories were recorded until both dyes bleached, followed by an additional 0.5 seconds to be used for background correction during data analysis.

Fluorescence time traces were selected for analysis based on the following criteria: (i) presence of anti-correlation between donor and acceptor signals coming from all three states (unbound, single-bound, double-bound); (ii) stable total photon count rates prior to bleaching; and (iii) ≥ 0.5 s of background signal available post-bleaching. For building transfer efficiency histograms, trajectories were binned in 50 ms bins. Counts of donor and acceptor photons where corrected for background and other effects analogous to the correction of burst counts described above. For the kinetic analysis, uncorrected trajectories were used with 20 ms binning and analyzed by hidden-Markov modelling as described previously [83]. Individual labeled and immobilized RNA molecules were observed in three states: (1) free, (2) with one RS domain, or (3) with two RS domains bound. The rate coefficients of the corresponding kinetic system

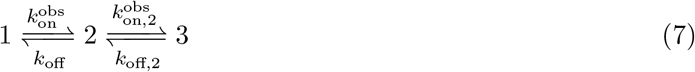

were obtained using a maximum-likelihood analysis of the trajectories, errors were determined using bootstrapping. The equilibrium dissociation constant for a single RS domain bound to RNA was determined using

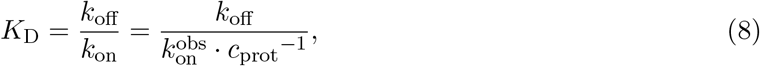

where *c*_prot_ is the concentration of the RS domain in solution. The error of the equilibrium dissociation constant was obtained by error propagation using an estimated error of 20% in protein concentration and the standard deviation of 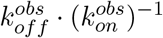 obtained from bootstrapping.

## ACKNOWLEDGEMENTS

We thank Guillaume Hautbergue for providing the pET24b-GB1-His_6_-SRPK1-His_6_ plasmid, Cristina Kim Xuan Nguyen for providing the pET24b-GB1-His_6_-TEV-SRSF1-YS plasmid, Nicholas Krebs for cloning the different SRSF1 constructs, Ella Knüpling for providing assistance with SRPK1 production, and Erik Holmstrom and Andrea Soranno for insightful discussion. Furthermore, the authors gratefully acknowledge the Functional Genomics Center Zurich (FGCZ) of the University of Zurich and ETH Zurich, and in particular Dr. Serge Chesnov, for the support on mass spectrometry analysis. Financial support for this project was provided by the Swiss National Science Foundation: CRSII5 205922 and CRSII5 205922/2 (FA, BS), 10006187 (BS), 200021 200,865 (NH) and 10000072 (NH). Additionally, MS gratefully acknowledges the support provided through a FEBS LTF.

## AUTHOR CONTRIBUTIONS

MS, BS, HB, NH designed research. MS, SH, LF, NK and MN prepared samples. HB synthesised the peptides. MS, LF, SH, AP and HB acquired the data. MS, AP, HB analyzed the data. Technical support and analytical tools were provided by AC and DN. BS, NH and FA supervised the project. MS and BS wrote the manuscript with input from all authors.

## ADDITIONAL INFORMATION

### Supplementary information

The Supporting Information is available online, and contains the supplementary figures, tables and methods as well as the peptide synthesis data.

