## Supplementary Information for "Phosphorylation tunes electrostatically driven protein-RNA interactions"

**CONTENTS**

I. Supplementary Figures 3

II. Supplementary Tables 10

III. Supplementary Experimental Methods 15
A. Preparation of recombinantly expressed proteins and peptides 15
1. SRSF1 15
2. RRM12 15
3. RS domain 16
4. RS domain glutamate variants 16
5. SRPK1 17
6. Ulp1 17
B. Peptide synthesis and purification 18
1. Reagents and solvents 18
2. Automated fast-flow peptide synthesis (AFPS) 18
3. Peptide cleavage and deprotection 18
4. Analytical Ultra-High Performance Liquid Chromatography (UHPLC) 19
5. Liquid Chromatography with High-Resolution Electrospray Ionization Mass Spectrometry (LCMS) 19
6. Semi-Preparative Reverse-Phase High Performance Liquid Chromatography (RP-HPLC) 20
7. Peptide synthesis of RS domain variants - peptide specific details 20

IV. References 22

A. Peptide synthesis data 23

### I. SUPPLEMENTARY FIGURES

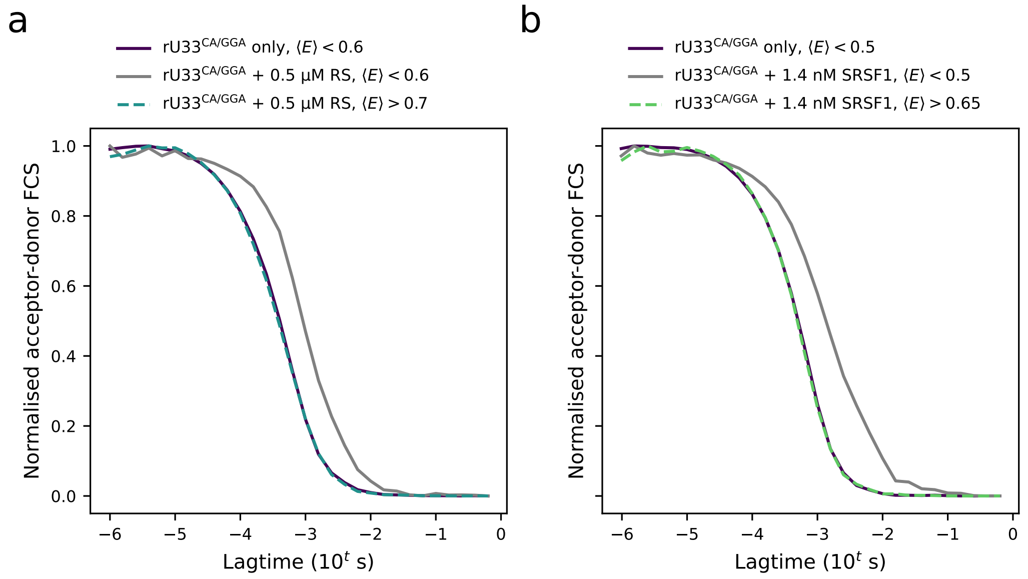


FIGURE S1. Using subpopulation-specific FCS to distinguish between monomeric and oligomeric species. Shown are the same curves displayed as insets for RS domain (a) and SRSF1 (b) histograms in Figure 1c, here overlaid with data of the respective RNA-only measurements (purple curves). At saturation, the data obtained for transfer efficiency ranges $E < 0.6$ (RS domain) and $E < 0.5$ (SRSF1) are shifted to larger lag times, indicating the presence of slower-diffusing molecules (gray curves). In contrast, molecules with transfer efficiency $E > 0.7$ (RS domain, dashed blue curve) and $E > 0.65$ (SRSF1, dashed green curve), corresponding to protein bound to RNA, diffuse on a similar timescale as the unbound RNA. Together, these data show that at saturating protein concentrations, particles with low apparent transfer efficiency are distinct from the unbound RNA and RNA bound to one protein molecule.

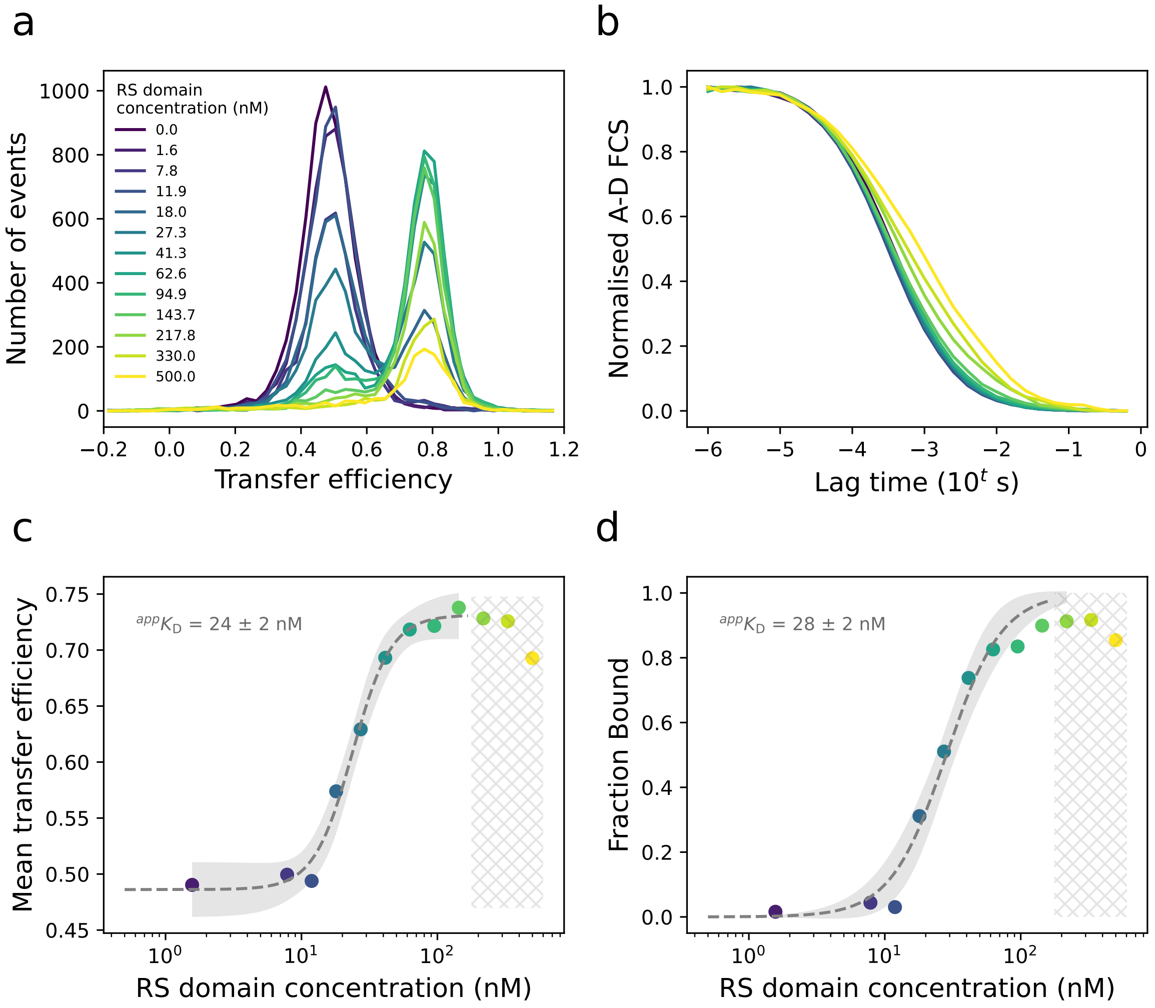


FIGURE S2. Illustration of the data analysis workflow for identifying and removing the contribution of aggregates in binding isotherms, using the rU33$^{CA/GGA}$ RNA and the RS domain as a representative example. (a) Overlay of transfer efficiency histograms of donor/acceptor-labeled RNA at different protein concentrations (see legend) showing a reduced number of bursts at high protein concentrations. (b) Overlay of normalised acceptor-donor (AD) fluorescence correlation curves for each protein concentration. At protein concentrations above ∼150 nM, the signal clearly contains slower-diffusing species in addition to the 1:1 RNA-RS domain complex. (c) At each protein concentration, the mean transfer efficiency is obtained from all selected bursts. Owing to the appearance of higher-molecular weight species with lower apparent transfer efficiencies, the mean transfer efficiency decreases at high protein concentrations. Hence, data containing substantial contributions from slower-diffusing species, as determined by FCS (hatched in gray), are excluded from the fit of the binding isotherm. This method of determining the affinity is only reliable if saturation is achieved to good approximation before burst loss becomes significant, as the top baseline is otherwise not representative of the transfer efficiency of the fully bound RNA. The gray dashed line and shaded area correspond to a fit of Equation (3) to the data and the 95% confidence band. Fit value and fit error for the apparent affinity are indicated. (d) Alternatively, the fraction of bound RNA at a given protein concentration can be obtained by globally fitting the histograms shown in (a) with two Gaussian peak functions, one for the bound and the unbound population, respectively. Since the transfer efficiencies of slower-diffusing species overlap with those of the unbound population, the apparent fraction bound will never be equal to 1 – even at saturation. Hence, data containing contributions from slower-diffusing species, as determined by FCS (b), are excluded from the fit of a binding isotherm. Since fitting isotherms to the fraction bound does not rely on fit parameters for the unbound and saturation baselines, it is still possible to obtain a reasonable estimate for the midpoint even in the presence of significant burst loss. The gray dashed line and shaded area correspond to a fit of Equation (2) to the data and the 95% confidence band. Fit value and fit error for the apparent affinity are indicated.

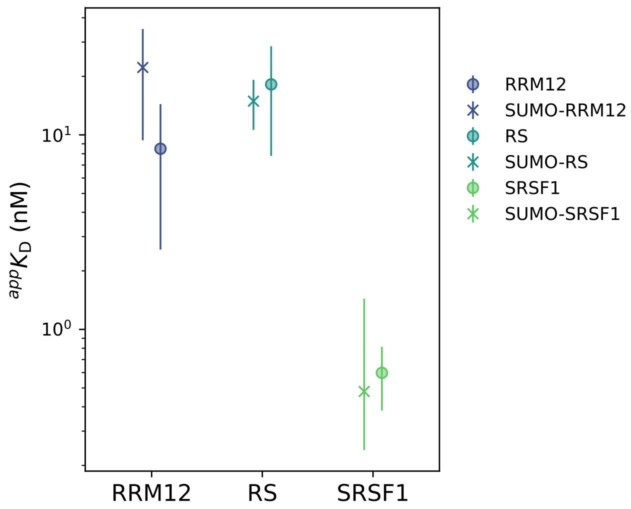


FIGURE S3. Presence of the SUMO-tag has no measurable effect on affinity. Comparison of affinities of different protein variants (see legend) to rU33$^{CA/GGA}$ RNA obtained with and without SUMO tag. For details on affinities and errors see Table S2. On average, all affinities of SUMO-tagged and un-tagged constructs are within error.

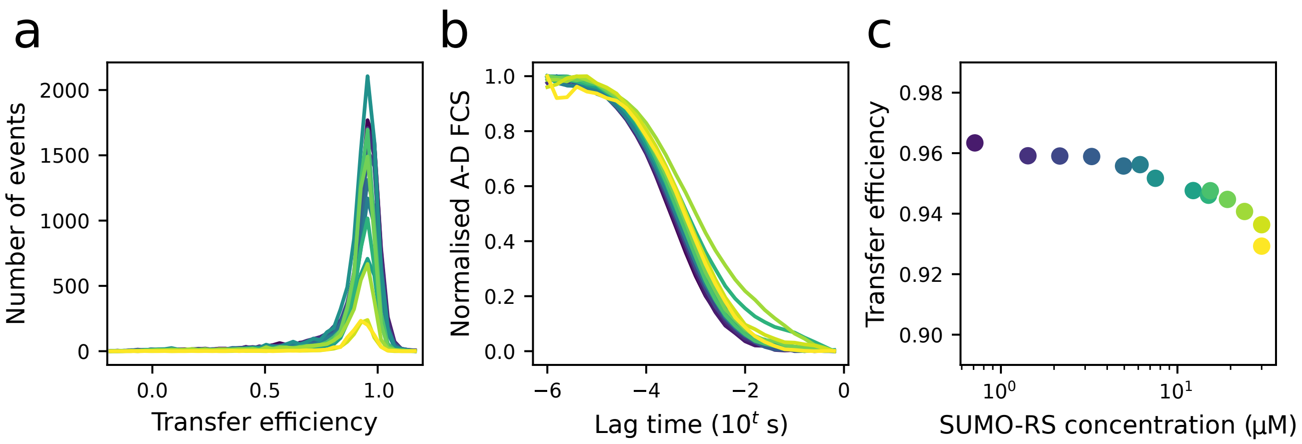


FIGURE S4. Obtaining an affinity for the interaction of the RS domain to the rU9$^{CA/GGA}$ RNA. (a) Transfer efficiency histograms of a titration of SUMO-RS into rU9$^{CA/GGA}$ show a substantial reduction in the number of bursts at the micromolar protein concentrations that are required for these measurements due to the reduced affinity of the RS domain for short ssRNAs . The SUMO-tagged construct was chosen to be able to increase the maximum protein concentration that could be used compared to the untagged RS domain. Colours corresponds to protein concentrations between 0 µM (purple) and 30 µM (yellow). Above 30 µM, burst loss was too severe to obtain meaningful histograms. (c) Fluorescence correlation curves indicate oligomerisation at high protein concentration as a likely source of the reduction in bursts. (b) Although the mean transfer efficiency (obtained by fitting Gaussian peak functions to the respective histograms in (a)) shows the onset of a binding response, it cannot be fitted meaningfully to obtain an apparent affinity. Hence, by estimating that 30 µM SUMO-RS corresponds to 5 to 20 % bound RNA, we infer an apparent affinity in the range of $150 < {}^{app}K_D < 600$ µM.

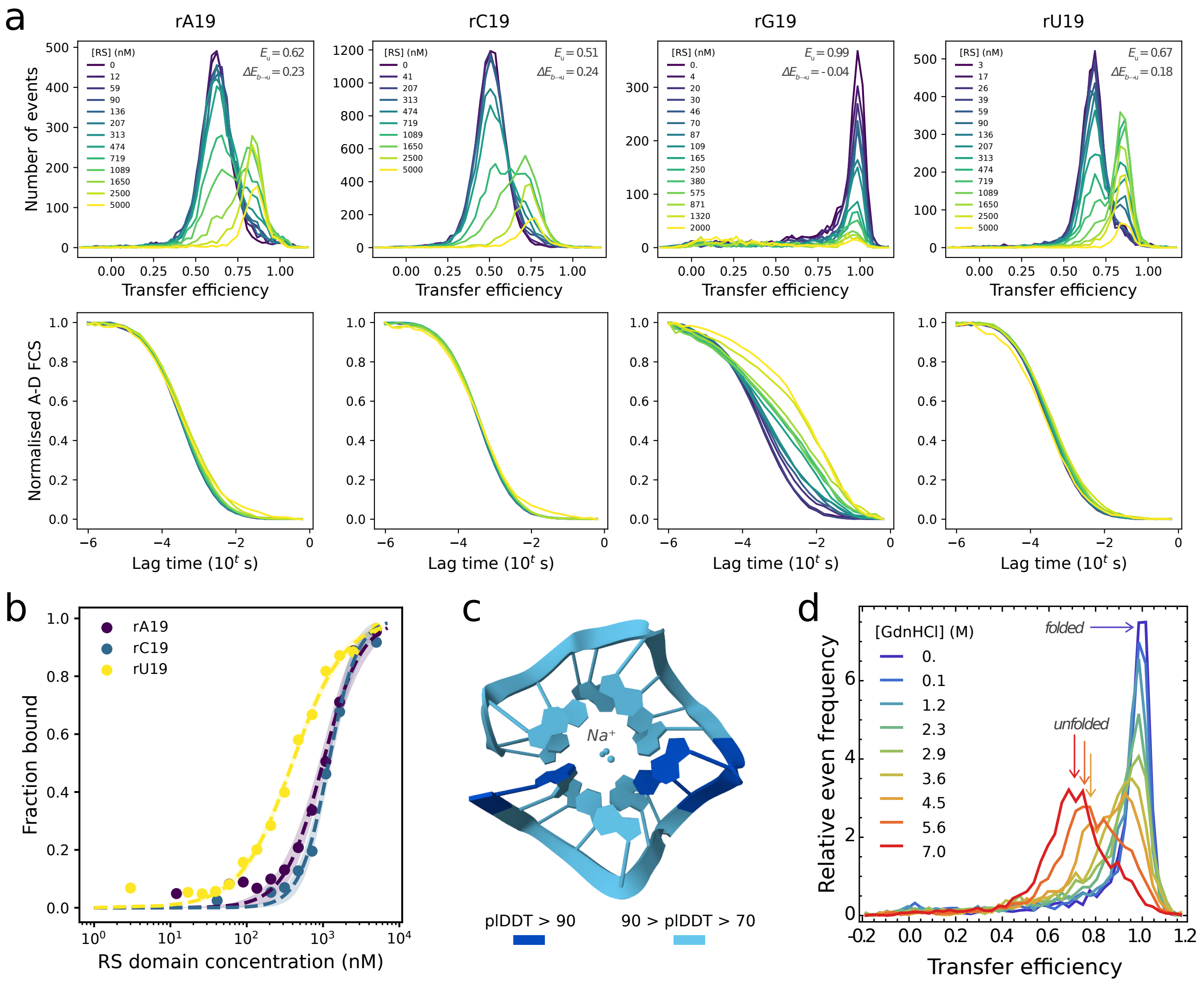


FIGURE S5. RS domain interactions with homopolymeric single-stranded RNA (ssRNA) containing different nucleobases. (a) Transfer efficiency histograms (top) and acceptor-donor (AD) fluorescence correlation curves (bottom) of titrations of donor/acceptor-labeled 19 bp ssRNAs with RS domain. Note that data obtained for rG19 differ from the other ssRNAs, likely due to it folding into a G-quadruplex-like structure (see (c) and (d)). All RNAs except rG19 are relatively expanded, with transfer efficiencies between 0.51 and 0.67, and they show compaction upon RS domain binding. In contrast, rG19 exhibits a very high transfer efficiency near 1, and addition of the RS domain results in only a small shift to lower transfer efficiency ($\Delta E \sim 0.04$) and pronounced aggregation/oligomerisation, as evident from the FCS measurements shown below. It has previously been proposed that the RS domain can unfold G-quadruplexes [1]. We do not observe indications for G-quadruplex unfolding, and based on our observations of ssRNA-RS domain interactions and previous findings regarding nucleic acid interactions with positively charged proteins [2], we expect binding of the RS domain to favour RNA compaction (as observed for rA19, rC19, and rU19) and structure formation. (b) Binding isotherms obtained from the transfer efficiency histograms shown in (a), including fits to Equation (2) and 95% confidence bands. The apparent affinities are listed in Table S2. Due to extensive aggregation and only a minor shift in transfer efficiency upon the addition of RS domain, it was not possible to obtain a meaningful binding isotherm for rG19. Although preferential interactions with arginine have been suggested for guanosine and cytosine [1, 3], we suggest that the minor differences in affinity observed ($\Delta G < 2\ k_BT$) could arise from differences in base stacking within ssRNAs (e.g. $\Delta G \sim 1\,k_BT$ for 2-3 stacked bases [4, 5]). (c) The model of the rG19 structure and 3 $Na^+$ ions generated with AlphaFold3 [6] resembles a G-quadruplex. However, due to the absence of non-G bases in loops, the topology appears strained and less ordered than usual for G-quadruplexes. Based on this model, the predicted distance between the terminal fluorophore attachment sites would be ∼2 nm, resulting in a theoretical transfer efficiency of $E \sim 1$ for the dye-pair Cy3B-CF660R ($R_0 = 6$ nm), in accord with the high transfer efficiency we observe (a). (d) The rG19 structure requires high concentrations of guanidine hydrochloride (GdnHCl) to be unfolded, highlighting the high stability of this structure. Denaturation of 50 pM donor/acceptor-labeled rG19 was performed in 10 mM HEPES pH 8, 0.01 % Tween-20, 5 mM DTT. Experiments were performed in untreated µ-slides (Ibidi), starting at 7 M GdnHCl (ThermoScientific) and then sequentially diluting the denaturant concentration.

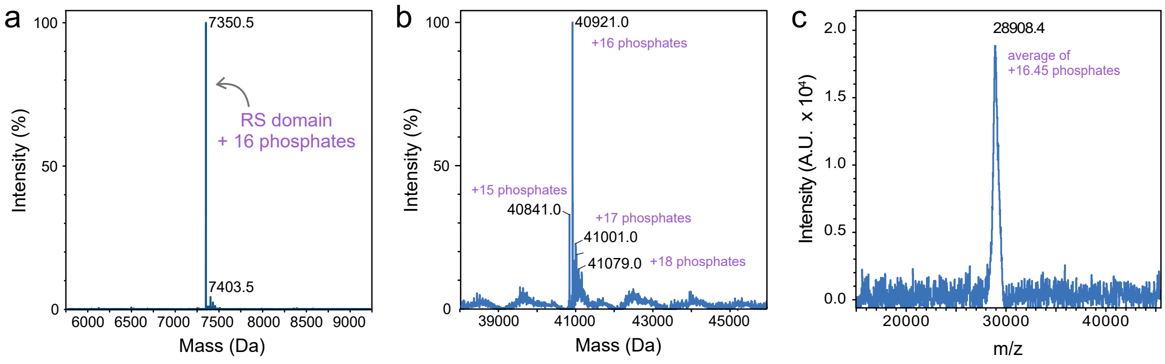


FIGURE S6. Mass spectrometry (MS) data of phosphorylated proteins. (a) Electrospray ionisation (ESI) MS of RS domain phosphorylated by SRPK1 corresponding to a species with 16 added phosphates (*cf.* calculated mass for unphosphorylated RS domain: 6071.59 Da). (b) ESI-MS of SUMO-SRSF1 phosphorylated by SRPK1 (*cf.* calculated mass for unphosphorylated SUMO-SRSF1 with N-terminal Met cleaved: 39 641.80 Da). Annotations highlight a major species of 16 added phosphates and several minor species with 15, 17 and 18 added phosphates. (c) MALDI-MS of SRSF1 phosphorylated by SRPK1 yielding on average of 16 phosphates added (*cf.* calculated mass for unphosphorylated SRSF1: 27 592.38 Da). Here, the apparent non-integer number of added phosphates can be due to lower accuracy and signal broadening typical of MALDI and/or sample heterogeneity. For all calculations, we assumed a mass increase of 79.98 Da per phosphorylation. Mass spectroscopy was performed by the Functional Genomics Center Zurich.

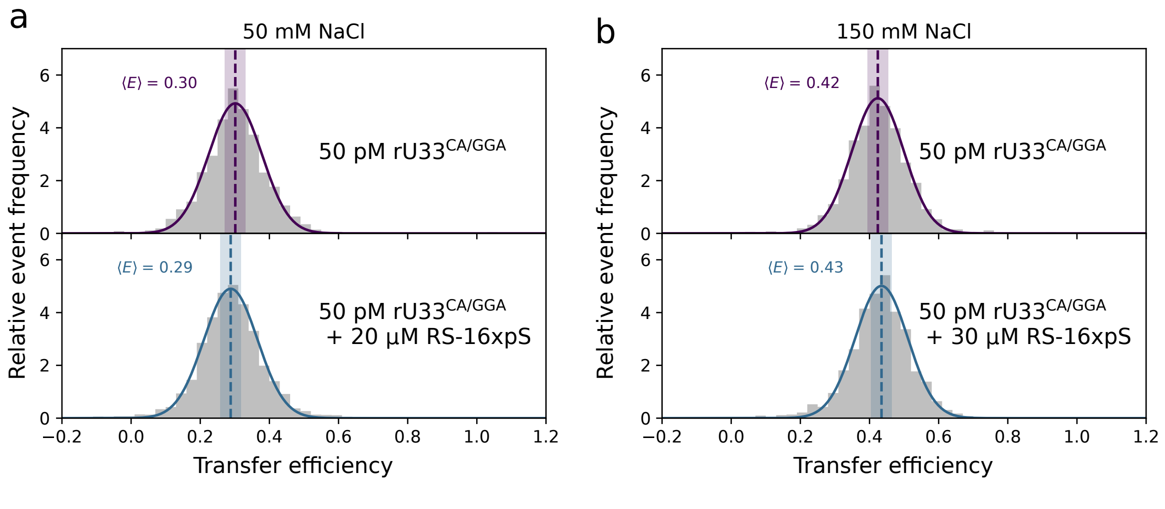


FIGURE S7. Probing possible interactions between RNA and the phosphorylated RS domain. Measurements performed using donor/acceptor-labeled $rU33^{CA/GGA}$ in standard measurement buffer but containing either (a) 50 mM NaCl or (b) 150 mM NaCl. Shown are normalised transfer efficiency histograms of RNA alone (top) and RNA with enzymatically phosphorylated RS domain (bottom). Even at reduced ionic strength, we cannot detect a significant change in RNA end-to-end distance upon the addition of phosphorylated RS domain, indicating that there is no binding at these protein concentrations.

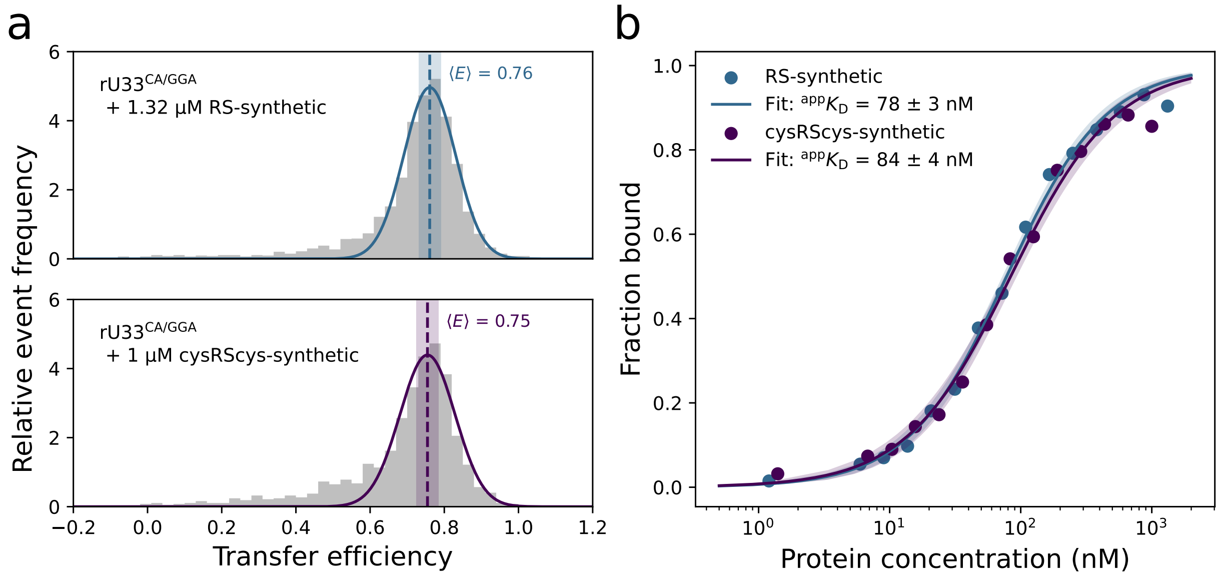


FIGURE S8. Cysteine residues have a negligible effect on the RNA affinity of synthetic WT RS domain peptides. (a) Representative normalised transfer efficiency histograms of donor/acceptor-labeled $rU33^{CA/GGA}$ RNA saturated with synthesised RS domain variants measured in 20 mM HEPES pH 8, 300 mM NaCl, 0.01 % Tween-20, 0.2 mg/mL BSA, 10 mM DTT, 0.1 U/µL RNAse inhibitor. The mean of the fit with a Gaussian peak function is indicated by a dashed line, the shaded band represents an uncertainty of 0.03 [7]. (b) Corresponding titration curves for each peptide showing the fraction bound as a function of protein concentration. Apparent affinities resulting from the fit to Equation (2) (given as best-fit value and fit error) are shown in the legend. Shaded areas around the respective fits indicate 95% confidence bands.

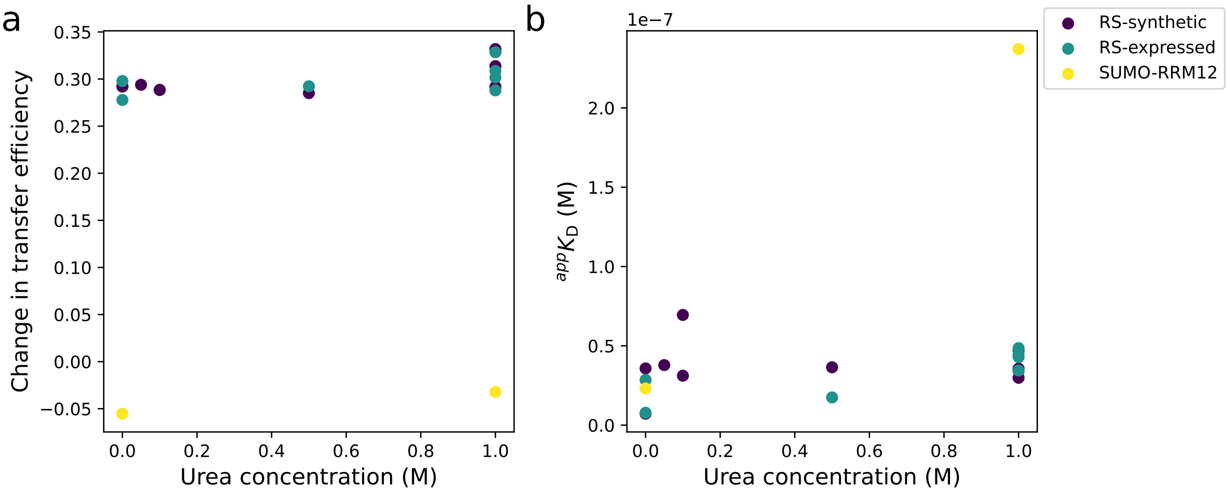


FIGURE S9. Assessing the impact of urea on RNA-protein interactions. Donor/acceptor-labeled $rU33^{CA/GGA}$ RNA was titrated with synthetically produced and with expressed wild-type RS domain at different urea concentrations to obtain binding curves. For comparison, we also include data for SUMO-RRM12 at 0 M and 1 M urea. The data shown are values obtained for individual titrations. (a) The change in the mean transfer efficiency between bound and unbound RNA as a function of urea concentration. The change in transfer efficiency upon protein binding remains similar with increasing urea concentration (< 10% change on average between 0 M and 1 M urea). (b) The apparent affinity between RS domain and RNA changes only moderately with increasing urea concentration (∼2-fold change on average between 0 M and 1 M urea). The scatter of the data at <0.2 M urea indicates that experimental variability (e.g. introduced by aggregation and burst loss) contributes substantially to the uncertainty of the apparent affinity obtained under (near-)native conditions. Compared to the RRM12-RNA affinity, which displays a decrease by almost one order of magnitude between 0 M and 1 M urea, the urea dependence observed for RS domain-RNA interactions is small.

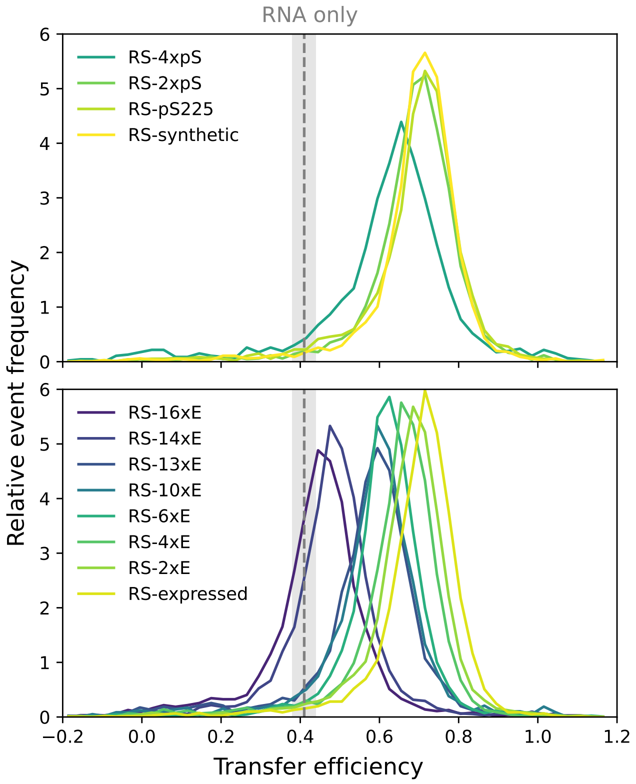


FIGURE S10. Representative normalised transfer efficiency histograms of donor/acceptor-labeled rU33$^{\mathrm{CA/GGA}}$ RNA saturated with different RS domain variants, one for each net charge, separated into phosphorylated (top) and phosphomimetic (bottom) peptides, measured in 1 M urea. The transfer efficiency of the unbound RNA is indicated with a gray dashed line and a shaded band representing an uncertainty of 0.03 [7]. For clarity, the histogram of only one of the peptides with a single phosphoserine is shown.

### II. SUPPLEMENTARY TABLES

TABLE S1. Synthetic single-stranded RNAs with 5'- and 3'-end, and internal modifications for site-specific labeling. Bases binding specifically to the RRMs of SRSF1 are highlighted in bold. Modification nomenclature: 5ThioMC6-D – 5'-end modification bearing a di-thiol after a 6-carbon linker; 3AmMO – 3'-end modification bearing an amino group; iAmMC6T – internal modification of a thymine base with a 6-carbon linker and an amino group; 3Bio - 3'-end biotinylation.

| Identifier | Sequence |
|---|---|
| rU9$^{CA/GGA}$ | /5ThioMC6-D/U**CA**UU**GGA**U/3AmMO/ |
| rU33$^{CA/GGA}$ | /5ThioMC6-D/UUUUUUUUUUUUU**CA**UU**GGA**UUUUUUUUUUUUU/3AmMO/ |
| rU59$^{CA/GGA}$ | /5ThioMC6-D/UUUUU**CA**UU**GGA**UUUUUUUUUUUUUUUUUUUUUUUUUUUUUUUUUUU UUUUUUUUU/iAmMC6T/UUUU/3Bio |
| rU33 | /5ThioMC6-D/UUUUUUUUUUUUUUUUUUUUUUUUUUUUUUUUU/3AmMO/ |
| rA19 | /5ThioMC6-D/AAAAAAAAAAAAAAAAAAA/3AmMO/ |
| rC19 | /5ThioMC6-D/CCCCCCCCCCCCCCCCCCC/3AmMO/ |
| rG19 | /5ThioMC6-D/GGGGGGGGGGGGGGGGGGG/3AmMO/ |
| rU19 | /5ThioMC6-D/UUUUUUUUUUUUUUUUUUU/3AmMO/ |

TABLE S2. List of affinities obtained for each construct. Errors shown are standard deviations for values with $N > 1$ measurements, and a factor of two for those values with only one measurement available.

| | Protein/peptide | RNA | Number of measurements | $^{\mathrm{app}}K_{\mathrm{D}}$ / M |
|---|---|---|---|---|
| native buffer | SUMO-RRM12 | $rU9^{CA/GGA}$ | 2 | $(2 \pm 1) \times 10^{-8}$ |
| | | $rU33^{CA/GGA}$ | 3 | $(2 \pm 1) \times 10^{-8}$ |
| | | rU33 | 1 | no binding detected |
| | RRM12 | $rU33^{CA/GGA}$ | 2 | $(9 \pm 6) \times 10^{-9}$ |
| | | rU33 | 1 | no binding detected |
| | SUMO-RS | $rU9^{CA/GGA}$ | 1 | $> 3 \times 10^{-5}$ |
| | | $rU33^{CA/GGA}$ | 2 | $(1.5 \pm 0.4) \times 10^{-8}$ |
| | | rU33 | 1 | $3^{+3}_{-1.5} \times 10^{-8}$ |
| | RS | $rU33^{CA/GGA}$ | 2 | $(2 \pm 1) \times 10^{-8}$ |
| | | rU33 | 1 | $(2.4 \pm 0.1) \times 10^{-8}$ |
| | | rA19 | 1 | $1^{+1}_{-0.5} \times 10^{-6}$ |
| | | rC19 | 1 | $1.2^{+1.2}_{-0.6} \times 10^{-6}$ |
| | | rG19 | 1 | could not be determined[a] |
| | | rU19 | 1 | $4^{+4}_{-2} \times 10^{-7}$ |
| | | $rU59^{CA/GGA}$ | n.a. | $(1.7 \pm 0.4) \times 10^{-9}$ [b] |
| | RS-synthetic | $rU33^{CA/GGA}$ | 2 | $(2 \pm 1) \times 10^{-8}$ |
| | SUMO-SRSF1 | $rU9^{CA/GGA}$ | 1 | $(1.7 \pm 0.1) \times 10^{-8}$ |
| | | $rU33^{CA/GGA}$ | 1 | $(5^{+5}_{-2.5}) \times 10^{-10}$ |
| | | rU33 | 2 | $(5 \pm 2) \times 10^{-9}$ |
| | SRSF1 | $rU33^{CA/GGA}$ | 2 | $(6 \pm 2) \times 10^{-10}$ |
| | | rU33 | 1 | $(6^{+6}_{-3}) \times 10^{-9}$ |
| | SUMO-pSRSF1 | $rU9^{CA/GGA}$ | 1 | $(7^{+7}_{-3.5}) \times 10^{-8}$ |
| | | $rU33^{CA/GGA}$ | 4 | $(7.0 \pm 0.8) \times 10^{-8}$ |
| | pSRSF1 | $rU33^{CA/GGA}$ | 1 | $(6.5^{+6.5}_{-3.3}) \times 10^{-8}$ |
| 1 M urea buffer | SUMO-RRM12 | $rU33^{CA/GGA}$ | 1 | $(2.4^{+2.4}_{-1.2}) \times 10^{-7}$ |
| | RS | $rU33^{CA/GGA}$ | 4 | $(4.3 \pm 0.5) \times 10^{-8}$ |
| | RS2×E | $rU33^{CA/GGA}$ | 3 | $(5.8 \pm 0.5) \times 10^{-8}$ |
| | RS4×E | $rU33^{CA/GGA}$ | 3 | $(1.5 \pm 0.2) \times 10^{-7}$ |
| | RS6×E | $rU33^{CA/GGA}$ | 5 | $(8 \pm 2) \times 10^{-7}$ |
| | RS10×E | $rU33^{CA/GGA}$ | 4 | $(1.7 \pm 0.3) \times 10^{-5}$ |
| | RS13×E | $rU33^{CA/GGA}$ | 4 | $(3.08 \pm 0.08) \times 10^{-5}$ |
| | RS14×E | $rU33^{CA/GGA}$ | 3 | $(1.3 \pm 0.3) \times 10^{-4}$ |
| | RS16×E | $rU33^{CA/GGA}$ | 3 | $(2.7 \pm 0.3) \times 10^{-4}$ |
| | RS-synthetic | $rU33^{CA/GGA}$ | 4 | $(3.7 \pm 0.7) \times 10^{-8}$ |
| | $RS\text{-}pS_{199}$ | $rU33^{CA/GGA}$ | 3 | $(6 \pm 1) \times 10^{-8}$ |
| | $RS\text{-}pS_{223}$ | $rU33^{CA/GGA}$ | 4 | $(9.0 \pm 0.8) \times 10^{-8}$ |
| | $RS\text{-}pS_{225}$ | $rU33^{CA/GGA}$ | 4 | $(1.3 \pm 0.2) \times 10^{-7}$ |
| | RS-2×pS | $rU33^{CA/GGA}$ | 4 | $(1.1 \pm 0.1) \times 10^{-7}$ |
| | RS-4×pS | $rU33^{CA/GGA}$ | 5 | $(9 \pm 1) \times 10^{-7}$ |

[a] Due to aggregation and burst loss, see Figure S5 for details.

[b] Value for the first binding event obtained from surface experiments, for error propagation, see Methods section.

TABLE S3. Overview of RS domain peptide synthesis data. Purities are based on peak area of absorbances at 214 nm in UHPLC traces. Reported yields of crude peptides are relative to resin loading. Purities of all peptides after purification were >95 %. Molecular weights are calculated from monoisotopic masses of individual residues. See Supplementary Experimental Methods and Appendix A for more details. MW – molecular weight, n.d. – not determined.

| Peptide | Crude peptide | | Pure peptide | | | |
|---|---|---|---|---|---|---|
| | mass (mg) | purity (%) | mass (mg) | yield (%) | MW (Da) | |
| | | | | | calculated | observed |
| RS-synthetic | 23 | 72 | 0.5 | 1.3 | 6067.1817 | 6067.1768 |
| cysRScys-synthetic | 8.4 | 71 | n.d. | n.d. | 6273.2001 | 6273.1808 |
| RS-pS$_{199}$ | 8.4 | 49 | 0.79 | 4.5 | 6353.1701 | 6354.1582 |
| RS-pS$_{223}$ | 6.2 | 51 | 0.9 | 5.3 | 6353.1701 | 6353.1682 |
| RS-pS$_{225}$ | 12.6 | 32 | 0.6 | 1.6 | 6353.1701 | 6353.0933 |
| RS-2×pS | 7.4 | 24 | 0.6 | 3.1 | 6433.1327 | 6433.1041 |
| RS-4×pS | 57 | 28 | 1 | 1.8 | 6593.0654 | 6593.0495 |

TABLE S4. Sequences of DNA oligonucleotides used for molecular biology.

| **Name** | **DNA sequence (5' → 3')** |
|---|---|
| SRSF1 Fwd / RRM12 Fwd | ACAGAGGACAGATTGGTGGGATGAGCGGTGGTGGTGTTATTCGTGGTCC |
| SRSF1 Rev / RS Rev | TTCGGATCCGGTCTTTATCAGGTGCGACTACGGCTGCGACTATGGC |
| RRM12 Rev | TTCGGATCCGGTCTTTATCAACCATCAACTTTCACACGAATATAGGCGGTTTCA CCTTC |
| RS Fwd | ACAGAGAACAGATTGGTGGGCGCAGTCCGAGCTATGGTCGTAGTCGTAGTC |
| pSumo Fwd | GAGCTCCGTCGACAAGCTTGCG |
| pSumo Rev | CCCACCAATCTGTTCTCTGTGAGCC |

TABLE S5. Amino acid sequences of recombinantly expressed proteins and peptides. The Ulp1 cleavage site is indicated by “/”. According to mass spectrometry, N-terminal methionines in parentheses were usually cleaved during expression in *E. coli.*

| **Name** | **Amino acid sequence** |
|---|---|
| SUMO-SRSF1 | (M)GHHHHHHGSDSEVNQEAKPEVKPEVKPETHINLKVSDGSSEIFFKIKKTTPLRRL<br>MEAFAKRQGKEMDSLRFLYDGIRIQADQTPEDLDMEDNDIIEAHREQIGG/MSGGGV<br>IRGPAGNNDCRIYVGNLPPDIRTKDIEDVFSKYGAIRDIDLKNRRGGPPFAFVEFEDP<br>RDAEDAVSGRDGYDYDGYRLRVEFPRSGRGTGRGGGGGGGGGAPRGRYGPPSRRS<br>ENRVVVSGLPPSGSWQDLKDHMREAGDVCYADVYRDGTGVVEFVRKEDMTYAVR<br>KLDNTKFRSHEGETAYIRVKVDGPRSPSYGRSRSRSRSRSRSRSRSNSRSRSYSPRRSR<br>GSPRYSPRHSRSRSRT |
| SUMO-RRM12 | (M)GHHHHHHGSDSEVNQEAKPEVKPEVKPETHINLKVSDGSSEIFFKIKKTTPLRRL<br>MEAFAKRQGKEMDSLRFLYDGIRIQADQTPEDLDMEDNDIIEAHREQIGG/MSGGGV<br>IRGPAGNNDCRIYVGNLPPDIRTKDIEDVFSKYGAIRDIDLKNRRGGPPFAFVEFEDP<br>RDAEDAVSGRDGYDYDGYRLRVEFPRSGRGTGRGGGGGGGGGAPRGRYGPPSRRS<br>ENRVVVSGLPPSGSWQDLKDHMREAGDVCYADVYRDGTGVVEFVRKEDMTYAVR<br>KLDNTKFRSHEGETAYIRVKVDG |
| SUMO-RS (RS-expressed) | (M)GHHHHHHGSDSEVNQEAKPEVKPEVKPETHINLKVSDGSSEIFFKIKKTTPLRRL<br>MEAFAKRQGKEMDSLRFLYDGIRIQADQTPEDLDMEDNDIIEAHREQIGG/RSPSYGR<br>SRSRSRSRSRSRSRSNSRSRSYSPRRSRGSPRYSPRHSRSRSRT |
| RS-2×E | (M)GHHHHHHGSDSEVNQEAKPEVKPEVKPETHINLKVSDGSSEIFFKIKKTTPLRRL<br>MEAFAKRQGKEMDSLRFLYDGIRIQADQTPEDLDMEDNDIIEAHREQIGG/RSPSYGR<br>SRSRSRSRSRSRSRSNSREREYSPRRSRGSPRYSPRHSRSRSRT |
| RS-4×E | (M)GHHHHHHGSDSEVNQEAKPEVKPEVKPETHINLKVSDGSSEIFFKIKKTTPLRRL<br>MEAFAKRQGKEMDSLRFLYDGIRIQADQTPEDLDMEDNDIIEAHREQIGG/REPSYG<br>RSRSRERSRSRSRERSNSRSREYSPRRSRGSPRYSPRHSRSRSRT |
| RS-6×E | (M)GHHHHHHGSDSEVNQEAKPEVKPEVKPETHINLKVSDGSSEIFFKIKKTTPLRRL<br>MEAFAKRQGKEMDSLRFLYDGIRIQADQTPEDLDMEDNDIIEAHREQIGG/REPSYG<br>RSRSRERSRSRSRERSNSRSREYSPRRERGSPRYEPRHSRSRSRT |
| RS-10×E | (M)GHHHHHHGSDSEVNQEAKPEVKPEVKPETHINLKVSDGSSEIFFKIKKTTPLRRL<br>MEAFAKRQGKEMDSLRFLYDGIRIQADQTPEDLDMEDNDIIEAHREQIGG/RSPSYGR<br>SRERERERERERERENEREREYSPRRSRGSPRYSPRHSRSRSRT |
| RS-13×E | (M)GHHHHHHGSDSEVNQEAKPEVKPEVKPETHINLKVSDGSSEIFFKIKKTTPLRRL<br>MEAFAKRQGKEMDSLRFLYDGIRIQADQTPEDLDMEDNDIIEAHREQIGG/REPEYG<br>RERERERERERERERENEREREYSPRRSRGSPRYSPRHSRSRSRT |
| RS-14×E | (M)GHHHHHHGSDSEVNQEAKPEVKPEVKPETHINLKVSDGSSEIFFKIKKTTPLRRL<br>MEAFAKRQGKEMDSLRFLYDGIRIQADQTPEDLDMEDNDIIEAHREQIGG/RSPSYGR<br>ERERERERERERERENEREREYSPRRSRGSPRYSPRHERERERT |
| RS-16×E | (M)GHHHHHHGSDSEVNQEAKPEVKPEVKPETHINLKVSDGSSEIFFKIKKTTPLRRL<br>MEAFAKRQGKEMDSLRFLYDGIRIQADQTPEDLDMEDNDIIEAHREQIGG/REPEYG<br>RERERERERERERERENEREREYSPRRSRGSPRYSPRHERERERT |
| Ulp1 | MLVPELNEKDDDQVQKALASRENTQLMNRDNIEITVRDFKTLAPRRWLNDTIIEFFM<br>KYIEKSTPNTVAFNSFFYTNLSERGYQGVRRWMKRKKTQIDKLDKIFTPINLNQSHW<br>ALGIIDLKKKTIGYVDSLSNGPNAMSFAILTDLQKYVMEESKHTIGEDFDLIHLDCPQQ<br>PNGYDCGIYVCMNTLYGSADAPLDFDYKDAIRMRRFIAHLILTDALKLEHHHHHH |
| SRPK1 | MQYKLILNGKTLKGETTTEAVDAATAEKVFKQYANDNGVDGEWTYDDATKTFTV<br>TEGSHHHHHHMERKVLALQARKKRTKAKKDKAQRKSETQHRGSAPHSESDLPEQEE<br>EILGSDDDEQEDPNDYCKGGYHLVKIGDLFNGRYHVIRKLGWGHFSTVWLSWDIQG<br>KKFVAMKVVKSAEHYTETALDEIRLLKSVRNSDPNDPNREMVVQLLDDFKISGVNGT<br>HICMVFEVLGHHLLKWIIKSNYQGLPLPCVKKIIQQVLQGLDYLHTKCRIIHTDIKPEN<br>ILLSVNEQYIRRLAAEATEWQRSGAPPPSGSAVSTAPQPKPADKMSKNKKKKLKKKQ<br>KRQAELLEKRMQEIEEMEKESGPGQKRPNKQEESESPVERPLKENPPNKMTQEKLE<br>ESSTIGQDQTLMERDTEGGAAEINCNGVIEVINYTQNSNNETLRHKEDLHNANDCDV<br>QNLNQESSFLSSQNGDSSTSQETDSCTPITSEVSDTMVCQSSSTVGQSFSEQHISQLQES<br>IRAEIPCEDEQEQEHNGPLDNKGKSTAGNFLVNPLEPKNAEKLKVKIADLGNACWVH<br>KHFTEDIQTRQYRSLEVLIGSGYNTPADIWSTACMAFELATGDYLFEPHSGEEYTRD<br>EDHIALIIELLGKVPRKLIVAGKYSKEFFTKKGDLKHITKLKPWGLFEVLVEKYEWSQ<br>EEAAGFTDFLLPMLELIPEKRATAAECLRHPWLNSVEHHHHHH |

TABLE S6. Amino acid sequences of synthetic peptides. Note that the C-terminal functional group is an amide ($CONH_2$).

| **Name** | **Amino acid sequence** |
|---|---|
| RS-synthetic | RSPSYGRSRSRSRSRSRSRSRSNSRSRSYSPRRSRGSPRYSPRHSRSRSRT |
| cysRScys-synthetic | CRSPSYGRSRSRSRSRSRSRSRSNSRSRSYSPRRSRGSPRYSPRHSRSRSRTC |
| RS-p$S_{199}$ | CRpSPSYGRSRSRSRSRSRSRSRSNSRSRSYSPRRSRGSPRYSPRHSRSRSRTC |
| RS-p$S_{223}$ | CRSPSYGRSRSRSRSRSRSRSRSNSRpSRSYSPRRSRGSPRYSPRHSRSRSRTC |
| RS-p$S_{225}$ | CRSPSYGRSRSRSRSRSRSRSRSNSRSRpSYSPRRSRGSPRYSPRHSRSRSRTC |
| RS-2×pS | CRSPSYGRSRSRSRSRSRSRSRSNSRpSRpSYSPRRSRGSPRYSPRHSRSRSRTC |
| RS-4×pS | CRpSPSYGRSRSRpSRSRSRSRpSRSNSRSRpSYSPRRSRGSPRYSPRHSRSRSRTC |

### III. SUPPLEMENTARY EXPERIMENTAL METHODS

#### A. Preparation of recombinantly expressed proteins and peptides

##### 1. *SRSF1*

The plasmid bearing $His_6$-SUMO-SRSF1 was transformed into chemically competent *E. Coli* BL21(DE3)-C41 cells (Lucigen). Cultures were grown in 2xYT supplemented with 50 µg/mL kanamycin at 37 °C until an OD600 of $\sim 1.5$ was reached, and protein expression was induced using 0.5 mM IPTG at 20 °C overnight. Cells were harvested, resuspended in lysis buffer (20 mM sodium phosphate (NaP) pH 7.5, 800 mM NaCl, 5 % glycerol, 0.01 % Tween-20, 2 mM DTT, 150 mM L-arginine, 150 mM L-glutamate) supplemented with 10 mM MgCl2, 10 U/mL benzonase (Merck) and 20 µg/mL RNase A (NEB), and lysed by passing the suspension three times through a high pressure homogeniser (HPL6, Maximator) cooled to 4 °C at 15,000-20,000 psi. The lysate was clarified by centrifugation at 35 000 ×g for 30 min at 4 °C and applied to a HisTrap Excel column (Cytiva, 5 ml per 1 l cell culture) equilibrated in lysis buffer. The column was washed with 15 column volumes (CVs) of lysis buffer, followed by 7 CVs of high-salt wash buffer (20 mM NaP pH 7.5, 3 M NaCl, 5 % glycerol, 0.01 % Tween-20, 2 mM DTT, 100 mM L-arginine, 100 mM L-glutamate) and 7 CVs of lysis buffer with 15 mM imidazole, before elution using 20 mM NaP pH 7.5, 800 mM NaCl, 5 % glycerol, 0.01 % Tween-20, 2 mM DTT, 150 mM L-arginine, 150 mM L-glutamate, 500 mM imidazole. Fractions containing protein were pooled and diluted at least 8-fold using 20 mM HEPES pH 8.0, 10 % glycerol, 0.01 % Tween-20, 0.5 mM TCEP and 5 M NaCl until the sample was clear (0.6 to 0.8 M ionic strength). Nucleic acid and protein contaminants were removed by ion exchange chromatography (IEX). After filtering (0.22 µm syringe filter, PES membrane, Techno Plastic Products AG), the diluted protein solution was applied to a MonoS 5/50 GL (Cytiva) equilibrated in 20 mM HEPES pH 8.0, 600 mM NaCl, 10 % glycerol, 0.01 % Tween-20, 0.5 mM TCEP. The column was then washed with 10 CV of the equilibration condition and a gradient of 0.6 to 1.5 M NaCl was used to elute $His_6$-SUMO-SRSF1. Fractions with absorbance ratios of $A260/A280 < 0.7$ were pooled, concentrated by centrifugation (Vivaspin 6, 10 kDa MWCO, PES, Cytiva, and Amicon Ultra 0.5, 10 kDa MWCO, cellulose, Merck Millipore), flash-frozen in liquid $N_2$ and stored at −80 °C. To obtain untagged SRSF1, $His_6$-SUMO-SRSF1 was diluted in of 20 mM HEPES pH 8, 150 mM NaCl, 5 % glycerol, 0.5 mM TCEP, 0.01 % Tween-20 and incubated with 100 µg ULP1 for 2 h at 30 °C. Cleaved constructs were again purified by IEX, as described above.

##### 2. *RRM12*

The plasmid bearing $His_6$-SUMO-RRM12 was transformed into *E. Coli* BL21(DE3)-C41 cells. An overnight culture of this transformation was used to inoculate 2xYT medium containing 50 µg/mL kanamycin, which was then grown at 37 °C until an OD600 of $\sim 0.8$ was reached. The culture was left to cool to 20 °C and induced with 1 mM IPTG overnight. Cells were harvested at 4000 ×g and the pellet was resuspended in lysis buffer (20 mM NaP pH 7.4, 800 mM NaCl, 10 mM imidazole, 1 % TritonX) with subsequent flash-freezing. After thawing, the cells were lysed using a Maximator high-pressure homogenizer HPL6 (Maximator) at 17 to 21 MPa for 3 iterations. The lysate was clarified by centrifugation for 25 min at 4 °C and 40 000 ×g. The soluble protein fraction was loaded onto ∼20 mL Ni Sepharose Excel resin (GE Healthcare) equilibrated in wash buffer (20 mM NaP pH 7.4, 800 mM NaCl, 20 mM imidazole, 150 mM L-Arg, 150 mM L-Glu, 6 % v/v glycerol, 0.005 % v/v Tween20, 5 mM DTT). The resin-supernatant mixture was incubated for 30 min at 4 °C on a shaker. After incubation and subsequent gravity-assisted elution of the flow through, the column was washed with 10 column volumes (CV) wash buffer. $H_6$-SUMO-RRM12 was eluted with 8 CV elution buffer (20 mM NaP pH 7.4, 800 mM NaCl, 500 mM imidazole, 100 mM L-Arg, 100 mM L-Glu, 6 % v/v glycerol, 0.005 % v/v Tween20, 5 mM DTT). IMAC elutions were combined and dialysed against dialysis buffer (20 mM $KP_i$ pH 7.4, 25 mM KCl, 10 % v/v glycerol, 0.01 % v/v Tween20) overnight at 4 °C. A HiPrep Q FF 16/10 column (GE Healthcare) was equilibrated in 20 mM $KP_i$ pH 7.4, 10 % v/v glycerol, 2 mM DTT before loading the dialysed SUMO-RRM12. After washing, the protein was eluted with a gradient of 0 to 0.44 M KCl over 20 CV. $His_6$-SUMO-RRM12 was incubated with Ulp1 (10 µg Ulp1 per 5 µg of $H_6$-

SUMO-RRM12) in cleavage buffer (20 mM HEPES pH 7, 200 mM NaCl, 2 mM DTT, 0.5 % v/v glycerol) overnight at 4 °C. The cleaved RRM12 was purified by reverse IMAC using 20 mL Ni Sepharose Excel resin (Cytavia) pre-equilibrated with wash buffer (20 mM NaP pH 7.4, 800 mM NaCl, 142 mM L-Arg, 142 mM L-Glu, 6 % v/v glycerol, 0.005 % v/v Tween20, 5 mM DTT). The resin was incubated with the cleavage reaction mixture for 40 min at 4 °C on a shaker. After collecting the flow through, the resin was washed with 3×4 CV wash buffer with subsequent elution of the $His_6$-SUMO tag, uncleaved $His_6$-SUMO-RRM12, and Ulp1 using 3×4 CV elution buffer (20 mM NaP pH 7.4, 800 mM NaCl, 500 mM imidazole, 142 mM L-Arg, 142 mM L-Glu, 6 % v/v glycerol, 0.005 % v/v Tween20, 5 mM DTT). The collected flow-through and wash were pooled and concentrated by centrifugal filtering (Vivaspin 20, 10 kDa MWCO), and dialysed into 20 mM HEPES pH 8.0, 400 mM NaCl, 5 mM DTT. Proteins were flash-frozen in liquid $N_2$ and stored at −80 °C.

#### *3. RS domain*

Chemically competent *E.coli* BL21(DE3)-C41 cells (Lucigen) were transformed with pET-$His_6$-SUMO-RS and grown in 2xYT medium with 50 µg/mL kanamycin at 37 °C. Protein expression was induced by the addition of 0.5 mM isopropyl-β-D-1-thiogalactopyranoside (IPTG) at an OD600 of ∼0.7, followed by overnight incubation at 20 °C. Purification of the RS domain was carried out using the same IMAC protocol as the one used for $His_6$-SUMO-SRSF1. Fractions containing protein were diluted to an approximate ionic strength of 0.3 M and further purified by IEX chromatography using a MonoS 5/50 GL column (GE Healthcare), equilibrated in 20 mM HEPES pH 8, 10 % Glycerol, 0.5 mM TCEP and 300 mM NaCl. After washing with the equilibration conditions, a gradient from 0.3 to 1.5 M NaCl was used to elute $His_6$-SUMO-RS. Fractions containing protein were analyzed by SDS-PAGE and subsequently, those of sufficient purity were pooled, flash-frozen in liquid $N_2$ and stored at −80 °C. To obtain untagged RS domain, 1 mL of $His_6$-SUMO-RS was diluted in 15 mL of 20 mM HEPES pH 8, 150 mM NaCl, 5 % glycerol, 0.5 mM TCEP, 0.01 % Tween-20 and incubated with 100 µg ULP1 for 2 h at 30 °C. Cleaved constructs were again purified by IEX, as described above.

#### *4. RS domain glutamate variants*

Plasmids were transformed into chemically competent *E. coli* BL21 (DE3) cells (Thermo Scientific, EC0114), and protein expression and purification was carried out similar to that of the RS domain with following modifications. Protein expression was induced at OD600∼1. All IMAC buffers were prepared without Tween-20, and the lysis buffer did not contain Benzonase, RNAse and $MgCl_2$. After loading clarified cell lysates of constructs containing 2 to 6 glutamates, the HisTrap Excel column was washed with 10 CV of lysis buffer, 5 CV of high-salt wash buffer, and 5 CV of 3 % elution buffer (lysis buffer with 15 mM imidazole). Columns loaded with constructs containing ≥10 glutamates were washed using 15 CV lysis buffer and 5 CV of 3 % elution buffer. After elution, fractions containing protein were combined and diluted ∼0.3 M ionic strength using $H_2O$ (peptides with 2 to 6 glutamates), or buffer exchanged into 20 mM NaP pH 7.5, 50 mM Glu, 50 mM Arg, 5 % glycerol, 2 mM DTT using a HiPrep 26/10 Desalting column (Pharmacia) (peptides with ≥10 glutamates). If phase separation occurred the construct was either diluted further, or 5 M NaCl was added gradually until condensates dissolved. SUMO-tag cleavage was achieved by the addition of 100 µg Ulp1 and incubation at 30 °C for ≥1 h. Precipitate was removed by filtration (0.22 µm syringe filter, PES membrane, Techno Plastic Products AG) and the cleavage reaction was loaded onto a MonoS 5/50 GL column (GE Healthcare) equilibrated in 20 mM HEPES pH 8, 5 % glycerol, 0.5 mM TCEP containing 300 mM NaCl (RS domain with 2 glutamates), 150 mM NaCl (peptides with 4 and 6 glutamates), or 50 mM (peptides with ≥10 glutamates). After a 10 CV wash using the respective equilibration conditions, elution gradients (20 CV in length) were adjusted to start at the equilibration salt concentration and end at 1.5 M NaCl (2 to 6 glutamates) or 1.2 M NaCl (≥10 glutamates). Fractions containing protein of sufficient purity as determined by SDS-PAGE were combined and concentrated by centrifugal filtration (Vivaspin 6, 3 kDa MWCO, PES, Cytiva, and Amicon Ultra 0.5 3 kDa MWCO, cellulose, Merck Millipore). All proteins but the variant containing 2 glutamates were further purified by size exclusion chromatography using a Superdex 75 Increase 10/300

GL column (Cytiva) equilibrated in 20 mM HEPES pH 8 and 1 M NaCl (4 and 6 glutamates), 0.75 M NaCl (10 and 13 glutamates) or 0.5 M NaCl (14 and 16 glutamates). Constructs containing 14 and 16 glutamates were purified twice by SEC to remove a closely eluting contaminant. Fractions containing pure protein were pooled, concentrated to maximally ∼0.9 mM (construct dependent), flash-frozen in liquid $N_2$ and stored at −80 °C. If necessary, proteins were further concentrated after thawing, directly prior to sample preparation for single-molecule experiments.

#### 5. *SRPK1*

GB1-$His_6$-SRPK1-$His_6$ was transformed into *E. coli* BL21-CodonPlus(DE3)-RIL cells and incubated in LB media supplemented with 50 µg/mL kanamycin and 100 µg/mL chloramphenicol overnight at 37 °C. The overnight culture was used to inoculate 6 L of media with antibiotics. At an OD600 ∼ 0.6 protein expression was induced by the addition of 0.5 mM IPTG and incubation for 4 h at 37 °C. Cells were harvested by centrifugation at 4000 ×g, 4 °C and resuspended in 10 mL lysis buffer (20 mM NaP pH 7.4, 800 mM NaCl, cOMPLETE EDTA-free protease inhibitor cocktail per 1 g cell pellet). Cells were lysed using a microfluidizer (M-110L, Microfluidics) using three passes at 15000 psi. The lysate was clarified by centrifugation at 17 000 ×g, 4 °C. The supernatant was applied to 6 ml Ni-NTA sepharose CL-6B (Qiagen) equilibrated in lysis buffer. The matrix was then washed with 2 CV of 3 M NaCl, followed by 2 CV of wash buffer (20 mM NaP pH 7.4, 800 mM NaCl, 20 mM imidazole, 3 mM DTT, 150 mM L-arginine, 150 mM L-glutamate). Protein was eluted with 4 to 5 CV of elution buffer (20 mM NaP pH 7.4, 800 mM NaCl, 500 mM imidazole, 3 mM DTT, 150 mM L-arginine, 150 mM L-glutamate). The protein was dialysed against 2 L of 20 mM NaP pH 7.4, 800 mM NaCl, 3 mM DTT, 150 mM L-arginine, 150 mM L-glutamate using a 3.5 kDa MWCO regenerated cellulose membrane (Spectra/Por®,Spectrumlabs), incubating at 4 °C overnight. For phosphorylation reactions of proteins that would later be used in single-molecule experiments, the protein was further purified by IEX using a MonoS 5/50 GL column (GE Healthcare) equilibrated in 20 mM HEPES pH 8, 50 mM NaCl, 0.5 mM TCEP. The protein was eluted with a gradient of 0.05 to 1 M NaCl over 15 CV. Fractions containing protein were pooled, flash-frozen in liquid $N_2$ and stored at −80 °C.

#### 6. *Ulp1*

The plasmid containing Ulp1-$His_6$ was transformed into *E. coli* BL21(DE3) and stored as a glycerol stock at −80 °C. For expression, 100 mL of 2xYT supplemented with 50 µg kanamycin were inoculated and incubated at 37 °C overnight. The overnight culture was diluted into 2 L of 2xYT and incubated until $OD600$ ∼ 0.6. Protein expression was induced with 1 mM IPTG for ∼4 h at 30 °C. After harvesting at 4000 ×g, 4 °C, cell pellets were resuspended in wash buffer (50 mM Tris-HCl pH 8.0, 350 mM NaCl, 10 mM imidazole, 2 mM DTT, 2% glycerol) and lysed mechanically sing a high pressure homogenizer (HPL6, Maximator). The lysate was clarified by centrifugation at 35 000 ×g, 4 celsius and the supernatant was applied to a 5 mL HisTrap Excel column (GE Healthcare). The column was washed with 10 CV wash buffer, 5 CV wash buffer containing 1 M NaCl and again 10 CV of normal wash buffer. The protein was eluted using 50 mM Tris-HCl pH 8.0, 350 mM NaCl, 400 mM imidazole, 2 mM DTT, 2% glycerol. Fractions containing sufficiently pure protein were pooled and buffer exchanged into 50 mM Tris-HCl pH 8.0, 400 mM NaCl, 2 mM DTT, 2% glycerol using a HiPrep 26/10 Desalting column (Pharmacia). Fractions containing protein were combined and concentrated to ∼4 mg/mL by centrifugation (Vivaspin 20, 10 kDa MWCO, PES). The protein was diluted to ∼2 mg/mL by the addition of 100% glycerol, flash-frozen in liquid $N_2$ and stored at −80 °C.

#### B. Peptide synthesis and purification

##### 1. Reagents and solvents

Fmoc- and side chain-protected L-amino acids (Fmoc-Arg(Pbf)-OH, Fmoc-Asn(Trt)-OH, Fmoc-Cys(Trt)-OH, Fmoc-Gly-OH, Fmoc-His(Trt)-OH, Fmoc-Pro-OH, Fmoc-Ser(*t*-Bu)-OH, Fmoc-Thr(*t*-Bu)-OH, Fmoc-Tyr(*t*-Bu)-OH) were purchased from the Novabiochem-line from Sigma-Aldrich Canada Ltd. Fmoc-L-Ser(PO(OBzl)OH)-OH was purchased from Iris Biotech GmbH. HATU (O-(7-azabenzotriazol-1-yl)-N,N,N',N'-tetramethyluronium hexafluorophosphate) and PyAOP (7-azabenzotriazol-1-yloxy)tripyrrolidinophosphonium hexafluorophosphate) were purchased from Advanced ChemTech CreoSalus. N,N-diisopropylethylamine ($iPr_2NEt$, DIPEA, 99.5 %), trifluoroacetic acid (TFA, for HPLC, ≥99.0 %), triisopropylsilane (TIPS, 98 %), 3,6-dioxa-1,8-octane-dithiol (DODT, 95 %), acetonitrile (MeCN, for HPLC gradient grade, ≥99.9 %) were purchased from Sigma-Aldrich. N,N-Dimethylformamide (DMF) was purchased from the Supelco-line of Sigma-Aldrich Canada Ltd. Dichloromethane (DCM, ≥99.8 %) was purchased from Fisher Scientific Ltd. Diethyl ether was purchased from Honeywell Riedel-de Haën. NovaPEG Rink Amide resin (loading 0.20 mmol/g) was purchased from the Novabiochem-line of Sigma-Aldrich Canada Ltd.

##### 2. Automated fast-flow peptide synthesis (AFPS)

Peptides were synthesized on an automated-flow system built in the Hartrampf lab that is similar to a previously published AFPS system [8]. The following settings were used for peptide synthesis: a flow rate of 20 mL/min for coupling and deprotection steps, the reactor was kept at 90 °C, the heating loop at 90 °C or 30 °C (specified below). In each synthetic cycle the resin is pre-washed at 90 °C for 60 s at 40 mL/min. During the coupling step, three HPLC pumps are used: a 50 mL/min pump head pumps the activating agent, a second 50 mL/min pump head pumps the amino acid, and a 5.0 mL/min pump head pumps DIPEA (*neat*). The 50 mL/min pump head delivered 0.398 679 mL of liquid per pump stroke, the 5.0 mL/min pump head pumps 0.039 239 mL of liquid per pump stroke.

All peptides were prepared by AFPS and were synthesized on commercially available Novabiochem® NovaPEG Rink Amide resin (0.20 mmol/g, exact amounts of resin used for each peptide are listed in Section Section III B 7) and standard Fmoc/*t*-Bu protected amino acids (0.40 M in DMF, 0.20 M final concentration) were coupled using HATU (0.38 M in DMF, 0.19 M final concentration) or PyAOP (0.38 M in DMF, 0.19 M final concentration) with DIPEA (delivered *neat*, approx. 0.27 M final concentration) at a total flow rate of 20 mL/min. Amino acids G, P, S, phosphoserine (pS), and Y were coupled using HATU. Amino acids C, H, N, R and T were coupled using PyAOP. For amino acids G, P, S, and Y, a total volume of 6.4 mL of the "coupling solution" (i.e. containing amino acid at 0.20 M, HATU or PyAOP at 0.19 M and DIPEA at 0.27 M final concentration in DMF) was applied for each coupling. For amino acids C, H, N, R, S and T, a total of 10.4 mL of "coupling solution" was applied for each coupling. For pS, a total of 2.4 mL "coupling solution" was applied for each coupling unless stated otherwise. Removal of the $N^{\alpha}$-Fmoc group was achieved using 20 % piperidine with 1 % formic acid in DMF (total volume 6.4 mL) at a flow rate of 20 mL/min. The deprotection solution was pre-heated at 30 °C for C, H and pS, and 90 °C for all other Fmoc-protected amino acids. Between each coupling and deprotection step, the resin was washed with DMF (32 mL) at a flow rate of 40 mL/min, preheated to 30 °C for amino acids C, H and pS, and at 90 °C for all others. Total synthesis time to afford resin-bound RS domain peptides was approximately 2.5 h. After completion of the peptide sequence, resins were manually washed with DCM (3 × 5 mL) and dried under reduced pressure.

##### 3. Peptide cleavage and deprotection

Exact amounts of resin taken forward for cleavage are listed in Section Section III B 7. Peptides were cleaved using a solution of TFA/TIPS/DODT/$H_2O$ (94:1:2.5:2.5, v/v/v/v, 1 to 3 mL) for 2 h at 23 °C with gentle mixing. TFA was removed by evaporation under a light stream of $N_2$, and peptides were precipitated and isolated by centrifugation from ice-cold diethyl ether using 2 × 15 mL. The resulting peptide pellets were briefly dried under a light stream of $N_2$, dissolved in an aqueous solution containing 10 to 50 % MeCN and 0.1 % TFA, and lyophilized. Crude peptides were analyzed for purity by LCMS and UHPLC.

##### *4. Analytical Ultra-High Performance Liquid Chromatography (UHPLC)*

Filtered peptide solutions were diluted in 500 µL of 10 to 50 % acetonitrile (MeCN) in water with 0.1 % TFA to a final concentration of approximately 0.5 mg/mL. To achieve the best possible resolution for each peptide, the samples were applied to either an Agilent ZORBAX RRHD Eclipse Plus C8 column (2.1 mm×50 mm, 1.8 µm particle size, hereafter referred to as column A), an Agilent ZORBAX RRHD 300Å StableBond C18 column (2.1 mm×100 mm, 1.8 µm particle size, hereafter referred to as column B), or an Agilent ZORBAX StableBond 300 C18 column (2.1 mm×150 mm, 5 µm particle size, hereafter referred to as column C), connected to an UHPLC system of the Agilent 1290 Infinity II Series controlled by Agilent OpenLab CDS and ChemStation software. The column was kept at 40 °C, the flow rate was 0.80 mL/min, and peptide absorbance was detected at 214 nm. A binary solvent system consisting of Solvent A (5 % MeCN in 95 % water with 0.1 % TFA) and Solvent B (95 % MeCN containing 5 % water and 0.1 % TFA) was used. After sample application, the column was washed with 100 % Solvent A for 1.5 min before peptides were eluted with a linear gradient corresponding to 5 to 95 % MeCN or 0 to 100 % MeCN over varying lengths of time (exact gradients are listed in the corresponding figure legends in Section Section III B 7). At the end of the gradient, the column was first washed in 100 % Solvent B at 0.3 mL/min for 2 min before re-equilibration in 100 % Solvent A for 5 min. Using the ChemStation software, purities of crude and purified peptides were determined by integration of all UV signals at 214 nm (integration intervals are listed in the corresponding figure legends in Section Section III B 7). Purities are given in percent of the product peak relative to all other integrated absorbances.

##### *5. Liquid Chromatography with High-Resolution Electrospray Ionization Mass Spectrometry (LCMS)*

To determine peptide mass and purity by LCMS, filtered peptide solutions were diluted in either 10 to 50 % acetonitrile (MeCN) in water with 0.1 % TFA (60 to 500 µL), or in guanidine hydrochloride (6.0 M) to a final concentration of approximately 0.1 mg/mL. The samples were analyzed on an Acquity UHPLC (Waters, Milford, USA) connected to an Acquity eλ diode array detector and a Synapt G2HR-ESI-QTOF-MS (Waters, Milford, USA). For standard analysis of all peptides, samples were applied to an Acquity BEH C8 HPLC column (2.1 mm×100 mm, 1.7 µm particle size, Waters) kept at 30 °C at a flow rate of 0.4 mL/min. UV absorbances were monitored at 190 to 300 nm at 1.2 nm resolution and 20/s. The binary solvent system used for LC-MS consisted of water containing 0.02 % formic acid and 0.04 % TFA (Solvent A) and MeCN containing 0.04 % formic acid and 0.02 % TFA (Solvent B). All samples were analyzed using one of the following gradients:

- **LC-MS Gradient A:** isocratic step at 10 % Solvent B for 3 min, linear gradient of 10 to 70 % Solvent B over 9 min, isocratic step at 70 % Solvent B for 1 min
- **LC-MS Gradient B:** isocratic step at 3 % Solvent B for 3 min, linear gradient of 3 to 95 % Solvent B over 9 min, isocratic step at 95 % Solvent B for 1 min

The exact gradient used is specified in the corresponding figure legend for each peptide in Section Section III B 7. The mass spectrometer was set to positive ionization mode with a capillary voltage of 3.0 kV, sampling cone set to 40 V, extraction cone set to 4 V, $N_2$ cone gas at 4 L/h, $N_2$ desolvation gas at 800 L/min, and a source temperature set to 120 °C. The mass analyzer in resolution mode was used to resolve masses in a range of 150 to 3000 m/z with a scan rate of 1 Hz, mass calibration set to <2 ppm within 50 to 2500 m/z using an aqueous solution of 5 mM $NaHCO_2$. The lock masses used were caffeine (m/z = 195.088, 0.7 ng/mL) and leucine-enkephalin (m/z = 556.2771, 2 ng/mL). All mass spectra shown are deconvolved masses calculated from raw m/z values using Mestrelab Research S.L.© MestReNova v. 14.1 Mnova MS Suite. Purity based on LC-MS was determined by calculating the area under the curve of the desired product peak as a percentage of the integral of all peaks within 2 to 10 min of the absorbance chromatogram at 214 nm.

#### 6. *Semi-Preparative Reverse-Phase High Performance Liquid Chromatography (RP-HPLC)*

Semi-preparative RP-HPLC was performed on a Shimadzu prominence HPLC system (Shimadzu Corp., Japan) with a CBM-40 system controller module, an FRC-10A fraction collector, two LC-20AR pumps, and an SPD-40 UV/VIS detector, using either an Agilent Zorbax Eclipse XDB-C8 column (9.4 mm×250 mm, 5 µm particle size) or an Agilent Zorbax Eclipse XDB-C18 Semi-Preparative column (9.4 mm×250 mm, 5 µm particle size), kept at 23 °C, with a flow rate of 4 mL/min or 3.5 mL/min, respectively. A binary solvent system was used, wherein Solvent A was $H_2O$ containing 0.1 % TFA, and Solvent B was MeCN containing 0.1 % TFA. Purifications were executed using the gradients specified for each peptide in Section Section III B 7. Fractions that contained peptide of the correct m/z and high purity, as determined by LC-MS, were combined and lyophilized to afford RS domain peptides as white amorphous solids.

#### 7. *Peptide synthesis of RS domain variants - peptide specific details*

*a. Synthetic RS domain* The peptide corresponding to the RS domain of SRSF1 (residues 198-248) was synthesized on NovaPEG Rink Amide resin (144 mg, 29 µmol) using the standard AFPS protocol. Cleavage of the peptidyl-resin (94.2 mg, approx. 8.3 µmol) afforded the crude peptide as a colorless solid (23 mg, 79 % purity by LCMS, 72 % purity by UHPLC, Figures S11 and S12). The purification of the crude material (18 mg) was carried out by semi-preparative RP-HPLC using an Agilent Eclipse C8 column (5 µm, 9.4 mm × 250 mm) at 60 °C with a flow rate of 4 mL/min and a gradient of 0.5 %B/min over 42.5 min starting at 1 %B. Fractions identified with the correct $m/z$ and high purity by LCMS analysis (Figure S13) were combined and lyophilized to afford RS domain peptide as a white amorphous solid (0.50 mg, 1.3 % yield based on resin loading, >95 % purity by UHPLC, Figure S14).

*b. cysRScys* The peptide corresponding to the RS domain of SRSF1 (residues 198-248) with terminal cysteines was synthesized on NovaPEG Rink Amide resin (156.6 mg, 31 µmol) using the standard AFPS protocol. Cleavage of the peptidyl-resin (50.3 mg, approx. 4.7 µmol) afforded the crude peptide as a colorless solid (8.4 mg, 52 % purity by LCMS, 71 % purity by UHPLC, Figures S15 and S16). The purification of the crude material was carried out by semi-preparative RP-HPLC using an Agilent Eclipse C8 column (5 µm, 9.4 mm × 250 mm) at 60 °C with a flow rate of 4 mL/min and a gradient of 1 %B/min over 62 min starting at 1 %B. Fractions identified with the correct $m/z$ and high purity by LCMS analysis (Figure S17) were combined and lyophilized to afford cysRScys domain peptide as a white amorphous solid (>95 % purity by UHPLC, Figure S18). The yield was not determined.

*c. RS-pS$_{199}$* The peptide corresponding to the RS domain of SRSF1 (residues 198-248) with terminal cysteines and bearing a phosphorylation at Ser199 was synthesized on NovaPEG Rink Amide resin (151.1 mg, 31 µmol) using the standard AFPS protocol. Fmoc-Ser(PO(OBzl)OH)-OH was incorporated at position Ser199. Cleavage of the peptidyl-resin (33 mg, approx. 2.9 µmol) afforded the crude peptide as a colorless solid (8.4 mg, 89 % purity by LCMS, 49 % purity by UHPLC, Figures S19 and S20). The purification of the crude material (8.4 mg) was carried out by semi-preparative RP-HPLC using an Agilent Eclipse C8 column (5 µm, 9.4 mm × 250 mm) at 60 °C with a flow rate of 4 mL/min and an initial isocratic gradient of 1 %B for 5 min followed by a gradient of 0.5 %B/min over 62 min starting at 1 %B. Fractions identified with the correct $m/z$ and high purity by LCMS analysis (Figure S21) were combined and lyophilized to afford RS-pS$_{199}$ as a white amorphous solid (0.79 mg, 4.5 % yield based on resin loading, >95 % purity by UHPLC, Figure S22).

*d. RS-pS$_{223}$* The peptide corresponding to the RS domain of SRSF1 (residues 198-248) with terminal cysteines bearing a phosphorylation at Ser223 was synthesized on NovaPEG Rink Amide resin (148 mg, 30 µmol) using the standard AFPS protocol. Fmoc-Ser(PO(OBzl)OH)-OH was incorporated at position Ser223. Cleavage of the peptidyl-resin (64 mg, approx. 5.4 µmol) afforded the crude peptide as a colorless solid (6.2 mg, 59 % purity by LCMS, 51 % purity by UHPLC, Figures S23 and S24). The purification of the crude material (3.1 mg) was carried out by semi-preparative RP-HPLC using an Agilent Eclipse C8 column (5 µm, 9.4 mm × 250 mm) at 60 °C with a flow rate of 4 mL/min and an initial isocratic gradient of 1 %B for 5 min followed by a gradient of 0.5 %B/min over 60 min starting at

1 %B. Fractions identified with the correct $m/z$ and high purity by LCMS analysis (Figure S25) were combined and lyophilized to afford RS-pS$_{223}$ as a white amorphous solid (0.90 mg, 5.3 % yield based on resin loading, 95 % purity by UHPLC, Figure S26).

*e. RS-pS$_{225}$* The peptide corresponding to the RS domain of SRSF1 (residues 198-248) with terminal cysteines bearing a phosphorylation at Ser225 was synthesized on NovaPEG Rink Amide resin (162.8 mg, 33 µmol) using the standard AFPS protocol. Fmoc-Ser(PO(OBzl)OH)-OH was incorporated at position Ser225. Cleavage of the peptidyl-resin (approx. 70 mg, approx. 5.4 µmol) afforded the crude peptide as a colorless solid (12.7 mg, mass determined by LCMS, 32 % purity by UHPLC, Figures S27 and S28). The purification of the crude material (12.6 mg) was carried out by semi-preparative RP-HPLC using an Agilent Eclipse C8 column (5 µm, 9.4 mm × 250 mm) at 60 °C with a flow rate of 4 mL/min and an initial isocratic gradient of 1 %B for 5 min followed by a gradient of 0.5 %B/min over 60 min starting at 1 %B. Fractions identified with the correct $m/z$ and high purity by LCMS analysis (Figure S29) were combined and lyophilized to afford RS-pS$_{225}$ as a white amorphous solid (0.6 mg, ∼ 2 % yield based on resin loading, >95 % purity by UHPLC, Figure S30).

*f. RS-2×pS* The peptide corresponding to the RS domain of SRSF1 (residues 198-248) with terminal cysteines bearing phosphorylations at Ser223 and Ser225 was synthesized on NovaPEG Rink Amide resin (150 mg, 30 µmol) using the standard AFPS protocol. Fmoc-Ser(PO(OBzl)OH)-OH was incorporated at positions Ser223 and Ser225. Cleavage of the peptidyl-resin (32 mg, approx. 2.9 µmol) afforded the crude peptide as a colorless solid (7.5 mg, 24 % purity by LCMS, 24 % purity by UHPLC, Figures S31 and S32). The purification of the crude material (7.4 mg) was carried out by semi-preparative RP-HPLC using an Agilent Eclipse C18 column (5 µm, 9.4 mm × 250 mm) at 60 °C with a flow rate of 3.5 mL/min and an initial isocratic gradient of 1 %B for 5 min followed by a gradient of 1.40 %B/min over 43 min starting at 1 %B. Fractions identified with the correct $m/z$ and high purity by LCMS analysis (Figure S33) were combined and lyophilized to afford RS-2×pS as a white amorphous solid (∼ 0.6 mg, ∼ 3 % yield based on resin loading, >95 % purity by UHPLC, Figure S34).

*g. RS-4×pS* The peptide corresponding to the RS domain of SRSF1 (residues 198-248) with terminal cysteines bearing phosphorylations at Ser199, Ser207, Ser217 and Ser225 was synthesized on commercially available NovaPEG Rink Amide resin (165.8 mg, 33 µmol) using the standard AFPS protocol. Since crude purity of the 2xpS peptide was already reduced three-fold compared to unphosphorylated peptides, Fmoc-Ser(PO(OBzl)OH)-OH was incorporated at positions Ser199, Ser207, Ser217 and Ser225 with 10.4 mL of "coupling solution". Cleavage of the peptidyl-resin (228 mg, approx. 1.7 µmol) afforded the crude peptide as a colorless solid (57 mg, mass determined by LCMS, 28 % purity by UHPLC, Figures S35 and S36). The purification of the crude material (29 mg) was carried out by semi-preparative RP-HPLC using an Agilent Eclipse C18 column (5 µm, 9.4 mm × 250 mm) at 60 °C with a flow rate of 3.5 mL/min and an initial isocratic gradient of 1 %B for 5 min followed by a gradient of 0.60 %B/min over 42.5 min starting at 1 %B. Fractions identified with the correct $m/z$ and high purity by LCMS analysis (Figure S37) were combined and lyophilized to afford RS-4×pS as a white amorphous solid (1.005 mg, 1.8 % yield based on resin loading, >95 % purity by UHPLC, Figure S38).

### Appendix A: Peptide synthesis data

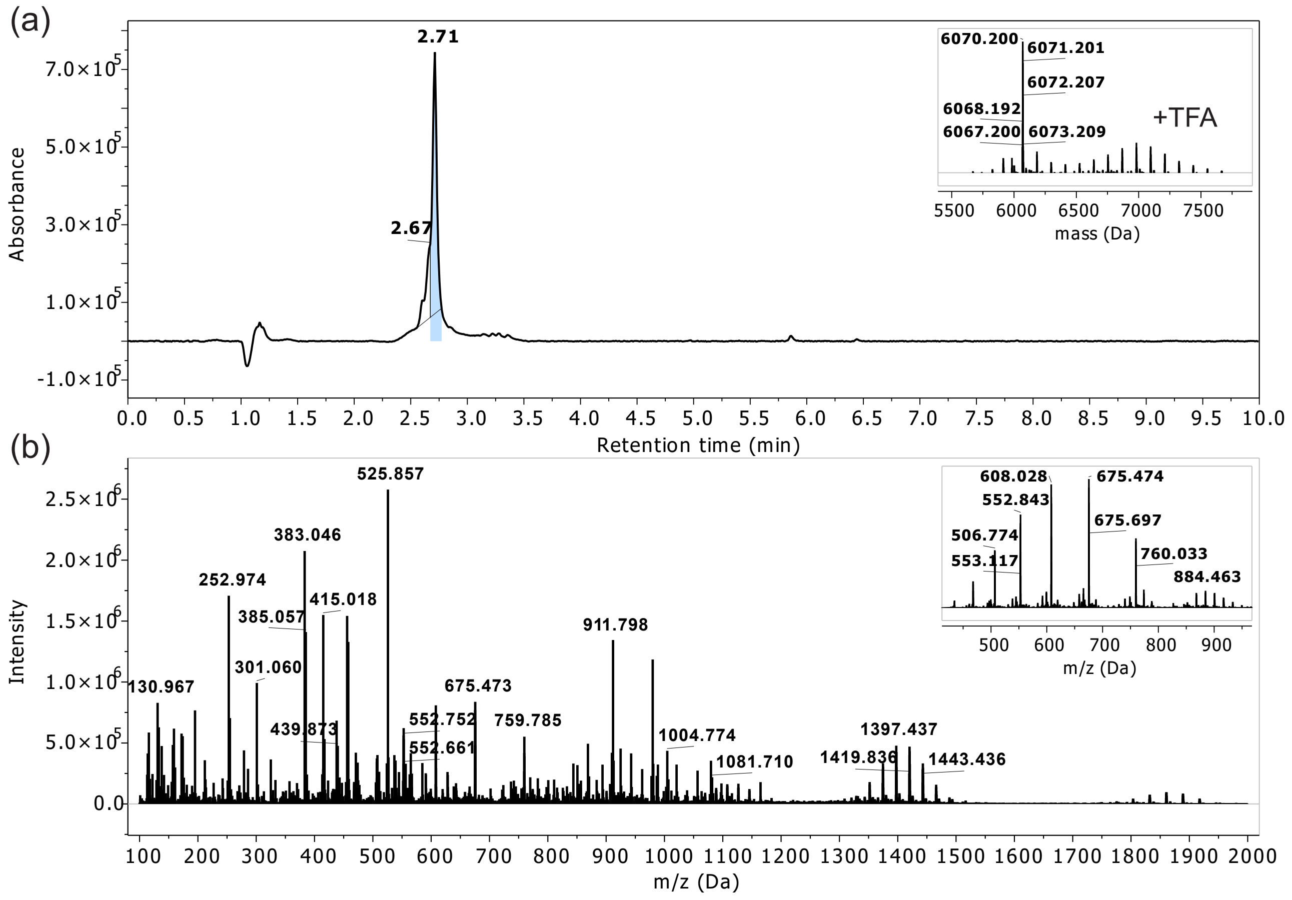


FIGURE S11. LCMS profile of crude RS-synthetic. (a) Absorbance chromatogram of RS at $\lambda = 214$ nm with a retention time (Rt) of 2.71 min (insert: deconvoluted masses). (b) ESI-TOF spectrum found within Rt 2 to 10 min (insert: convoluted spectra at Rt 2.71 min). Monoisotopic mass (ESI+) calculated for $C_{239}H_{412}N_{112}O_{76}$ is 6067.1817 Da; found 6067.1885 Da. LCMS Gradient A. Peptide ionizes with TFA, and mass corresponding to [Peptide + n*TFA] can be observed.

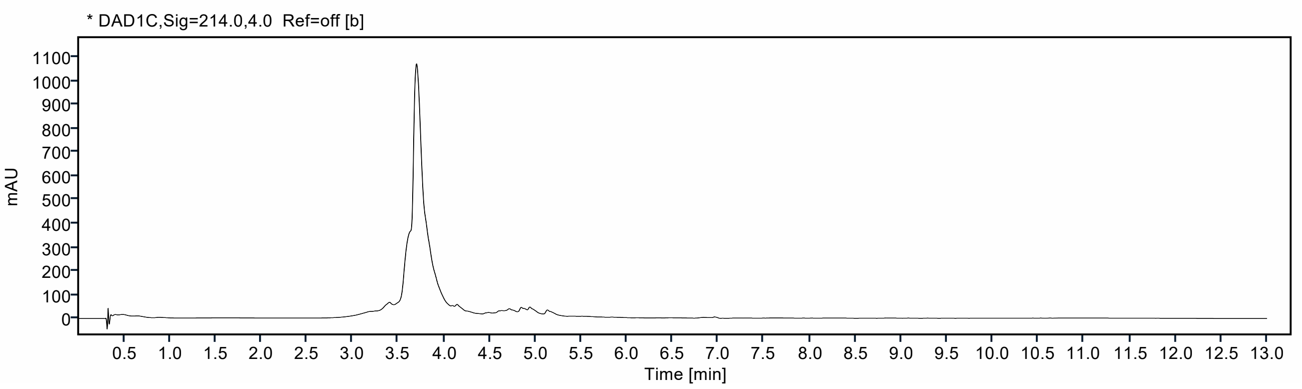


FIGURE S12. UHPLC profile of crude RS-synthetic using column A and a gradient of 5 to 95 % MeCN over 10 min at 40 °C. The peptide eluted at 3.70 min. Integration of the absorbance at $\lambda = 214$ nm between 1 to 10 min yields 72 % purity.

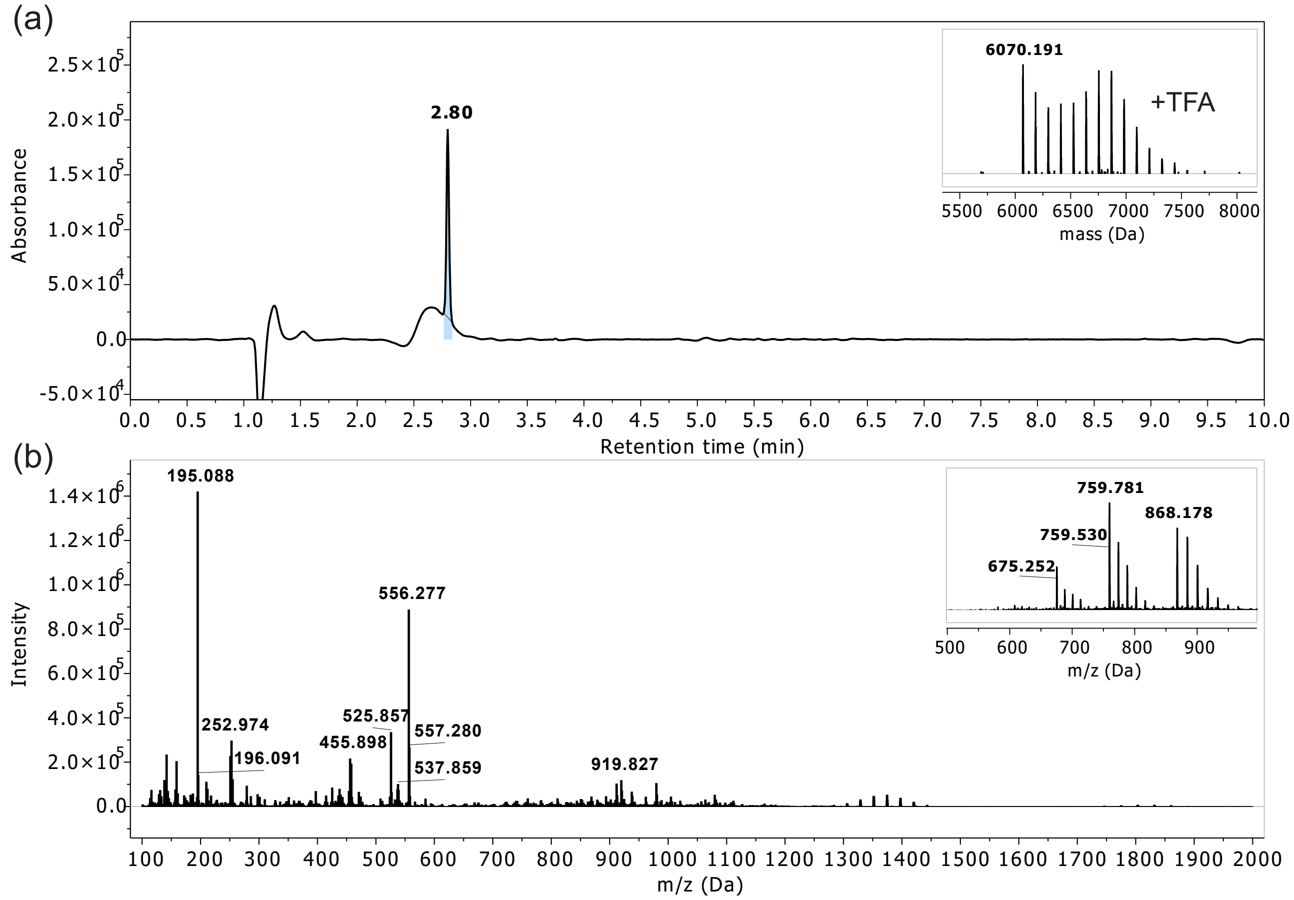


FIGURE S13. LCMS profile of pure RS-synthetic. (a) Absorbance chromatogram of pure RS at $\lambda = 214\,\mathrm{nm}$ with an Rt of 2.80 min (insert: deconvoluted masses). (b) ESI-TOF spectrum found within Rt 2 to 10 min (insert: convoluted spectra of RT 2.80 min). Monoisotopic mass (ESI+) calculated for $C_{239}H_{412}N_{112}O_{76}$ 6067.1817 Da, found 6067.1768 Da. LCMS Gradient A. Peptide ionizes with TFA, and mass corresponding to [Peptide + n*TFA] can be observed.

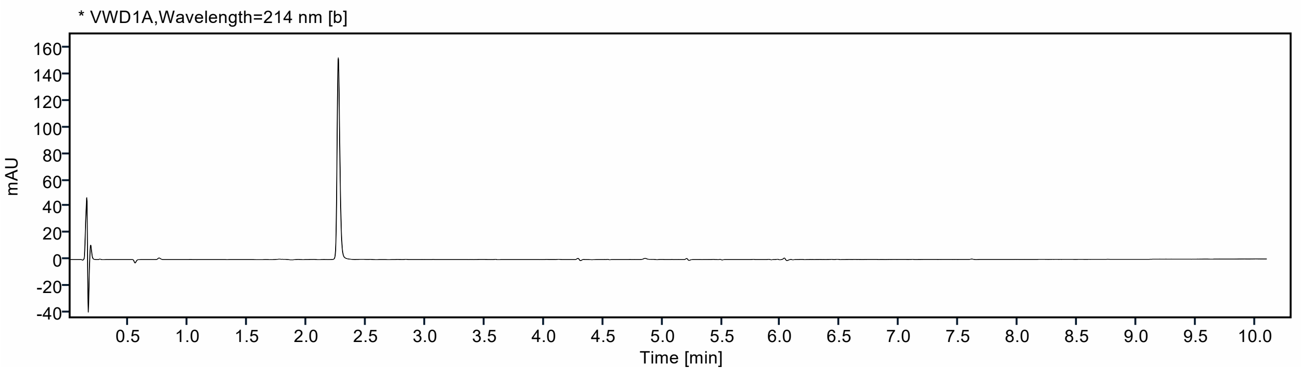


FIGURE S14. UHPLC profile of pure RS suing column B and a gradient of 0 to 100 % MeCN over 10 min at 40 °C. The peptide eluted at 2.36 min. Integration of the absorbance at $\lambda = 214\,\mathrm{nm}$ between 1 to 10 min yields 99 % purity.

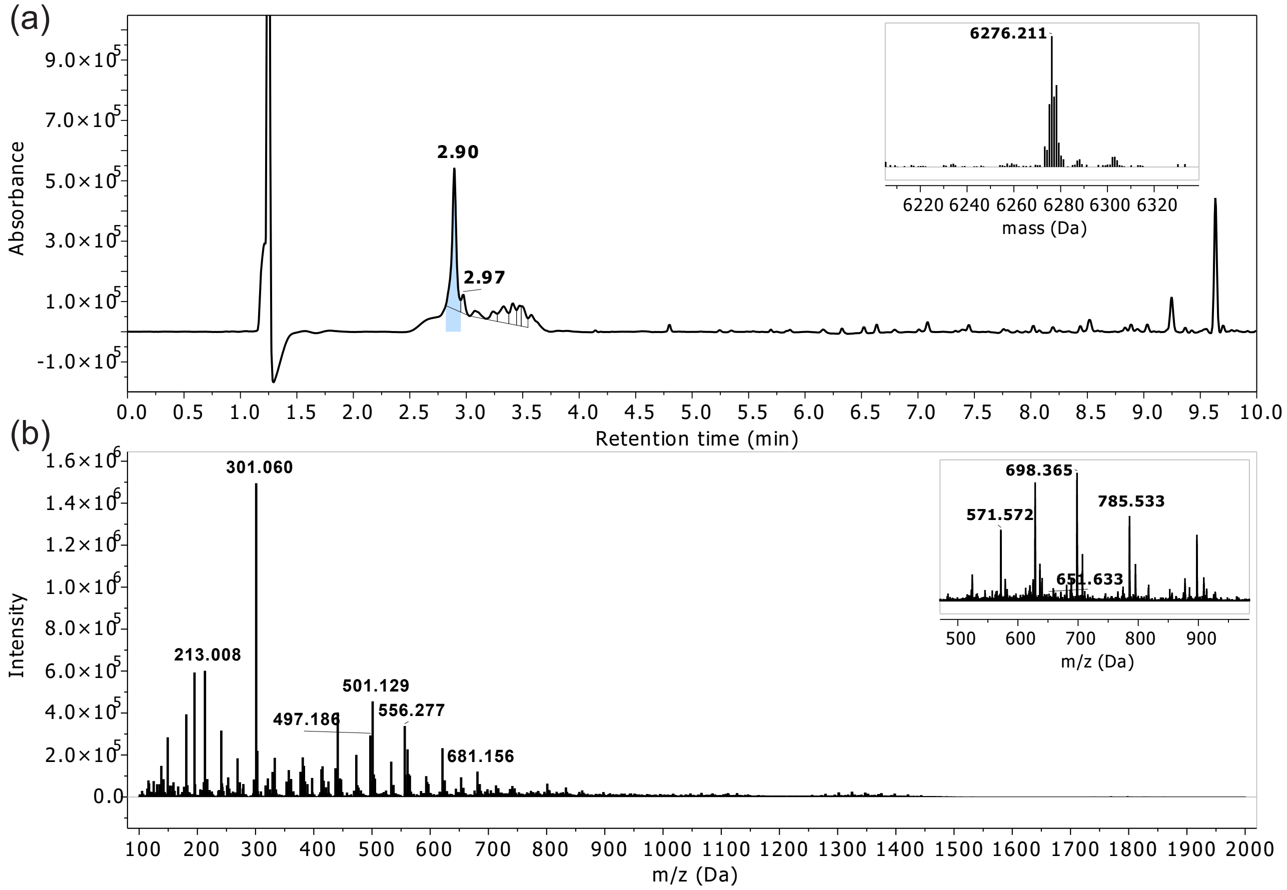


FIGURE S15. LCMS profile of crude cysRScys. (a) Absorbance chromatogram of cysRScys at $\lambda = 214\,\mathrm{nm}$ with an Rt of 2.90 min (insert: deconvoluted masses). (b) ESI-TOF spectrum found within Rt 2 to 10 min (insert: convoluted spectra at Rt 2.90 min). Monoisotopic mass (ESI+) calculated for $C_{245}H_{422}N_{114}O_{78}S_2$ is 6273.2001 Da; found 6273.1780 Da. LCMS Gradient B.

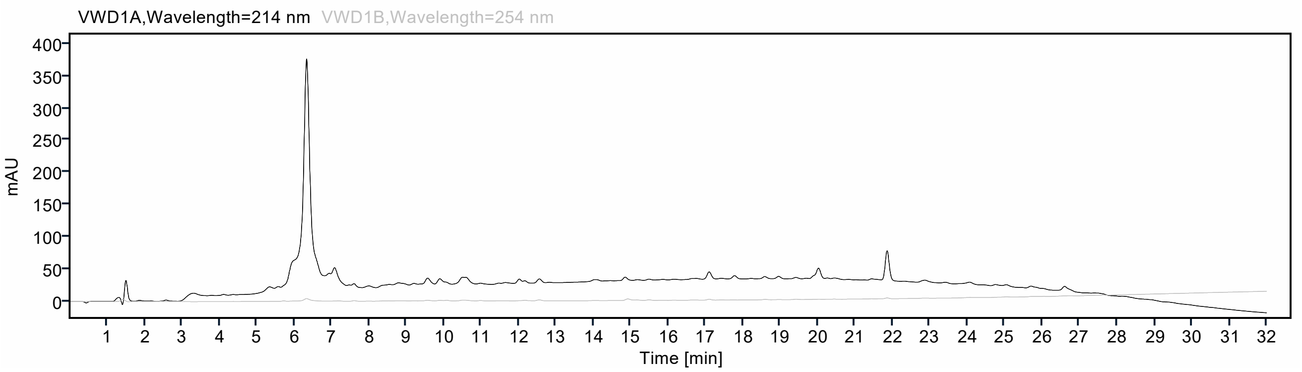


FIGURE S16. UHPLC profile of crude cysRScys using column C and a gradient of 0 to 100 % MeCN over 30 min at 40 °C. The peptide eluted at 6.34 min. Integration of the absorbance at $\lambda = 214\,\mathrm{nm}$ between 2 to 30 min yields 71 % purity.

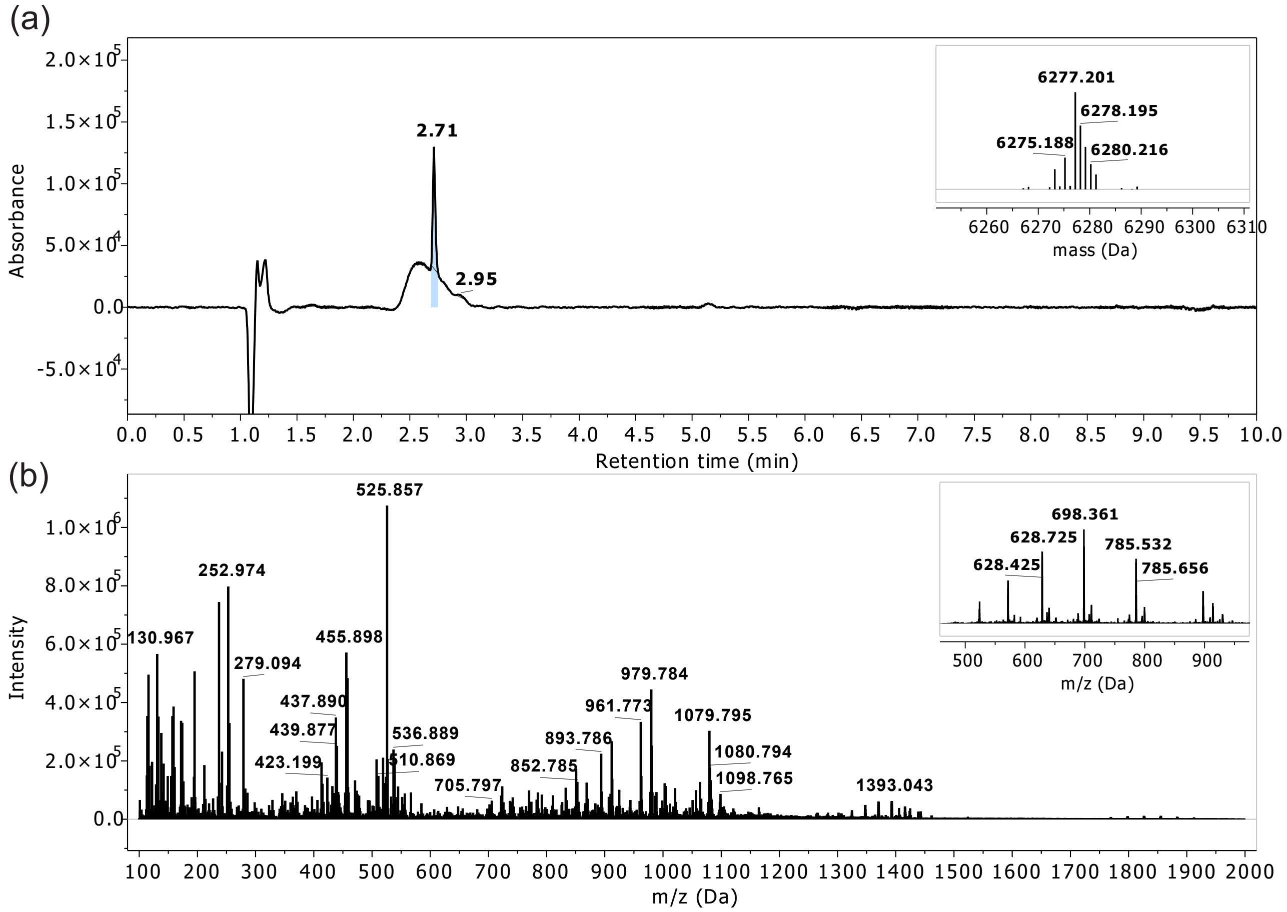


FIGURE S17. LCMS profile of pure cysRScys. (a) Absorbance chromatogram of cysRScys at $\lambda = 214$ nm with an Rt of 2.71 min (insert: deconvoluted masses). (b) ESI-TOF spectrum found within Rt 2 to 10 min (insert: convoluted spectra at Rt 2.71 min). Monoisotopic mass (ESI+) calculated for $C_{245}H_{422}N_{114}O_{78}S_2$ is 6273.2001 Da; found 6273.1808 Da. LCMS Gradient B. Peptide ionizes with TFA, and mass corresponding to [Peptide + n*TFA] can be observed.

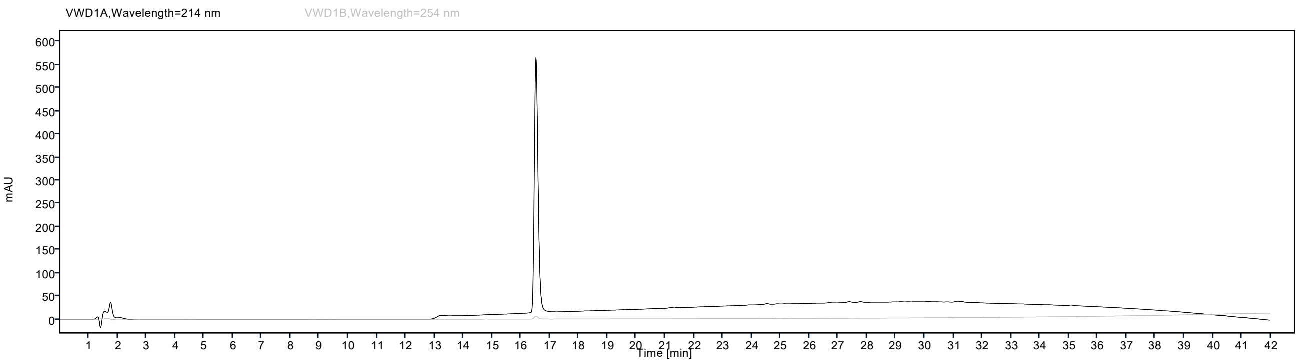


FIGURE S18. UHPLC profile of pure cysRScys using column C and a gradient of 0 to 100 % MeCN over 40 min at 40 °C. The peptide eluted at 16.52 min. Integration of the absorbance at $\lambda = 214$ nm between 3 to 40 min yields 99 % purity.

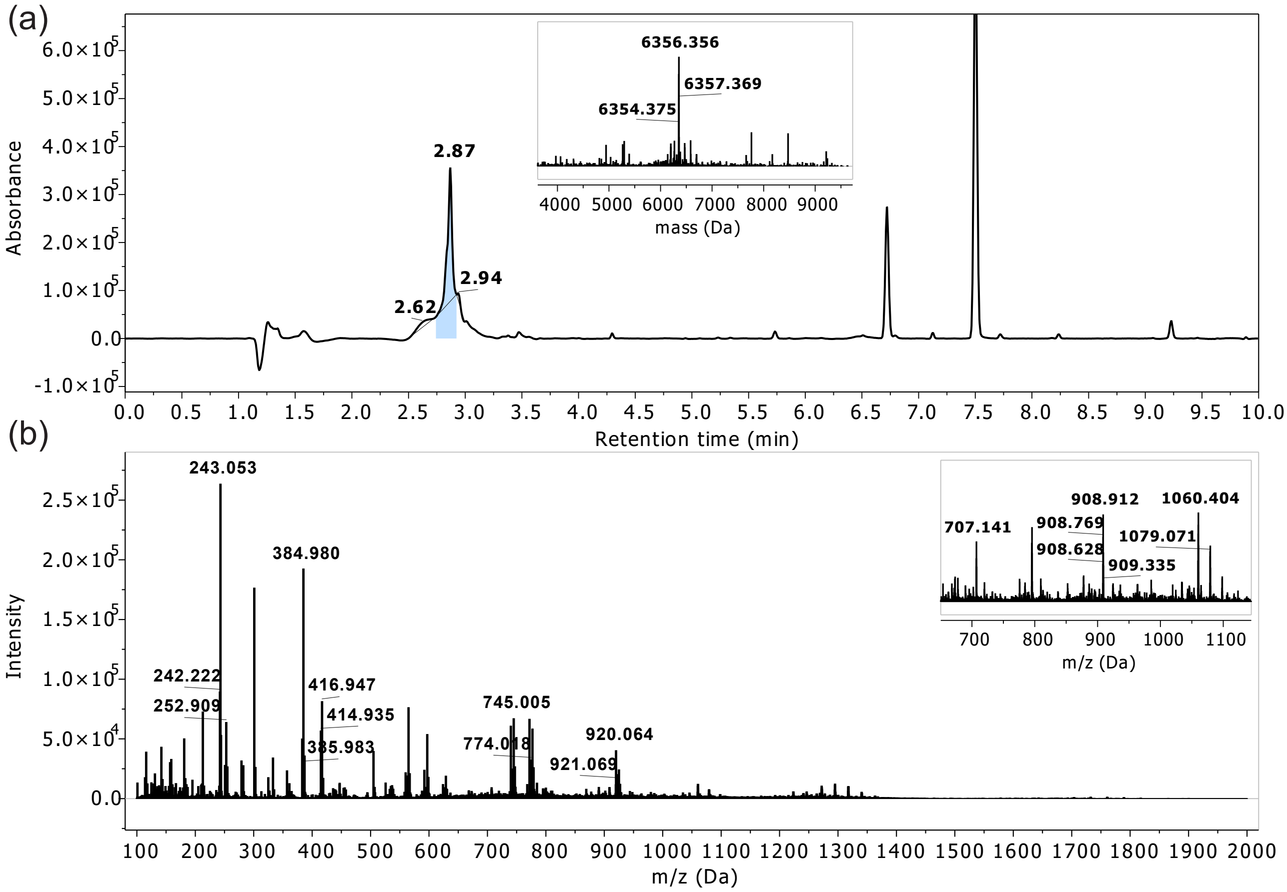


FIGURE S19. LCMS profile of crude RS-pS199. (a) Absorbance chromatogram of RS-pS199 at $\lambda$ = 214 nm with an Rt of 2.87 min (insert: deconvoluted masses). (b) ESI-TOF spectrum found within Rt 2 to 10 min (insert: convoluted spectra at Rt 2.87 min). Monoisotopic mass (ESI+) calculated for $C_{245}H_{423}N_{114}O_{81}PS_2$ is 6353.1701 Da; found 6353.0729 Da. LCMS Gradient A.

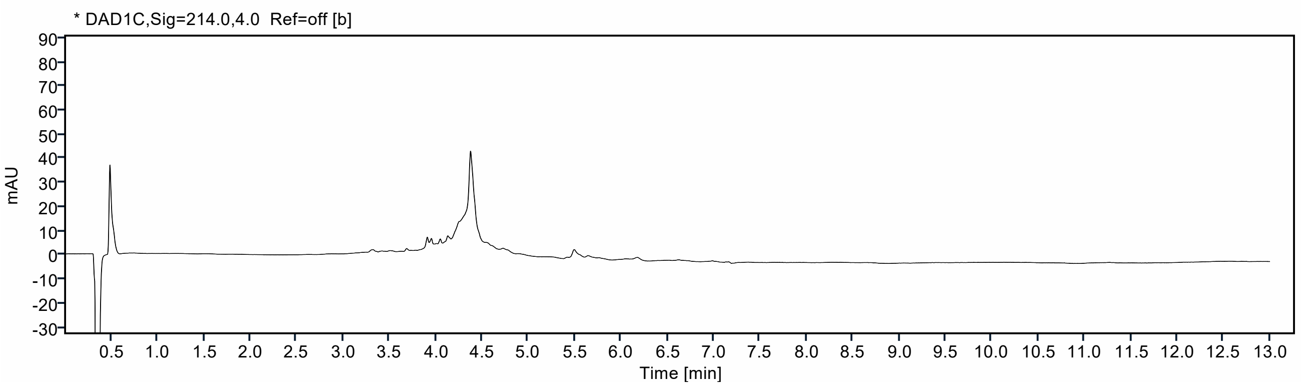


FIGURE S20. UHPLC profile of crude RS-pS199 using an column A and a gradient of 5 to 95 % MeCN over 10 min at 40 °C. The peptide eluted at 4.38 min. Integration of the absorbance at $\lambda$ = 214 nm between 1 to 10 min yields 49 % purity.

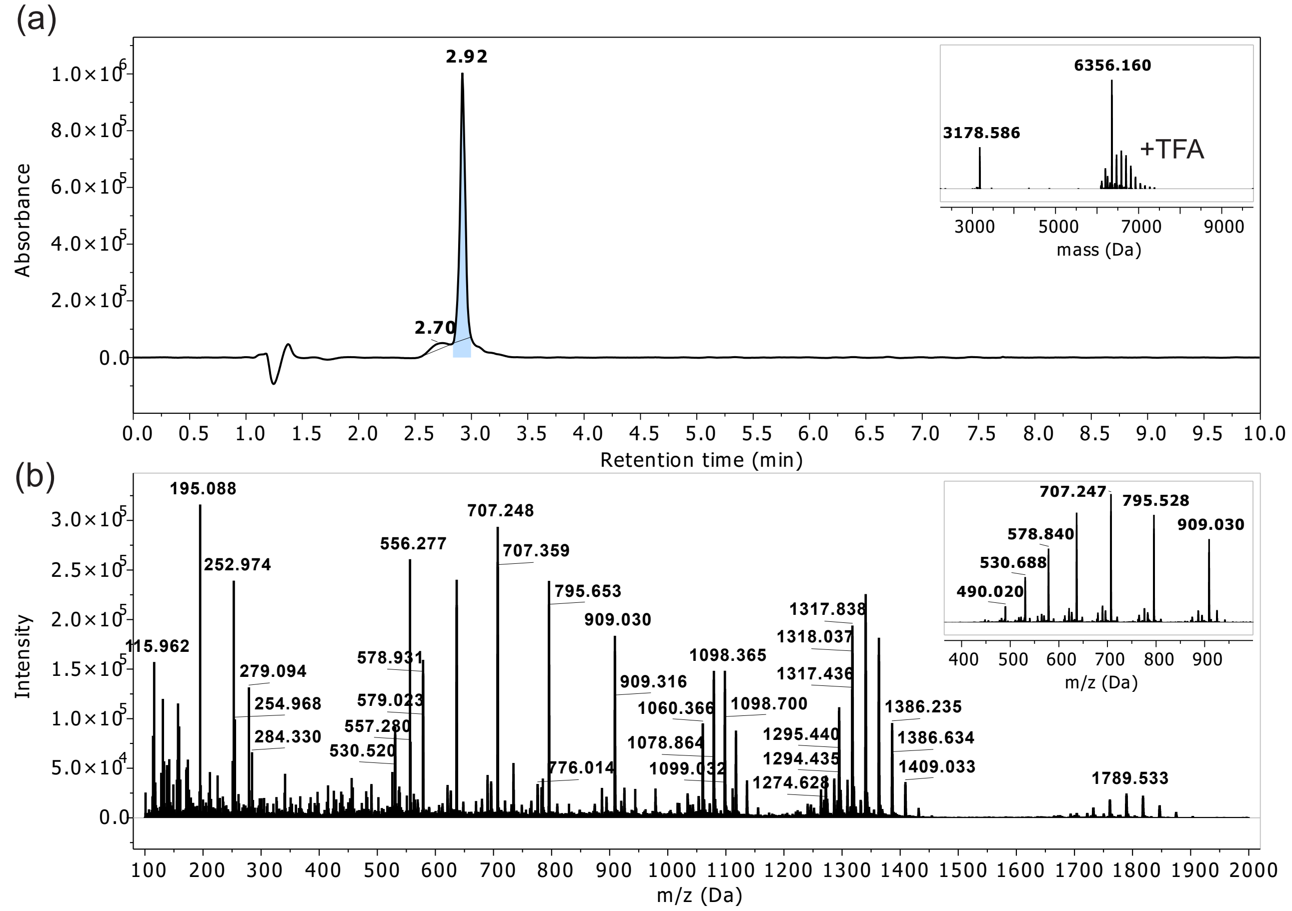


FIGURE S21. LCMS profile of pure RS-pS199. (a) Absorbance chromatogram of pure RS-pS199 at $\lambda = 214$ nm with an Rt of 2.92 min (insert: deconvoluted masses). (b) ESI-TOF spectrum found within Rt 2 to 10 min (insert: convoluted spectra at Rt 2.92 min). Monoisotopic mass (ESI+) calculated for $C_{245}H_{423}N_{114}O_{81}PS_2$ is 6353.1701 Da; found 6354.1582 Da. LCMS Gradient A. Peptide ionizes with TFA, and mass corresponding to [Peptide + n*TFA] can be observed.

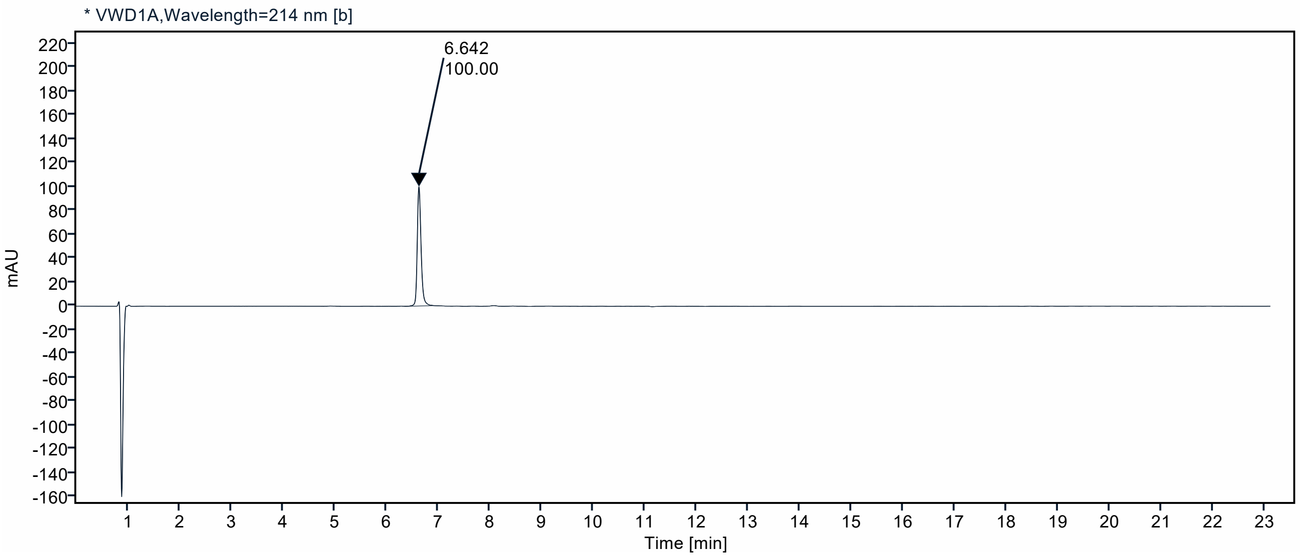


FIGURE S22. UHPLC profile of pure RS-pS199 using column C and a gradient of 0 to 100 % MeCN over 20 min at 40 °C. The peptide eluted at 6.642 min. Integration of the absorbance at $\lambda = 214$ nm between 2 to 20 min yields 100 % purity.

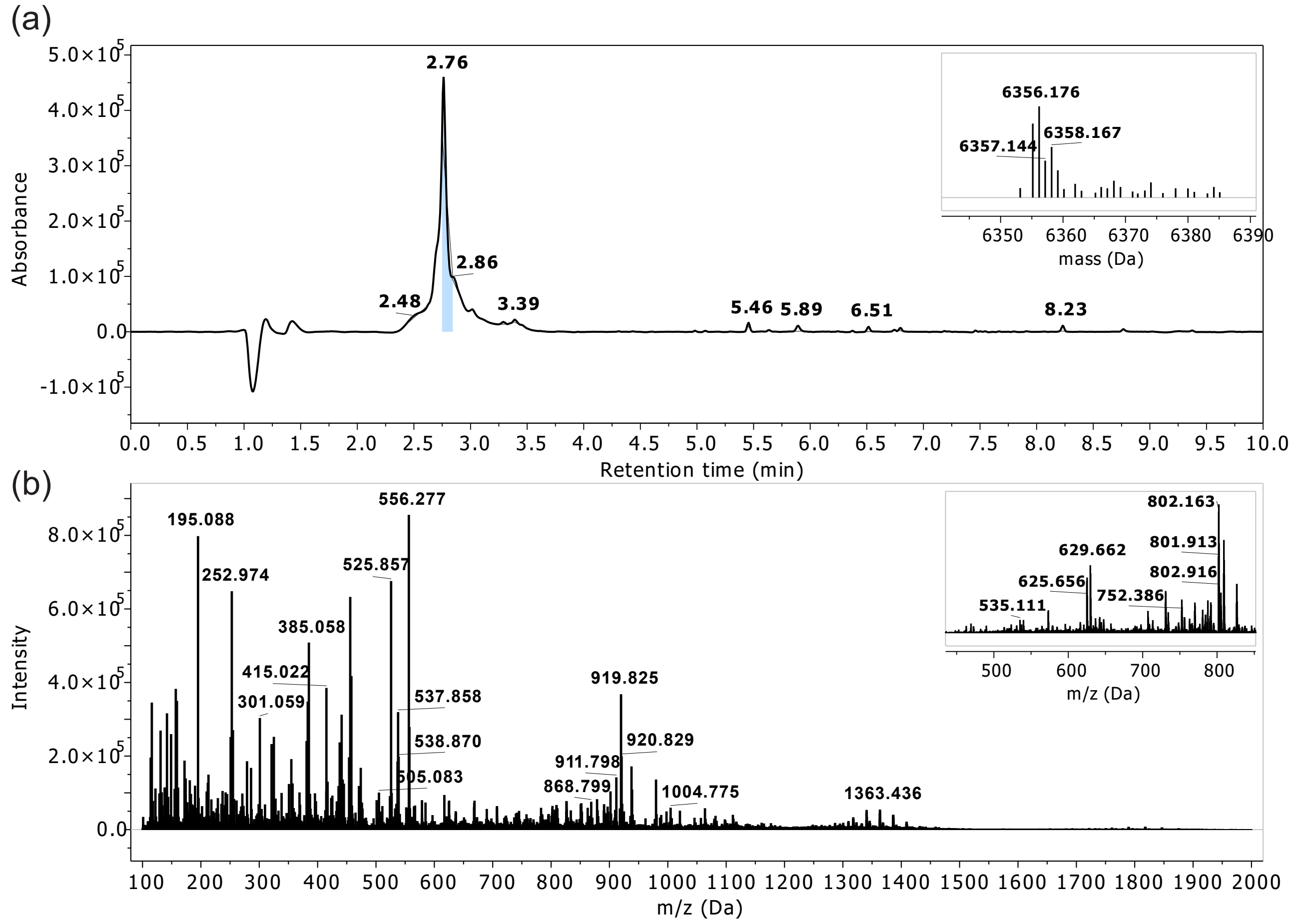


FIGURE S23. LCMS profile of crude RS-pS223. (a) Absorbance chromatogram of RS-pS223 at $\lambda$ = 214 nm with an Rt of 2.76 min (insert: deconvoluted masses). (b) ESI-TOF spectrum found within Rt 2 to 10 min (insert: convoluted spectra at Rt 2.76 min). Monoisotopic mass (ESI+) calculated for $C_{245}H_{423}N_{114}O_{81}PS_2$ is 6353.1701 Da; found 6353.1527 Da. LCMS Gradient B.

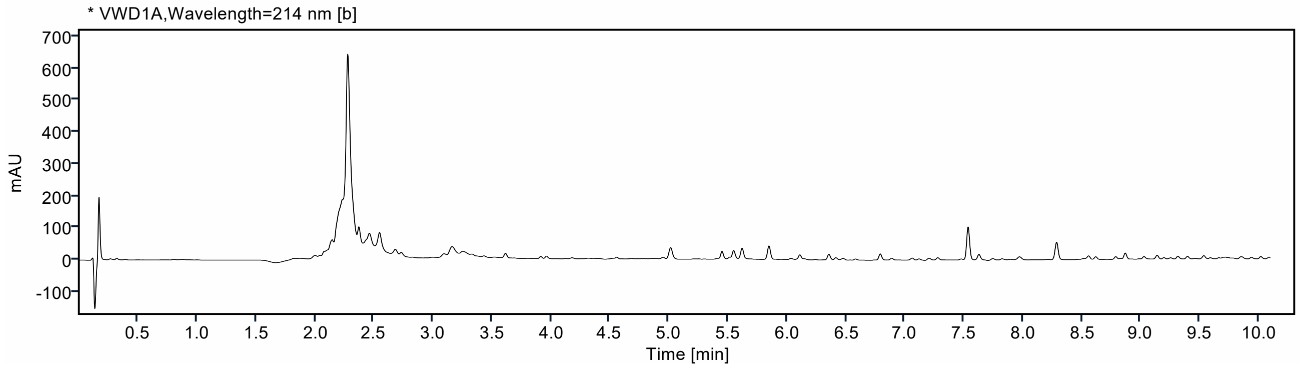


FIGURE S24. UHPLC profile of crude RS-pS223 using an column B and a gradient of 0 to 100 % MeCN over 10 min at 40 °C. The peptide eluted at 2.27 min. Integration of the absorbance at $\lambda$ = 214 nm between 1 to 10 min yields 51 % purity.

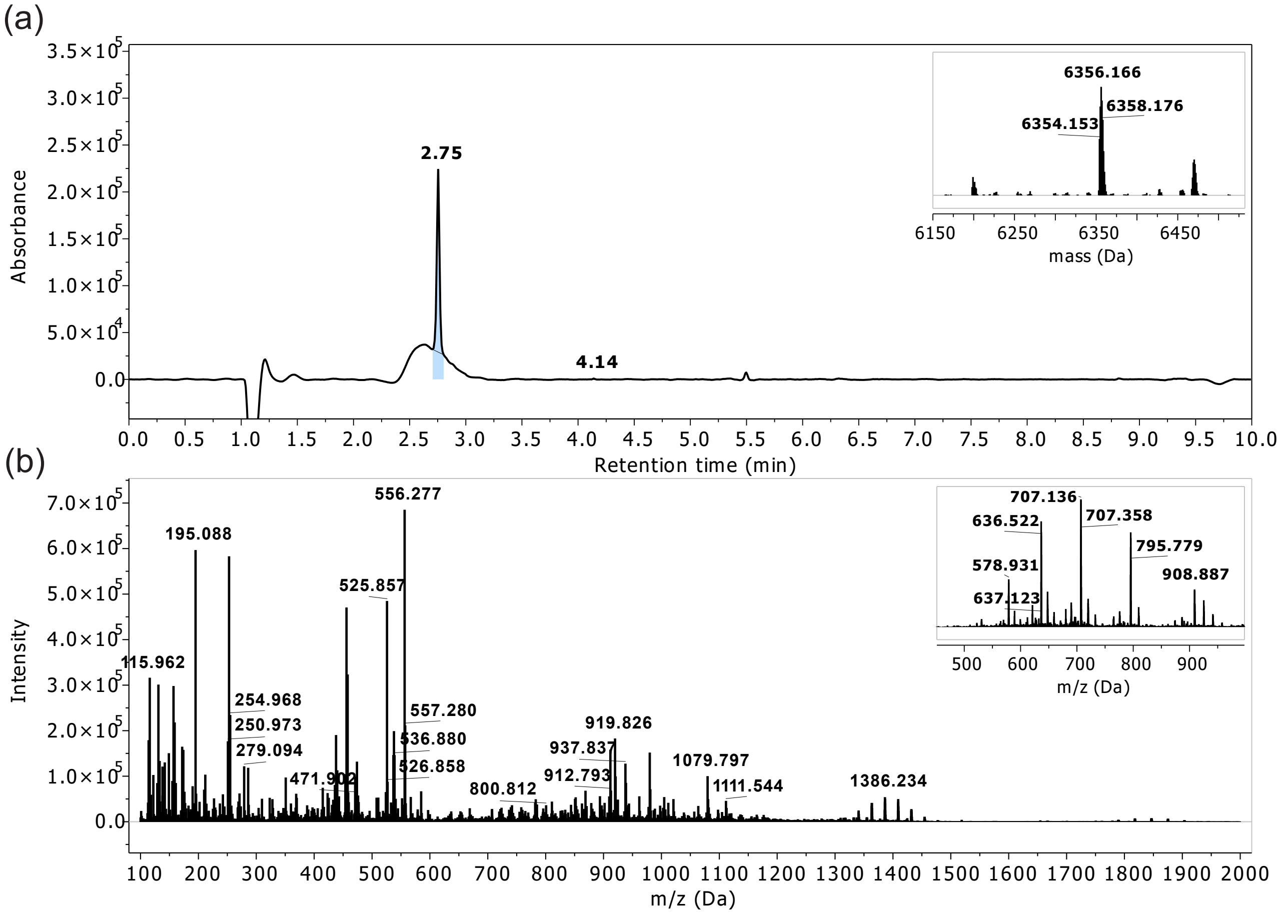


FIGURE S25. LCMS profile of pure RS-pS223. (a) Absorbance chromatogram of pure RS-pS223 at $\lambda = 214$ nm with an Rt of 2.75 min (insert: deconvoluted masses). (b) ESI-TOF spectrum found within Rt 2 to 10 min (insert: convoluted spectra at Rt 2.75 min). Monoisotopic mass (ESI+) calculated for $C_{245}H_{423}N_{114}O_{81}PS_2$ is 6353.1701 Da; found 6353.1682 Da. LCMS Gradient B.

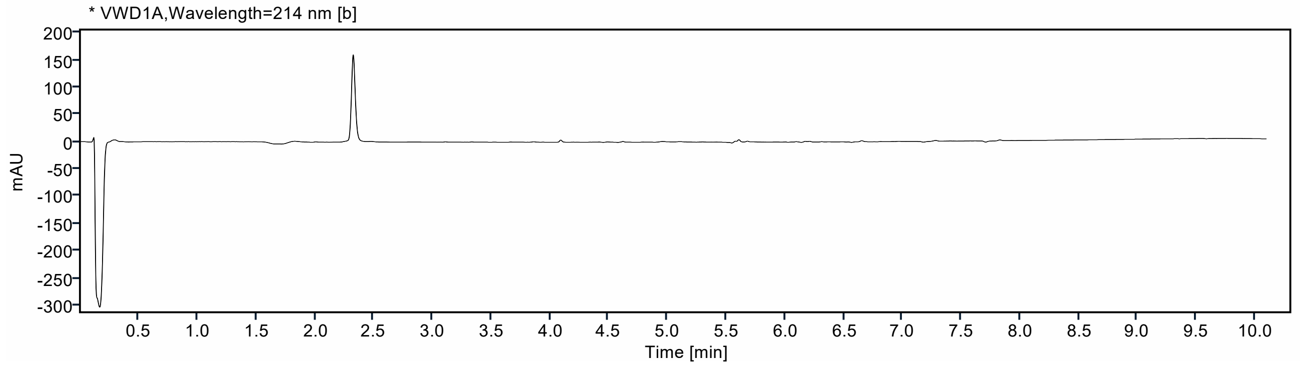


FIGURE S26. UHPLC profile of pure RS-pS223 using column B and a gradient of 0 to 100 % MeCN over 10 min at 40 °C. The peptide eluted at 2.33 min. Integration of the absorbance at $\lambda = 214$ nm between 1 to 10 min yields 95 % purity.

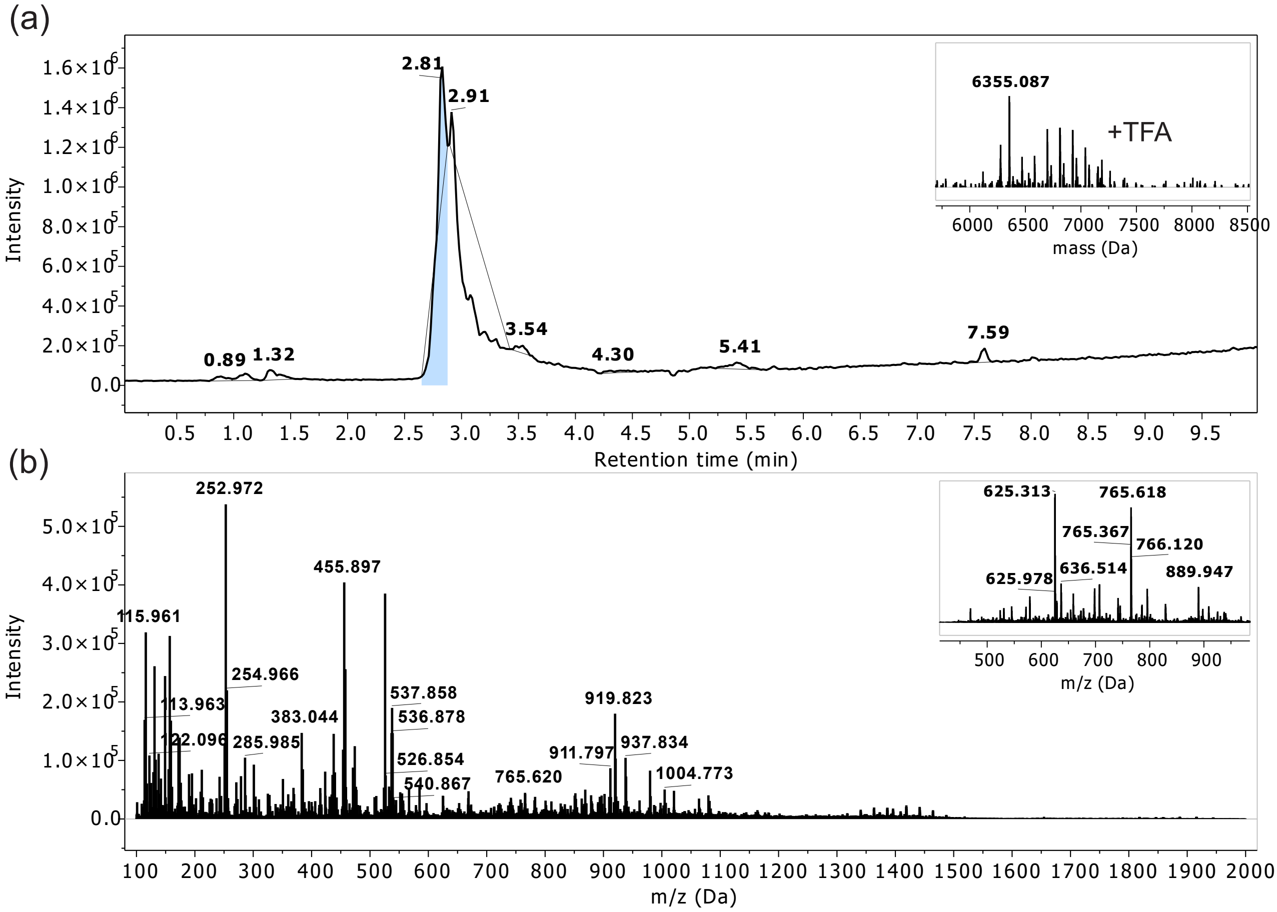


FIGURE S27. LCMS profile of crude RS-pS225. (a) Total Ion Chromatogram (TIC) of RS-pS225 at $\lambda = 214$ nm with an Rt of 2.81 min (insert: deconvoluted masses). (b) ESI-TOF spectrum found within Rt 2 to 10 min (insert: convoluted spectra at Rt 2.81 min). Monoisotopic mass (ESI+) calculated for $C_{245}H_{423}N_{114}O_{81}PS_2$ is 6353.1701 Da; found 6353.077 Da. LCMS Gradient A. Peptide ionizes with TFA, and mass corresponding to [Peptide + n*TFA] can be observed.

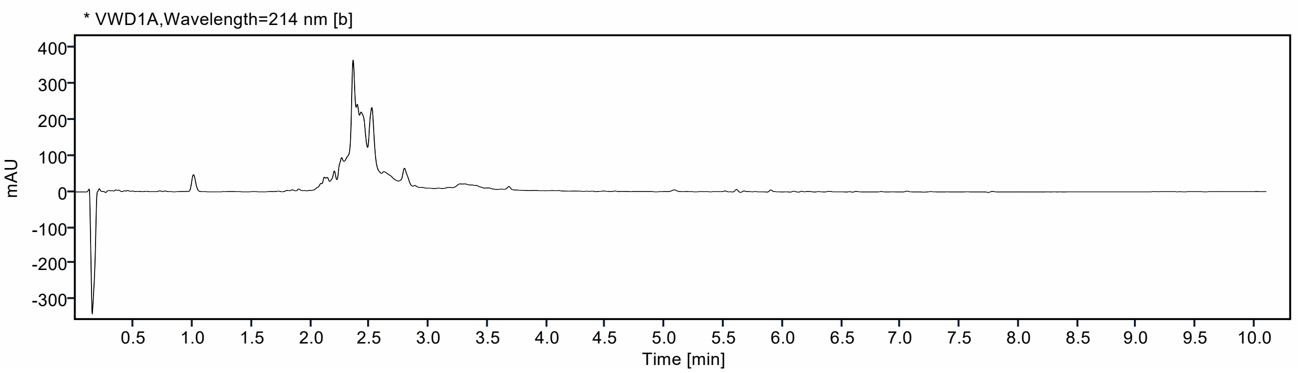


FIGURE S28. UHPLC profile of crude RS-pS225 using column B and a gradient of 0 to 100 % MeCN over 10 min at 40 °C. The peptide eluted at 2.36 min. Integration of the absorbance at $\lambda = 214$ nm between 0.5 to 10 min yields 32 % purity.

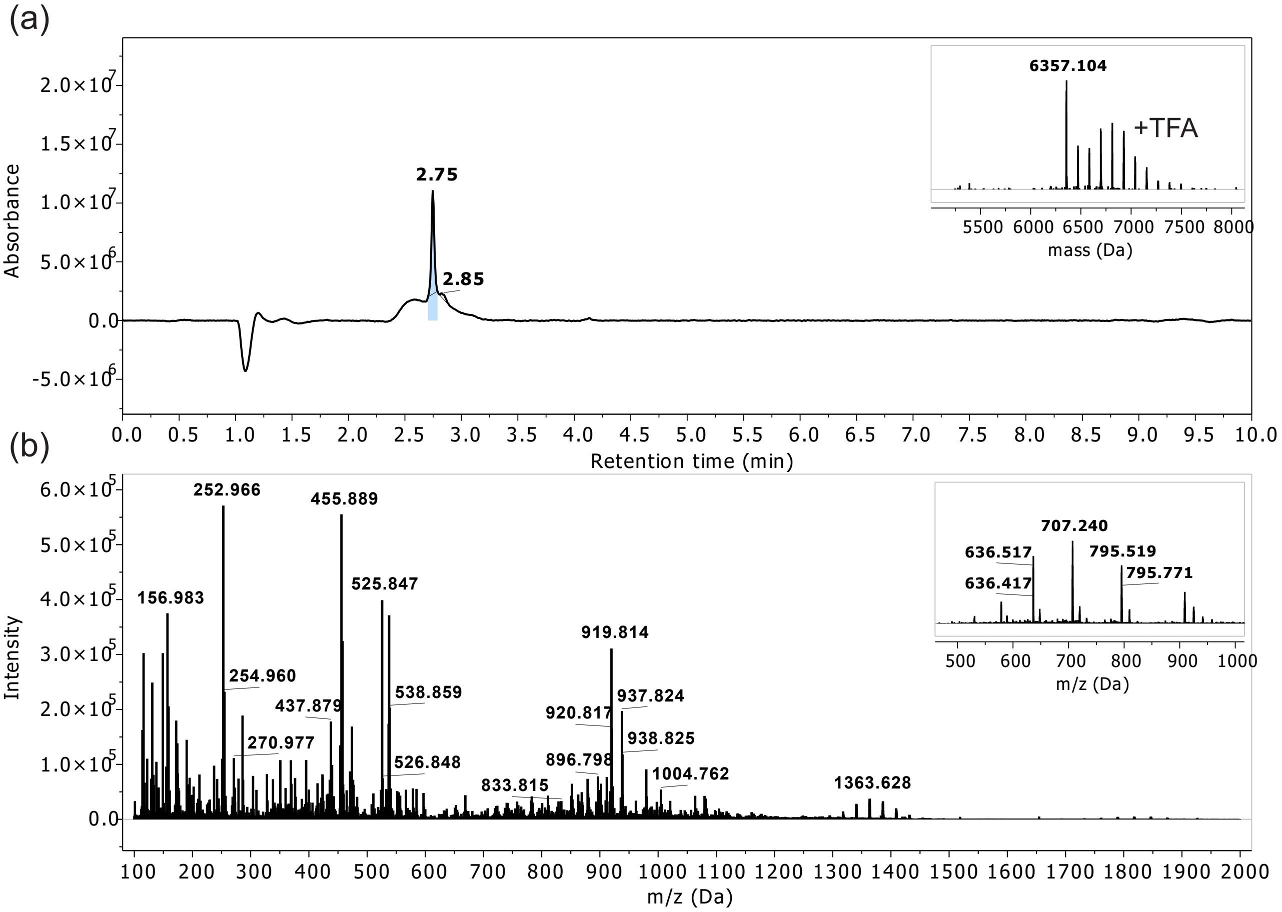


FIGURE S29. LCMS profile of pure RS-pS225. (a) Absorbance chromatogram of pure RS-pS225 at $\lambda = 214$ nm with an Rt of 2.75 min (insert: deconvoluted masses). (b) ESI-TOF spectrum found within Rt 2 to 10 min (insert: convoluted spectra at Rt 2.75 min). Monoisotopic mass (ESI+) calculated for $C_{245}H_{423}N_{114}O_{81}PS_2$ is 6353.1701 Da; found 6353.0933 Da. LCMS Gradient B. Peptide ionizes with TFA, and mass corresponding to [Peptide + n*TFA] can be observed.

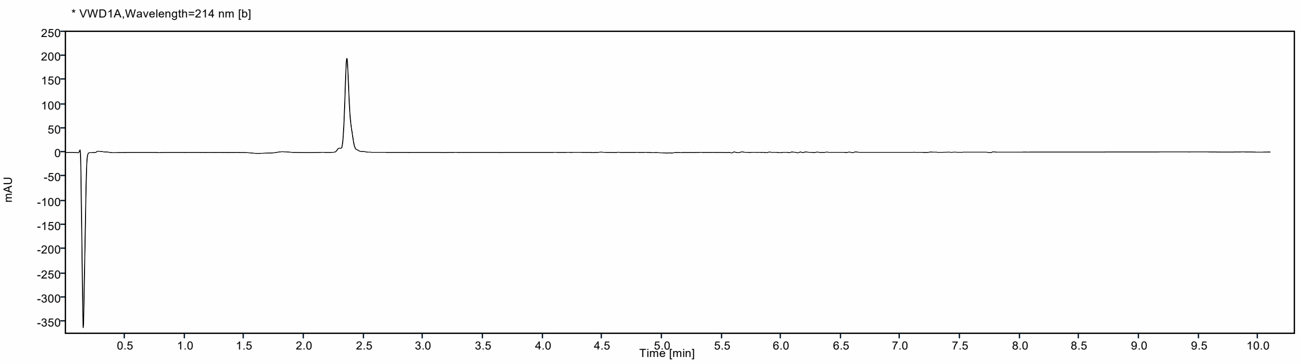


FIGURE S30. UHPLC profile of pure RS-pS225 using column B and a gradient of 0 to 100 % MeCN over 10 min at 40 °C. The peptide eluted at 2.36 min. Integration of the absorbance at $\lambda = 214$ nm between 1 to 10 min yields 96 % purity.

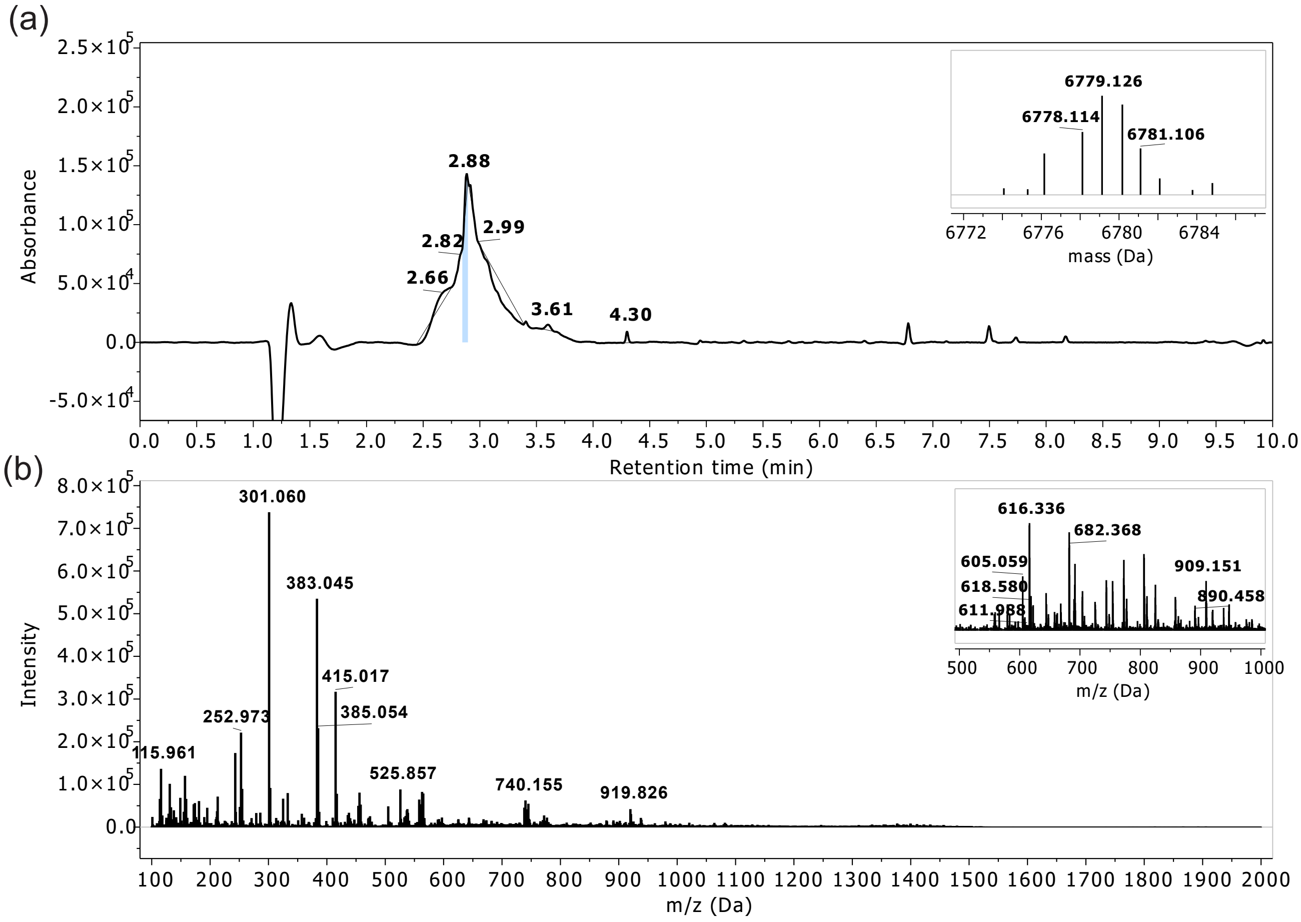


FIGURE S31. LCMS profile of crude RS-2×pS. (a) Absorbance chromatogram of RS-2×pS at $\lambda = 214$ nm with an Rt of 2.88 min (insert: deconvoluted masses). (b) ESI-TOF spectrum found within Rt 2 to 10 min (insert: convoluted spectra at Rt 2.88 min). Monoisotopic mass (ESI+) calculated for $[C_{246}H_{424}N_{114}O_{85}P_2S_2 + 2\text{*TFA}]$ is 6775.0401 Da; found 6775.324 Da. LCMS Gradient A.

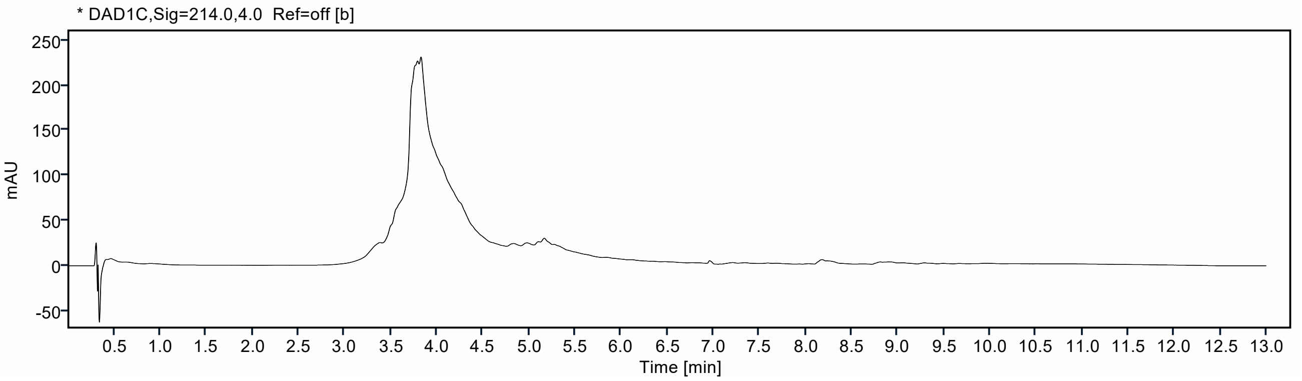


FIGURE S32. UHPLC profile of crude RS-2×pS using column A and a gradient of 5 to 95 % MeCN over 10 min at 40 °C. The peptide eluted at 3.79 min. Integration of the absorbance at $\lambda = 214$ nm between 1 to 10 min yields 24 % purity.

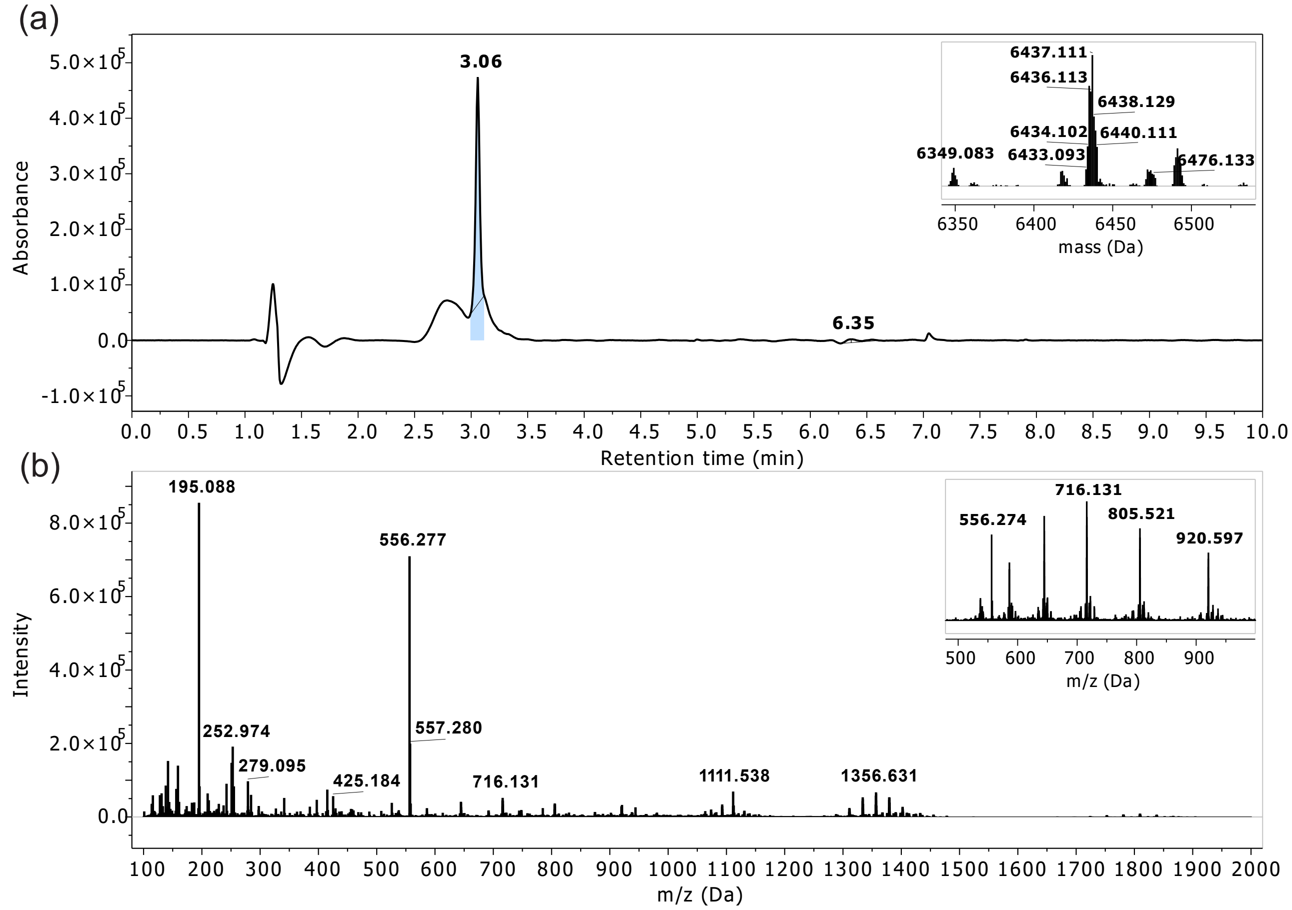


FIGURE S33. LCMS profile of pure RS-2×pS. (a) Absorbance chromatogram of pure RS-2×pS at $\lambda = 214\,\text{nm}$ with an Rt of 3.06 min (insert: deconvoluted masses). (b) ESI-TOF spectrum found within Rt 2 to 10 min (insert: convoluted spectra at Rt 3.06 min). Monoisotopic mass (ESI+) calculated for $C_{245}H_{424}N_{114}O_{84}P_2S_2$ is 6433.1327 Da; found 6433.1041 Da. LCMS Gradient A.

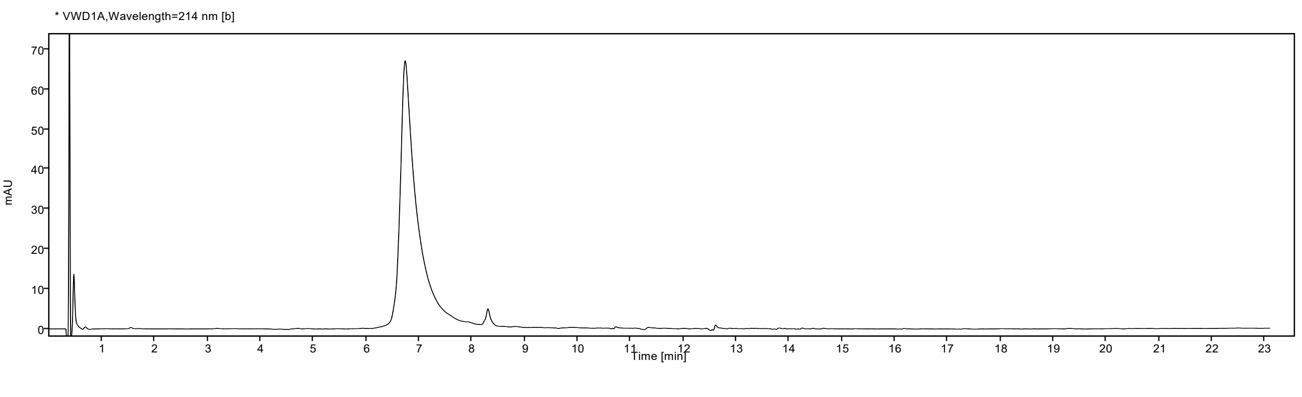


FIGURE S34. UHPLC profile of pure RS-2×pS using column B and a gradient of 0 to 100 % MeCN over 20 min at 40 °C. The peptide eluted at 6.74 min. Integration of the absorbance at $\lambda = 214\,\text{nm}$ between 1 to 20 min yields 98 % purity.

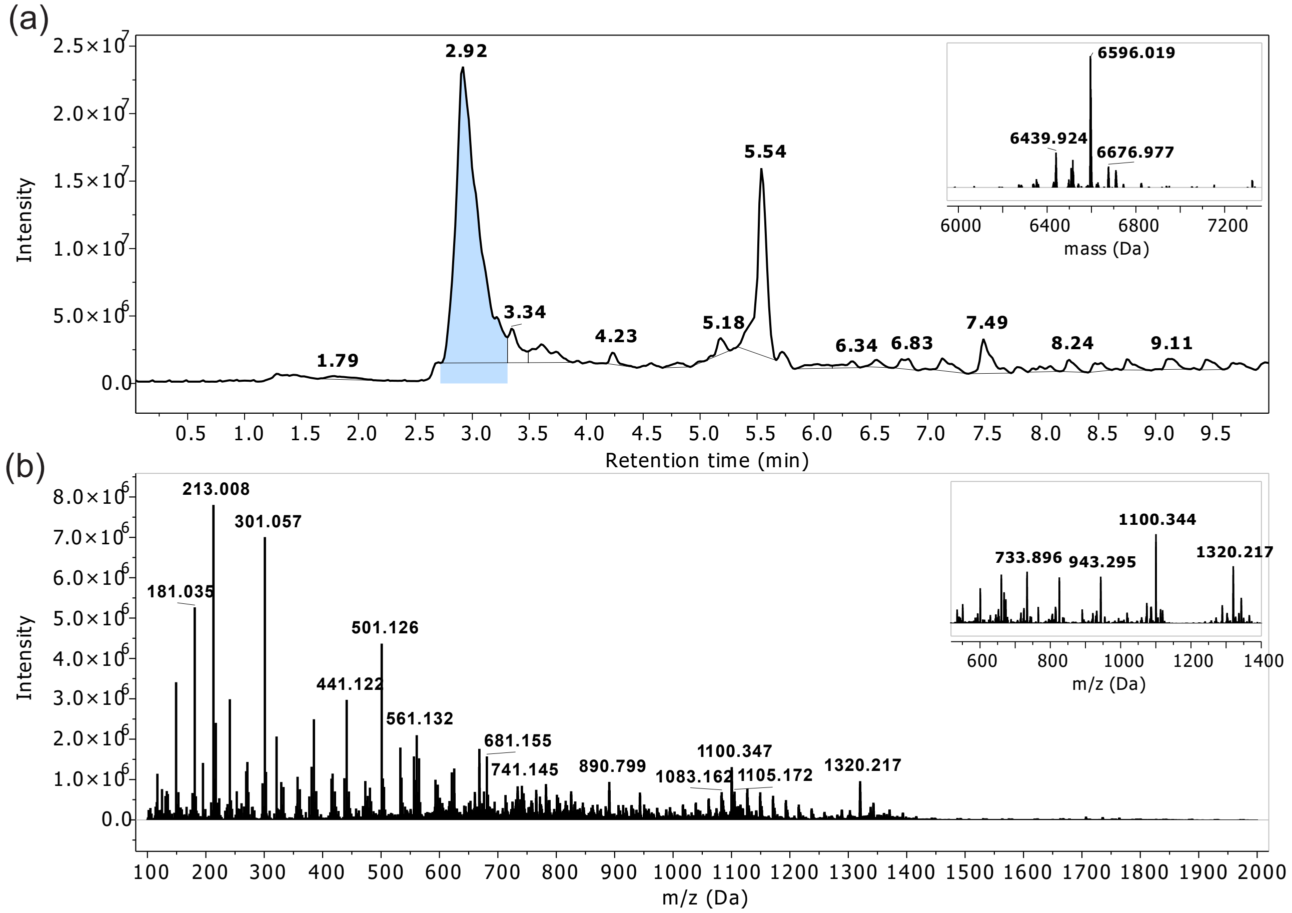


FIGURE S35. LCMS profile of crude RS-4×pS. (a) TI) of RS-4×pS at $\lambda = 214$ nm with an Rt of 2.92 min (insert: deconvoluted masses). (b) ESI-TOF spectrum found within Rt 2 to 10 min (insert: convoluted spectra at Rt 2.92 min). Monoisotopic mass (ESI+) calculated for $C_{245}H_{426}N_{114}O_{90}P_4S_2$ is 6593.0654 Da; found 6593.0037 Da. LCMS Gradient A.

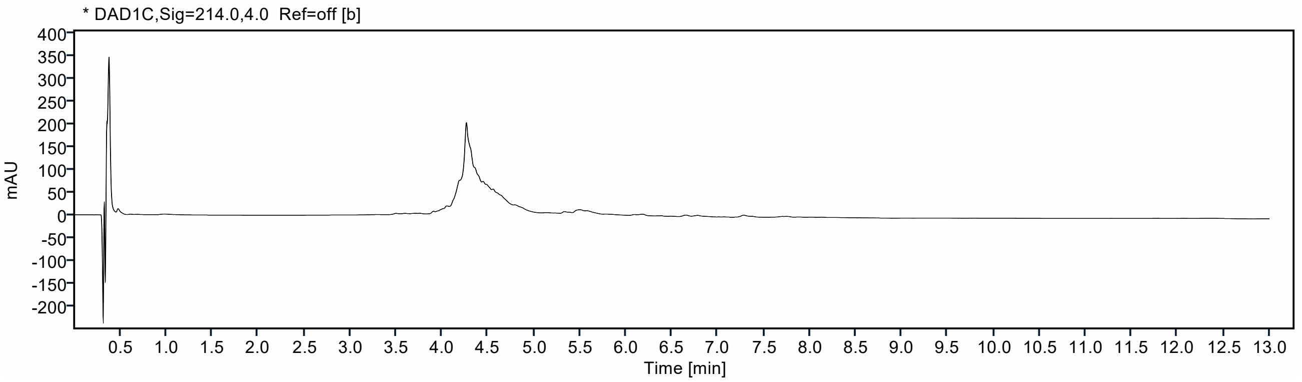


FIGURE S36. UHPLC profile of crude RS-4×pS using column A and a gradient of 5 to 95 % MeCN over 10 min at 40 °C. The peptide eluted at 4.27 min. Integration of the absorbance at $\lambda = 214$ nm between 1 to 10 min yields 28 % purity.

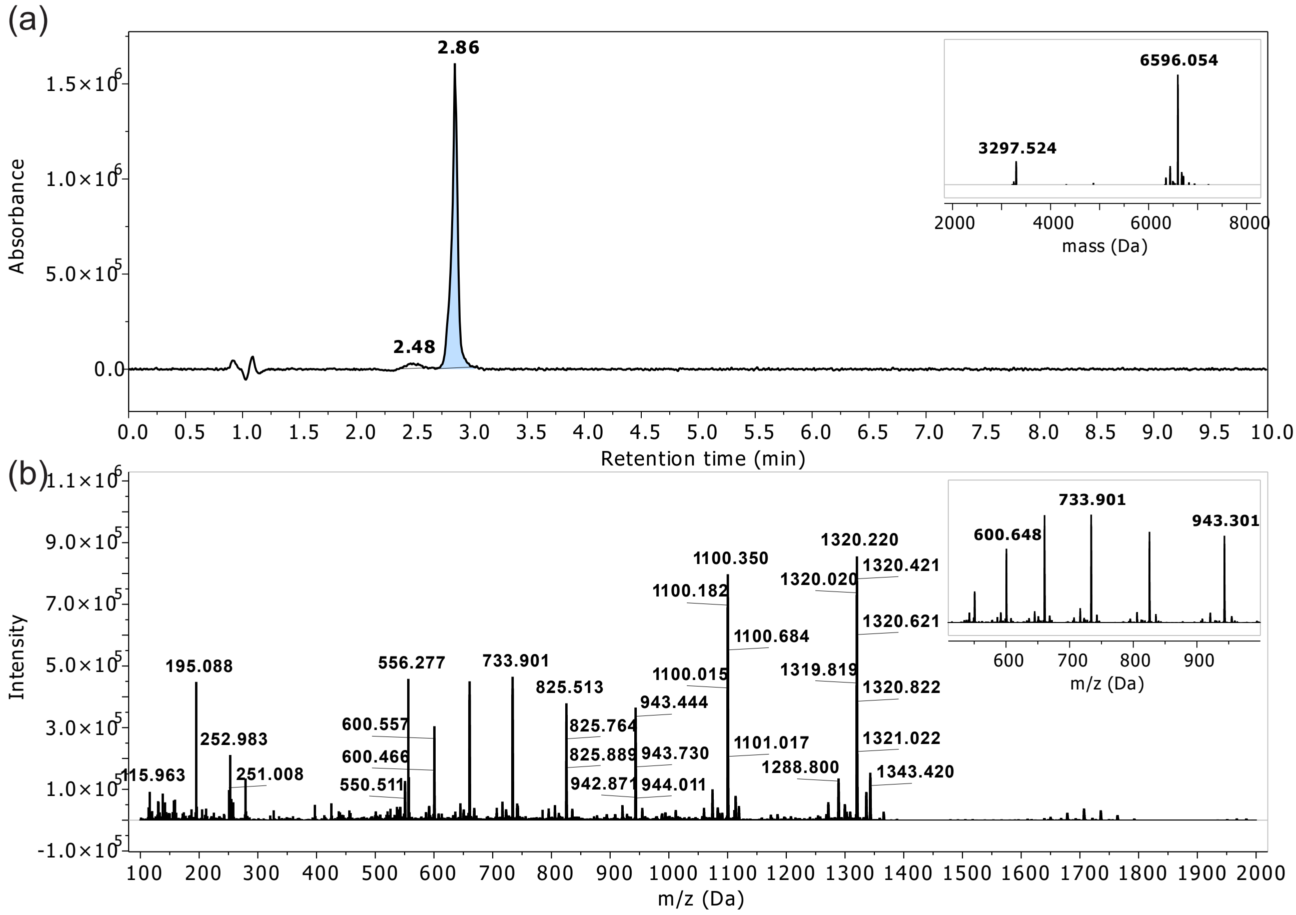


FIGURE S37. LCMS profile of pure RS-4×pS. (a) Absorbance chromatogram of pure RS-4×pS at $\lambda = 214$ nm with an Rt of 2.86 min (insert: deconvoluted masses). (b) ESI-TOF spectrum found within Rt 2 to 9 min (insert: convoluted spectra at Rt 2.86 min). Monoisotopic mass (ESI+) calculated for $C_{245}H_{426}N_{114}O_{90}P_4S_2$ is 6593.0654 Da; found 6593.0495 Da. LCMS Gradient A.

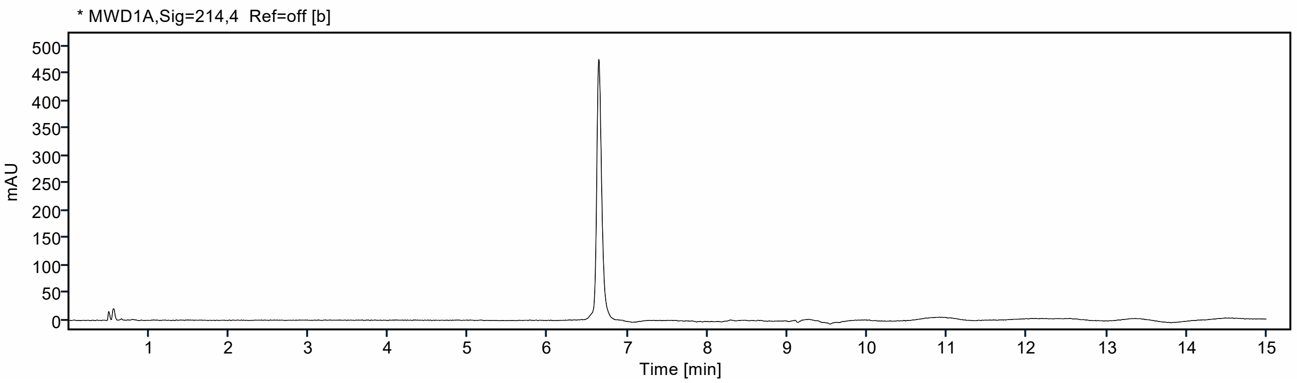


FIGURE S38. UHPLC profile of pure RS-4×pS using column C and a gradient of 0 to 100 % MeCN over 15 min at 40 °C. The peptide eluted at 6.64 min. Integration of the absorbance at $\lambda = 214$ nm between 1 to 15 min yields 99 % purity.
